# Cross-species analysis of ARPP19 phosphorylation during oocyte meiotic maturation charts the emergence of a new cAMP-dependent role in vertebrates

**DOI:** 10.1101/2023.07.05.547804

**Authors:** Ferdinand Meneau, Pascal Lapébie, Enrico Maria Daldello, Tran Le, Sandra Chevalier, Evelyn Houliston, Catherine Jessus, Marika Miot

## Abstract

In many animal species, elevated cAMP-PKA signaling initiates oocyte meiotic maturation upon hormonal stimulation, whereas in vertebrates, it acts as a negative regulator of this process. To address this “cAMP paradox”, we have focused on ARPP19 proteins. Dephosphorylation of *Xenopus* ARPP19 on a specific PKA site has been identified as a key step in initiating oocyte maturation. We first tracked evolution of the ARPP19 PKA phosphorylation site, revealing that it appeared early during the emergence of metazoans. This contrasts with strong conservation across eukaryotes of a phosphorylation site for the kinase Gwl in ARPP19 proteins, able to transform them into potent PP2A-B55 inhibitors and thus promote M-phase entry. We then compared the phosphorylation and function of *Xenopus* ARPP19 with its orthologue from the jellyfish *Clytia*, a model species showing cAMP-induced oocyte maturation. We confirmed that *Clytia* ARPP19 is phosphorylated on the conserved Gwl site *in vitro* as well as in maturing *Xenopus* and *Clytia* oocytes, behaving as a PP2A inhibitor and contributing to Cdk1 activation. However, Gwl-phosphorylated ARPP19 was unable to initiate oocyte maturation in *Clytia*, suggesting the presence of additional locks released by hormonal stimulation. *Clytia* ARPP19 was *in vitro* phosphorylated by PKA uniquely on the predicted site, but it was a much poorer substrate of PKA and of its antagonizing phosphatase, PP2A-B55δ, than the *Xenopus* protein. Correspondingly, PKA-phosphomimetic *Clytia* ARPP19 had a much weaker inhibitory activity on meiosis resumption in *Xenopus* oocytes than its *Xenopus* counterpart. Hence, poor recognition of *Clytia* ARPP19 by PKA and the absence of its targets in *Clytia* oocytes account for the cAMP paradox. This cross-species study of ARPP19 illustrates how initiation of oocyte maturation has complexified during animal evolution, and provides further insight into its biochemical regulation.

## INTRODUCTION

Meiosis in ovarian oocytes is arrested at the first prophase stage, until appropriate hormonal stimulation triggers progression into the meiotic divisions and the accompanying “maturation” events that prepare the egg for fertilisation. The molecular mechanisms responsible for maintaining and then releasing the prophase arrest of fully-grown ovarian oocytes have been the subject of intense study in several animal model species, but the regulation has proved very difficult to fully decipher and many questions remain open (Jessus *et al*, 2020). This reflects in part the convoluted evolutionary history of reproductive biology, in which the tissular origin and molecular nature of upstream regulating hormones has complexified independently in different animal lineages. As a consequence, the “Maturation Inducing Hormones” (MIHs) directly responsible for releasing the prophase arrest have varied molecular natures across species, and thus trigger distinct cytoplasmic signaling pathways (Quiroga Artigas *et al*, 2020; Von Stetina & Orr-Weaver, 2011; Voronina & Wessel, 2003). These diverse signaling pathways nevertheless all converge to activate a highly conserved kinase activation system that initiates entry into M-phase and thus the first meiotic division. The core element of this system is the Cdk1-Cyclin B complex known as MPF (M-phase promoting factor), which initiates M-phase by the phosphorylation of multiple substrates (Kishimoto, 2018; Masui, 2001). In this study, we take a comparative approach between two species that use opposing signaling pathways to trigger oocyte maturation initiation, in order to help unpick the hierarchy of regulations upstream of MPF activation. Specifically, we address the “cAMP paradox”, whereby cAMP/PKA signaling inhibits oocyte maturation initiation in vertebrates but triggers it in many other species, focusing on the sequence, phosphorylation characteristics and function of the regulatory protein ARPP19 from the amphibian *Xenopus* and its orthologue in the hydrozoan *Clytia*.

In vertebrates, high levels of intracellular cAMP and cAMP-dependent kinase (PKA) activity are essential in maintaining the prophase arrest. MIH stimulation induces a drop of both cAMP levels and of PKA activity to launch a signaling pathway leading to MPF activation, manifest as disassembly of the nuclear envelope, referred as GVBD for germinal vesicle breakdown (Maller *et al*, 1979; Ozon *et al*, 1979; Wang & Liu, 2004). cAMP levels and PKA activity can be experimentally manipulated by either inhibiting or stimulating the two antagonistic enzyme families that control cAMP level, adenylate cyclases and phosphodiesterases, or by overexpression or specific inhibition of PKA. Such experiments have revealed that in mammals, amphibia and fish oocytes, any elevation in cAMP or PKA activity blocks meiotic resumption induced by MIH, whereas cAMP decrease or PKA inhibition is sufficient to trigger meiotic resumption in the absence of MIH (Conti *et al*, 2002; Eyers *et al*, 2005; Huchon *et al*, 1981; Kovo *et al*, 2006; Maller & Krebs, 1977). Although the necessity of a drop in cAMP and PKA activity for meiosis resumption in *Xenopus* has been questioned (Nader *et al*, 2016), high levels of cAMP and PKA activity thus clearly play central roles in maintaining the prophase arrest in all vertebrates studied to date.

In various studied animal species from various non-vertebrate taxa, distinct signaling systems have been shown to be involved in maintaining or releasing the oocyte prophase arrest, some dependent on cAMP and PKA and some not. Remarkably, in several cases, cytoplasmic cAMP concentrations and PKA activity in the oocyte rise in response to MIH stimulation, and thus act as positive rather than negative regulators (Deguchi *et al*, 2011). Such cAMP rises during meiosis resumption have been documented in the brittle star *Amphipholis kochii* (echinoderm) (Yamashita, 1988), various nemerteans (Stricker & Smythe, 2001), the surf clam *Spisula solidissima* (mollusk) (Yi *et al*, 2002) and multiple hydrozoan species (Freeman & Ridgway, 1988; Takeda *et al*, 2006). Moreover, in these species, but also in *Boltenia villosa* (ascidian) (Lambert, 2011), *Pseudopotamilla occelata* (annelid) (Deguchi *et al*., 2011) and hydrozoans including *Cytaeis uchidae* and *Clytia hemisphaerica* (Amiel & Houliston, 2009; Takeda *et al*., 2006), meiosis resumption is triggered by externally-applied membrane-permeable cAMP, or cAMP injection, or by activators of adenylate cyclase or inhibitors of phosphodiesterase that increase cAMP. From these observations it is clear that in many animal species PKA is essential for triggering the pathway leading to MPF activation in response to MIH, and thus has a role opposite to that in vertebrates. The evolutionary history of this regulation is complex since in other cases including some echinoderms and ascidians, no role for cAMP has been detected in meiotic resumption, which is mediated by distinct signaling pathways (Deguchi *et al*., 2011).

The small protein ARPP19 (cAMP-regulated phosphoprotein 19), which in vertebrates has a paralog called ENSA, belongs to the Endosulfine protein family, widespread across eukaryotes (Labandera *et al*, 2015). In *Xenopus* oocytes, ARPP19 is phosphorylated by PKA on serine 109 (S109) and this S109-phosphorylated form is involved in maintaining the arrest in prophase (Dupre *et al*, 2014). In response to the *Xenopus* MIH, progesterone, ARPP19 becomes dephosphorylated on S109 by a specific phosphatase, PP2A-B55δ, authorizing the release of the prophase block (Labbe *et al*, 2021; Lemonnier *et al*, 2021). Subsequently, ARPP19 plays a second role in meiotic maturation, one widely shared with other dividing cells. It mediates inhibition of the PP2A-B55δ phosphatase, which is necessary for Cdk1 activation and thus entry into M-phase (Mochida *et al*, 2009; Vigneron *et al*, 2009). This inhibition is achieved by a distinct phosphorylation of ARPP19, by the protein kinase Greatwall (Gwl), on serine 67 (S67) (Dupre *et al*, 2013; Gharbi-Ayachi *et al*, 2010; Mochida *et al*, 2010). This specific phosphorylation converts ARPP19 into a potent and specific inhibitor of PP2A-B55δ and allows Cdk1 activation. ARPP19 thus assumes successively two distinct functions during *Xenopus* meiosis resumption. Its phosphorylation on S109 by PKA maintains the prophase arrest such that dephosphorylation on this site by PP2A-B55δ releases the prophase block. A poorly understood molecular signaling cascade is then induced leading to the phosphorylation of ARPP19 on S67 by Gwl. This inhibits PP2A-B55δ and activates Cdk1.

The positive role of ARPP19 on Cdk1 activation, through the S67 phosphorylation by Gwl, appears to be widespread across all eukaryotic mitotic and meiotic divisions (Dupre & Jessus, 2017). In contrast, the PKA-depending function of ARPP19 on the oocyte meiotic prophase arrest has only been studied so far in *Xenopus* (Dupre *et al*., 2013; Dupre *et al*., 2014; Dupre *et al*, 2017; Lemonnier *et al*., 2021). Here we investigate the phospho-regulation and role during oocyte maturation of ARPP19 from *Clytia hemisphaerica*, a laboratory model hydrozoan species well suited to oogenesis studies (Amiel *et al*, 2009; Jessus *et al*., 2020; Lechable *et al*, 2020; Munro *et al*, 2023; Quiroga Artigas *et al*, 2018). Meiotic maturation in hydrozoan oocytes is initiated by a rise in intracellular cAMP and PKA activity downstream of GPCR-Gα_s_ signaling (Freeman & Ridgway, 1988; Quiroga Artigas *et al*., 2020; Takeda *et al*., 2006). We first compared the sequence of ARPP19 proteins across eukaryotes to track its evolution, focusing specifically on the phosphorylation site by PKA. We then explored and compared the phosphorylation of *Clytia* and *Xenopus* ARPP19 proteins on the PKA-site and the Gwl-site. This functional comparative study revealed that, despite a recognisable and functional site of phosphorylation by PKA that emerged during animal evolution, *Clytia* ARPP19 is a poor substrate for PKA and PP2A-B55, the two enzymes regulating this phosphorylation site. Furthermore, the targets of ARPP19 regulating the resumption of meiosis in *Xenopus* are not functionally detectable in *Clytia* oocytes. These two features account for the ability of PKA activity to trigger meiosis resumption in *Clytia* oocyte despite the presence of ARPP19. More broadly, our results provide new perspectives regarding the intramolecular regulation of ARPP19 functions and the evolutionary scenarios underlying a universal feature of animal reproductive biology: maintenance and release of the oocyte prophase arrest.

## RESULTS

### The PKA phosphorylation site of ARPP19 is restricted to animals

To understand better the dual role of ARPP19 in regulating *Xenopus* oocyte meiotic maturation, we addressed the evolutionary conservation of the key sequence motifs phosphorylated by Gwl and PKA. Previous analyses focused on the site for phosphorylation by Gwl, which is highly conserved across eukaryotes suggesting an ancient origin and conserved function (Labandera *et al*., 2015). In this work, we examined more specifically the conservation of the PKA phosphorylation site, identified by functional analysis in various metazoans (Dulubova *et al*, 2001; Dupre *et al*., 2014; Horiuchi *et al*, 1990).

We aligned ARPP19 protein sequences recovered from publicly available genomic /transcriptome data from plants, fungi, choanoflagellates, teretosporeans and a range of metazoans (Fig. 1A, Supp Figs. S1 and S2). As highlighted in previous analysis (Labandera *et al*., 2015), the key motif for Gwl phosphorylation (FDS*G/AD) is very highly conserved across eukaryotes. The central serine of this motif is phosphorylated by Gwl to generate a potent PP2A-B55 inhibitor (Dupre *et al*., 2013; Gharbi-Ayachi *et al*., 2010; Juanes *et al*, 2013; Mochida *et al*., 2010). In contrast with the Gwl phosphorylation site, the known *Xenopus* ARPP19 PKA phosphorylation motif was only recognizable in the alignment in sequences from metazoan species (Labandera *et al*., 2015). It was not found in ARPP19 sequences from sponge species, but was clearly present in the ctenophore sequences. Under the scenario that ctenophores are the sister to all other animals (Schultz *et al*, 2023), this implies that it was present in a common metazoan ancestor. On the C-terminal side of the PKA site, we noted a SxL motif present in species scattered right across the metazoan phylogeny (including sponges), and also in the choanoflagellate *Hartaetosiga balthica* and the teretosporean *Sphaeroforma artica*. Even more widely conserved was a sequence lying N terminal side of the PKA phosphorylation motif, GxxxPTPxxϕP (where ϕ is a hydrophobic amino acid), detectable in the ARPP19 sequences across the opisthokonts (Fig. 1A, Supp Figs. S1 and S2). The significance of these motifs is unknown (see Discussion).

**Figure 1.**
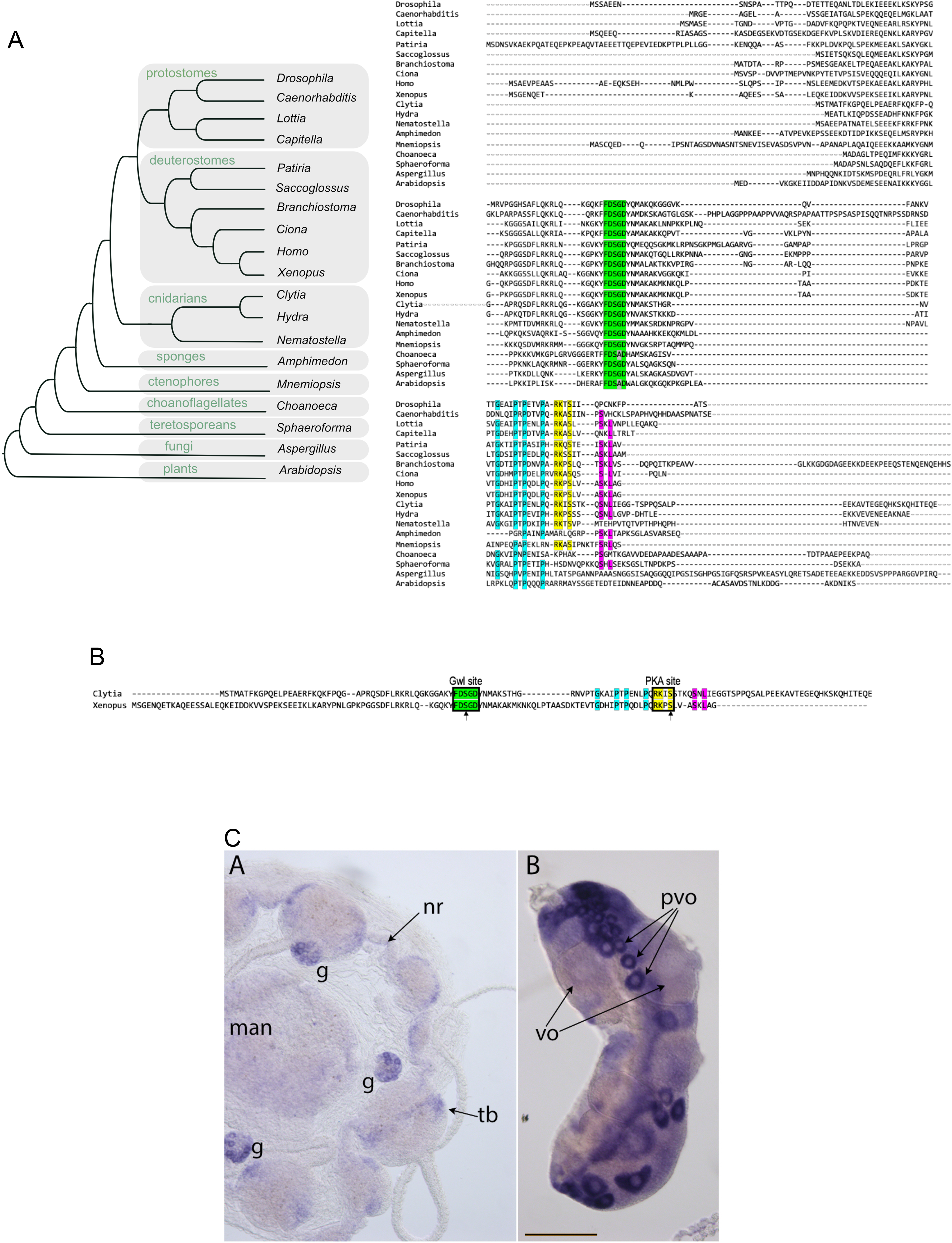
The conservation of the PKA site of ARPP19 is restricted to the animal kingdom. (A) Comparison of ARPP19 protein sequences from a selection of eukaryotic species (right). The Gwl (F-D-S*-G-D-Y) and PKA (R/K-R/K-X-S*/T*) phosphorylation sites, the GxxxPTPxxϕP sequence and the SxL terminal motif are colored in green, purple, yellow blue and red respectively. Residues in yellow and grey may vary from one species to another. The phylogenetic tree on the left indicates relationships between the main eukaryotic clades to which these species belong, adapted from(Grau-Bove *et al*, 2017). Additional sequence alignments and all accession numbers are provided in Supp Figs. S1 and S2. (B) Alignments of ClyARPP19 and XeARPP19 (same color code as in A). The Gwl (F-D-S*-G-D-Y) and PKA (R/K-R/K-X-S*/T*) phosphorylation sites identified in XeARPP19 are conserved in ClyARPP19. S67 and S109 correspond to the serines phosphorylated by Gwl and PKA respectively in XeARPP19. Their equivalents are respectively S49 and S81 in ClyARPP19. They are indicated by an arrow. (C) *In situ* hybridization detecting ClyARPP19 mRNA in a female baby medusa (left) and an adult isolated ovary (right). G= gonad; nr= nerve ring; tb= tentable bulb; man: manubrium (feeding organ); vo = vitellogenic oocytes; pvo= pre-vitellogenic oocytes. Scale bar= 100µm.

In conclusion, the site for PKA phosphorylation likely appeared as an important feature of ARPP19/ENSA proteins early during the emergence of metazoans. It remains very well conserved across the clade Eumetazoa (Cnidaria plus Bilateria), suggesting that phosphorylation on this site conveys an important biological property common to these animals. We thus undertook a comparative functional analysis of the PKA site of ARPP19 in the cnidarian *Clytia hemisphaerica* (ClyARPP19) and the amphibian *Xenopus laevis* (XeARPP19). The alignment of two sequences in Fig. 1B highlights the serines phosphorylated by Gwl and PKA in XeARPP19 (S67 and S109 respectively) and the corresponding serines in ClyARPP19 (S49 and S81 respectively). We focused on the process of oocyte maturation, in which intriguingly PKA plays opposite roles in the two species (Jessus *et al*., 2020). From available genome transcriptome data (Leclere *et al*, 2019; Takeda *et al*, 2018), we determined that *Clytia* has a single ARPP19 orthologue, expressed in growing and fully grown oocytes and across all life cycle stages. *In situ* hybridization experiments confirmed that ARPP19 mRNA is strongly expressed in ovarian oocytes and detected expression also in other tissues of the *Clytia* medusa, notably the nerve rings that run around the bell margin (Fig. 1C).

### The Cdk1 activation function of ARPP19 is conserved in *Clytia* and *Xenopus*

In *Xenopus* oocytes, ARPP19 phosphorylated by Gwl on S67 becomes a direct inhibitor of PP2A-B55δ, activating the Cdk1 auto-amplification loop and leading to meiotic division (Dupre *et al*., 2013). Injection into *Xenopus* prophase-arrested oocytes of XeARPP19 thiophosphorylated on S67 bypasses the progesterone-triggered maturation initiation mechanism to promote M-phase entry and meiotic divisions (Dupre *et al*., 2013) (Fig. 2A-B). To test whether ClyARPP19 can similarly induce meiosis resumption, we produced ClyARPP19 thiophosphorylated on S49 (Supp Fig. S3), injected it into *Xenopus* prophase oocytes, and then monitored GVBD as well as the phosphorylation level of MAPK and Cdk1, two molecular markers of meiotic cell division (Fig. 2A-B). Cdk1 is activated by Y15 dephosphorylation while MAPK is activated by phosphorylation (Jessus *et al*, 1991). Injection of S49-thiophosphorylated ClyARPP19 promoted meiosis resumption accompanied by Cdk1 and MAPK activation. ClyARPP19 phosphorylated on S49 is thus sufficient to promote meiotic maturation and Cdk1 activation in *Xenopus* oocyte, strongly suggesting that the *Clytia* protein shares with the *Xenopus* protein the inhibitory capacity of Gwl-phosphorylated ARPP19 towards PP2A-B55δ, leading to Cdk1 activation.

**Figure 2.**
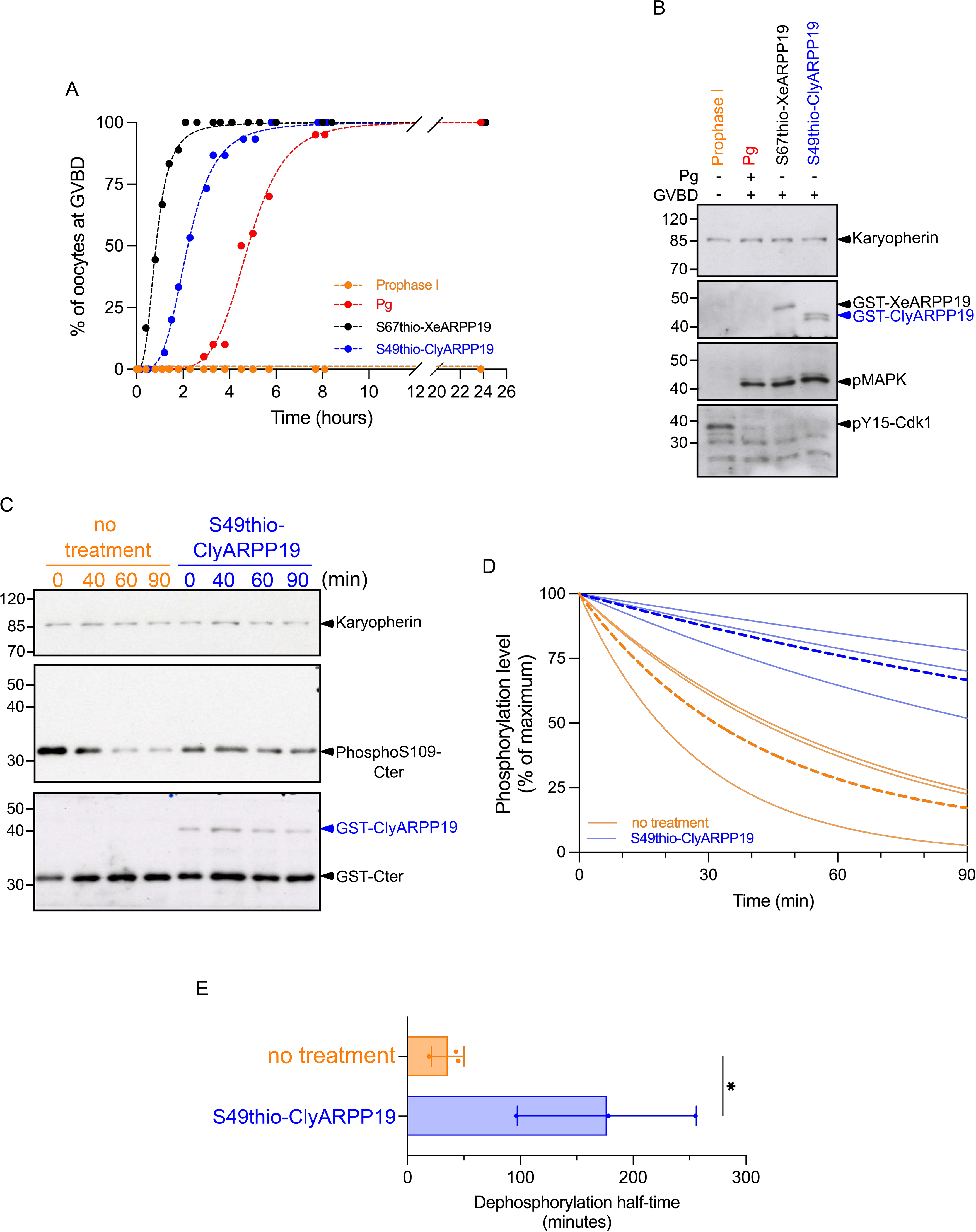
XeARPP19 and ClyARPP19 thio-phosphorylated on the Gwl site induce meiosis resumption in *Xenopus* oocytes and inhibit PP2A. **(A)** GVBD time-course of oocytes stimulated or not by progesterone (Pg) or injected with 200 ng of either GST-S67thio-XeARPP19 or GST-S49thio-ClyARPP19. One representative experiment out of 3 is shown. **(B)** Oocytes from the experiment illustrated in (A) were collected at GVBD and lysates were immunoblotted for karyopherin as a loading control, GST (tag of ARPP19 proteins), phosphorylated MAPK (pMAPK) and Y15-phosphorylated Cdk1 (pY15-Cdk1). The experiment was repeated 3 times with similar results. **(C)** *Xenopus* prophase oocyte extracts were incubated with hexokinase/glucose and PKI. Extracts were supplemented or not with S49-thiophosphorylated ClyARPP19 for 5 min, then with C-ter-XeARPP19 phosphorylated on S109 as a substrate. At indicated times, the phosphorylation of C-ter-XeARPP19 was analyzed by western blot with antibodies raised against phosphoS109-XeARPP19. The total amount of the substrate and of ClyARPP19 were detected by an anti-GST antibody. Karyopherin was used as a loading control. **(D)** Quantification of the phosphorylation level of C-ter-XeARPP19 of 3 independent experiments performed as in (C). The dashed lines correspond to the mean of the 3 experiments. **(E)** Comparison of the half-time dephosphorylation of C-ter-XeARPP19 calculated from 3 independent experiments performed as in (C), each represented by a dot. Paired T-test no treatment/S49thio-ClyARPP19 half-time Pvalue = 0.0388.

The inhibitory potential of S49-thiophosphorylated ClyARPP19 towards PP2A was confirmed in *Xenopus* prophase oocyte extracts. As already described (Lemonnier *et al*., 2021), PP2A is active in these extracts, efficiently dephosphorylates an artificial substrate, the C-ter part of XeARPP19 (amino-acids 68 to 117) phosphorylated on S109, and is inhibited by S67-thiophosphorylated XeARPP19. The *Xenopus* prophase oocyte extracts were supplemented with this phosphorylated C-ter substrate in the absence or presence of S49-thiophosphorylated ClyARPP19. PP2A activity was impaired by Gwl-phosphorylated ClyARPP19 (Fig. 2C-E), confirming the inhibitory activity of this protein towards PP2A.

### Gwl phosphorylation of ClyARPP19 is insufficient to initiate oocyte maturation

Although entry into first meiotic M-phase in both species is mediated by Cdk1 activation, the mechanisms that promote this activation following oocyte hormonal stimulation are molecularly distinct and far from fully understood (Jessus *et al*., 2020). In *Xenopus*, key progesterone-triggered events are a drop in cytoplasmic cAMP concentration, leading to synthesis of Cyclin B and Mos proteins, while in hydrozoans like *Clytia* binding of the peptide hormone MIH to its GPCR on the oocyte surface causes an immediate endogenous cytoplasmic cAMP rise, via Gα_s_ signaling (Deguchi *et al*., 2011; Quiroga Artigas *et al*., 2020; Takeda *et al*., 2018). Subsequently, meiotic maturation proceeds much more rapidly in *Clytia* than in *Xenopus*, GVBD occurring 10-20 minutes after treatment with MIH rather than several hours after progesterone stimulation. To gain insight into the molecular regulation of meiotic maturation in *Xenopus* vs *Clytia* oocytes, we investigated whether ARPP19 is phosphorylated on the Gwl site in *Clytia* oocytes during meiosis resumption. As endogenous ClyARPP19 is barely detectable by western blot, we injected exogenous ClyARPP19 and XeARPP19 into *Clytia* oocytes and analyzed the phosphorylation of the Gwl site in response to MIH stimulation. Both proteins were strongly phosphorylated on their Gwl sites (S49 and S67 respectively) in oocytes collected 15 minutes following MIH treatment (Fig. 3A-B). Thus, the mechanism of MPF activation in maturing *Clytia* oocytes matches that observed in *Xenopus* oocytes in involving phosphorylation of ARPP19 by Gwl or a related kinase downstream of MIH stimulation.

**Figure 3.**
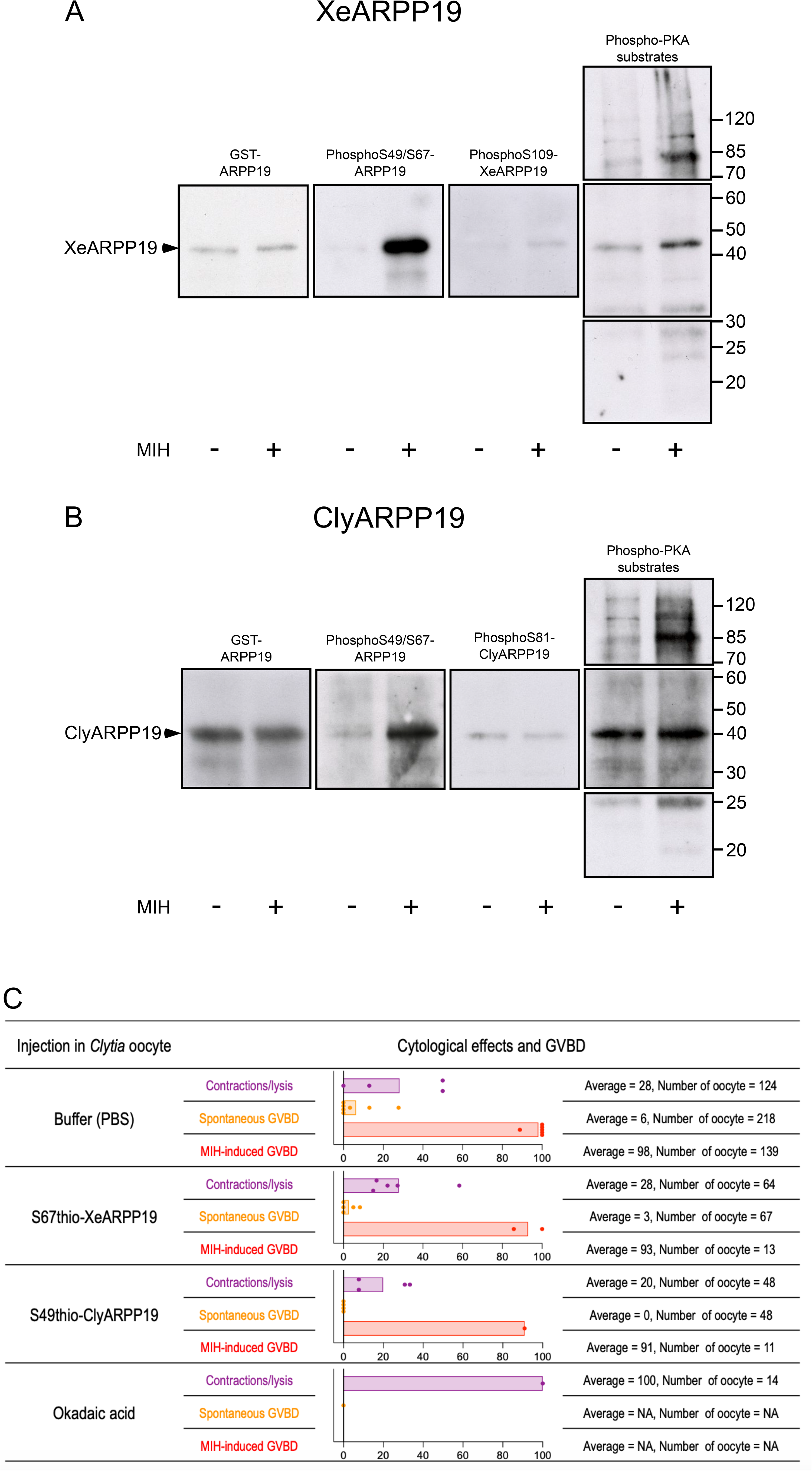
Phosphorylation and effects of XeARPP19 and ClyARPP19, or OA, on maturation in *Clytia* oocytes. **(A)** GST-XeARPP19 (3mg/ml) or control buffer were injected in *Clytia* prophase-arrested oocytes. One hour after injection, MIH was added or not. Oocytes were collected either at GVBD (15 minutes after MIH addition), or at the prophase stage (without MIH). Lysates were immunoblotted with antibodies against GST, phosphoS67-XeARPP19, phosphoS109-XeARPP19 and phosphoPKA substrates. **(B)** GST-ClyARPP19 (3mg/ml) or control buffer were injected in *Clytia* prophase-arrested oocytes. One hour after injection, MIH was added or not. Oocytes were collected either at GVBD (15 minutes after MIH addition), or at the prophase stage (without MIH). Lysates were immunoblotted with antibodies against GST, phosphoS67-XeARPP19, phosphoS81-ClyARPP19 and phosphoPKA substrates. **(C)** Summary of experiments in which GST-XeARPP19 (4mg/ml), GST-ClyARPP19 (4mg/ml), OA (2μM) or control buffer were injected into *Clytia* prophase-arrested oocytes. Within one hour after injection, contractions followed by the lysis of the oocytes were observed in some cases (purple). The percentage of oocytes exhibiting this cytological effect is indicated. In other experiments, MIH (10^-7^ M WPRP-amide) was added and GVBD was monitored. The percentage of oocytes that underwent GVBD in the absence (spontaneous GVBD, orange) or in the presence (MIH-induced GVBD, red) of MIH is indicated. Each point represents one experiment. The average of the experiments (A) is indicated for each condition. Full details and data for these experiments is provided in Supplementary Table 1.

We further tested whether injection of Gwl-phosphorylated ARPP19 proteins could promote oocyte meiotic maturation in *Clytia*. We were unable to induce maturation of *Clytia* oocytes by injection of any versions of *Clytia* or *Xenopus* ARPP19 including wild-type XeARPP19 or ClyARPP19 thiophosphorylated by Gwl. Injecting 4mg/ml pipette concentration of these two proteins induced only non-specific incidence of GVBD (0 or 1 oocyte from each of 4 groups of 9-20 oocytes injected with each of the thiophosphorylated proteins, comparable with buffer injections; results summarized in Fig. 3C, full details in Supplementary Table 1). It was not possible to increase injection volumes or protein concentrations without inducing high levels of non-specific toxicity. These experiments suggest that, unlike in *Xenopus*, PP2A inhibition is not sufficient to induce oocyte meiotic division in *Clytia*. To confirm that this lack of maturation induction was not due to ineffective PP2A inhibition by the injected proteins, we repeated the experiment injecting okadaic acid (OA), a powerful PP2A inhibitor (Takai *et al*, 1987) that induces meiosis resumption in *Xenopus* oocytes (Goris *et al*, 1989). Injection of *Clytia* oocytes with OA at 2µM did not promote GVBD, however between 40 and 90 minutes post-injection all injected oocytes showed cortical contractions followed by fragmentation and/or degradation (Fig. 3C), suggesting that prolonged PP2A inhibition is lethal to the oocytes. These results suggest that PP2A inhibition is not sufficient to induce oocyte maturation in *Clytia*, although we cannot rule out that the quantity of OA or Gwl-thiophosphorylated ARPP proteins delivered was insufficient to trigger GVBD.

To summarise these findings, we have shown that ClyARPP19 is phosphorylated on the Gwl site in response to MIH and that Gwl-phosphorylated ARPP19 from either *Clytia* or *Xenopus* origin can inhibit PP2A, confirming the strong conservation of this function in eukaryotes. In *Clytia* oocytes, however, unlike in *Xenopus* oocytes, this inhibition appears insufficient to override the prophase arrest mechanism

### ClyARPP19 is phosphorylated on S81 by PKA *in vitro*

In *Xenopus*, phosphorylation of ARPP19 by PKA on S109 within the PKA consensus motif RKPS_109_L maintains the oocyte prophase arrest (Dupre *et al*., 2014). The general consensus PKA site is reported to be R/K-R/K-X-S*/T*. The phosphopredict program retrieved the site RKIS_81_S_82_ in ClyARPP19 recognized in our alignments (Fig1. A) as a PKA consensus sequence (Fig. 1B). It includes two serines, S81 and S82, the first matching the position of the phosphorylated residue in the general PKA consensus site. To determine if one or both of these are phosphorylated by PKA, we produced four forms of ClyARPP19 tagged by GST: wild-type (WT), the single amino acid substitutions S81D and S82D, and the double mutant S81D-S82D (Fig. 4A), the ‘phosphomimetic’ substitution of Serine with Aspartic acid preventing phosphorylation. All recombinant proteins were incubated *in vitro* in the presence of purified recombinant bovine PKA and γS-ATP. An extended period of incubation (2 hours) was required to detect their phosphorylation by western blot with an antibody that detects thiophosphorylated residues (Fig. 4B, Supp Fig. S4). Wild-type ClyARPP19 but not the double S81D-S82D was phosphorylated by PKA *in vitro*, showing that the only residues phosphorylated by PKA in ClyARPP19 are located in the PKA consensus motif. As predicted, no phosphorylation was detected in the S81D mutant. The S82D mutant was phosphorylated by PKA, although the level of phosphorylation was reduced compared to that of WT-ClyARPP19 (Fig. 4B, Supp Fig. S4), indicating that mutation of this residue impairs the phosphorylation of the neighboring S81. This *in vitro* assay thus places S81 as the sole residue in ClyARPP19 for phosphorylation by PKA.

**Figure 4.**
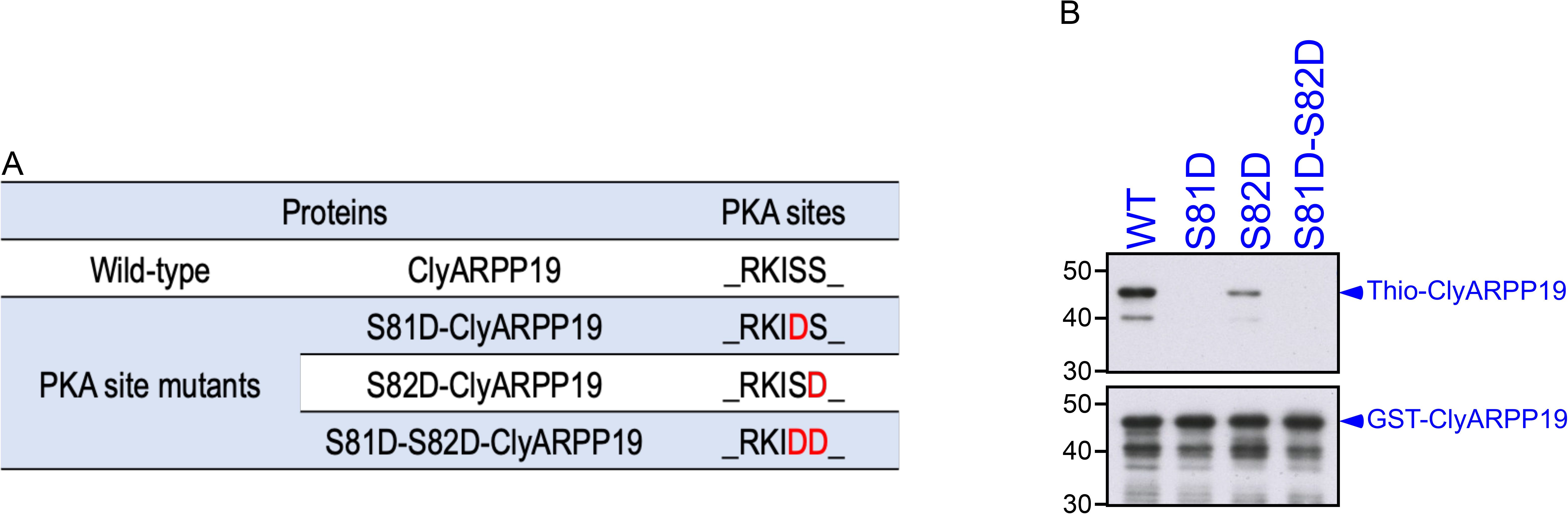
Identification of the ClyARPP19 residue phosphorylated by PKA. **(A)** Summary of the different mutations introduced into ClyARPP19. **(B)** Wild-type (WT), single or double mutants of ClyARPP19 at S81 and S82 (S81D, S82D and S81D-S82D) were thiophosphorylated *in vitro* using recombinant bovine PKA catalytic subunit in the presence of γS-ATP for 2 hours. The thiophosphorylation was analyzed by western-blot with an anti-thiophosphate ester antibody. The levels of ClyARPP19 proteins were detected with an anti-GST antibody. The experiment was repeated 3 times with similar results.

### ClyARPP19 only weakly affects *Xenopus* oocyte meiosis resumption

When injected into prophase-arrested *Xenopus* oocytes, XeARPP19 rapidly becomes phosphorylated by PKA on S109, and this exogenous protein prevents progesterone from inducing GVBD in a dose-dependent manner (Dupre *et al*., 2014). To test whether ClyARPP19 has equivalent inhibitory properties, we injected it into *Xenopus* oocytes and monitored GVBD following progesterone stimulation. Whereas injection of XeARPP19 delayed meiosis resumption (2-fold increase in GVBD_50_) and fully inhibited GVBD in 35% of oocytes (Fig. 5A-C, Supp Fig. S5), injection of the same amount (800 ng) of ClyARPP19 had no effect on the timing of GVBD, with 100% of oocytes injected with ClyARPP19 showing a similar time-course to uninjected oocytes (Fig. 5A-C, Supp Fig. S5).

**Figure 5.**
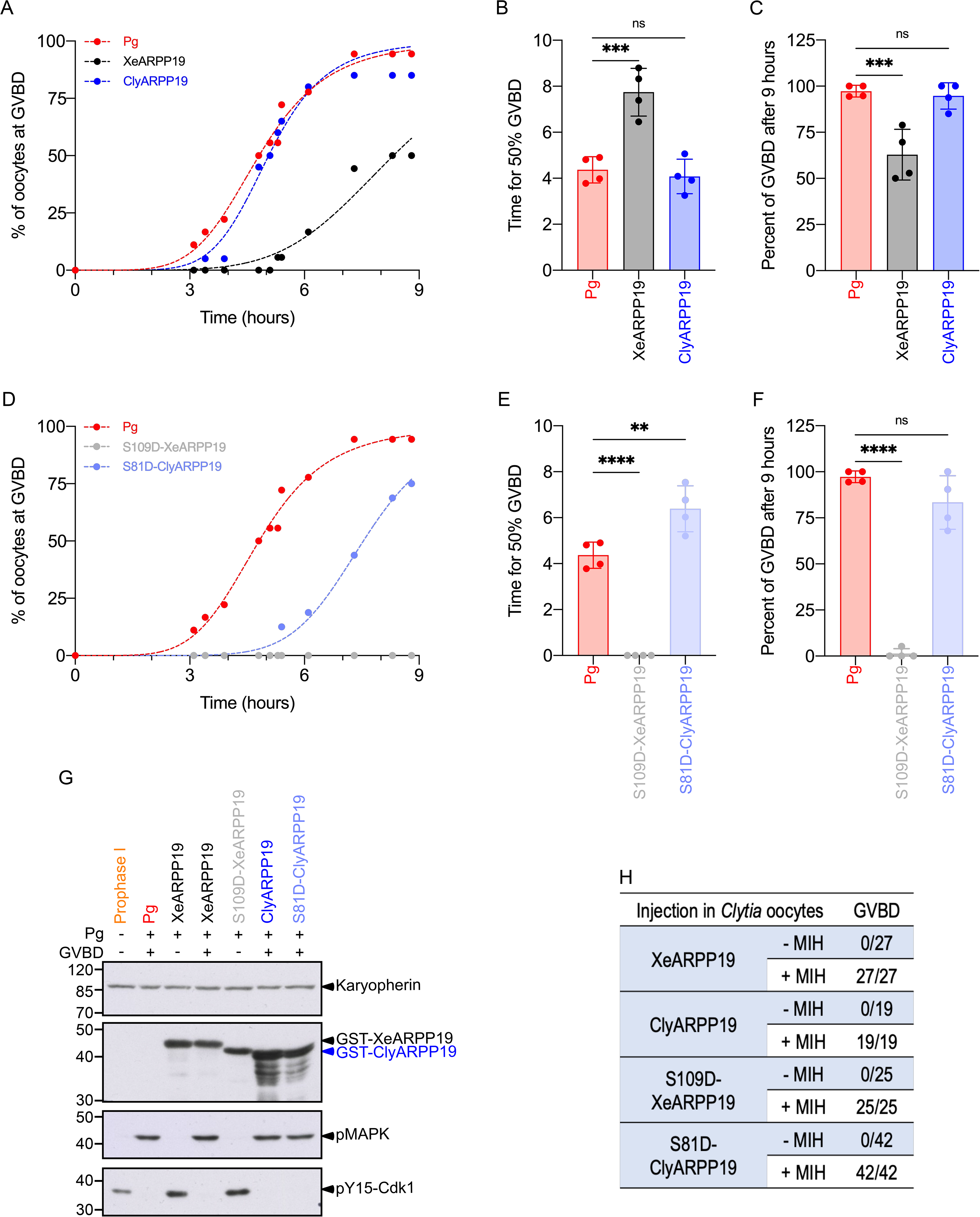
ClyARPP19 and XeARPP19 phosphorylated on the PKA sites modulate oocyte meiosis resumption. **(A)** GVBD time course of oocytes injected with either XeARPP19 or ClyARPP19 (800 ng/oocyte) and then stimulated by Pg. One representative experiment from 4 independent experiments is shown. **(B)** Time to reach 50% GVBD for the 3 conditions of panel (A). Ordinary one-way ANOVA analysis: XeARPP19/Pg Pvalue=0.0005, ClyARPP19/Pg Pvalue=0.8593 (ns). Each dot represents an independent experiment. **(C)** % of GVBD 9 hours after Pg addition for the 3 conditions of panel (A). Ordinary one-way ANOVA: XeARPP19/Pg Pvalue=0.0009, ClyARPP19/Pg Pvalue=0.9093 (ns). Each dot represents an independent experiment. **(D)** Same experiment as in (A) with injection of S109D-XeARPP19 or S81D-ClyARPP19 (800 ng/oocyte). One representative experiment from 4 independent experiments is shown. **(E)** Time to reach 50% GVBD for the 3 conditions of panel (D). Note that S109D-XeARPP19 injected oocytes never reached GVBD. Ordinary one-way ANOVA: Pg/S81D-ClyARPP19 Pvalue=0.0040. Each dot represents an independent experiment. **(F)** % of GVBD 9 hours after Pg addition for the 3 conditions of panel (D). Ordinary one-way ANOVA: Pg/S109D-XeARPP19 Pvalue<0.0001, Pg/S81D-ClyARPP19 Pvalue=0.0988. Each dot represents an independent experiment. **(G)** Oocytes from experiments represented in (A) and (D) were collected at GVBD and lysates were immunoblotted for karyopherin as a loading control, GST (tag of ARPP19 proteins), phosphorylated MAPK (pMAPK) and Y15-phosphorylated Cdk1 (pY15-Cdk1). The experiment was repeated 4 times with similar results. **(H)** ARPP19 proteins, either wild-type or mutated (S81D-ClyARPP19 or S109D-XeARPP19) were injected in *Clytia* prophase-arrested oocytes. MIH was then added or not and GVBD was monitored. The number of oocytes having undergone GVBD is indicated.

How PKA-phosphorylated XeARPP19 inhibits meiosis resumption in *Xenopus* is not yet understood, but must involve unidentified cis or trans interactions specific to this form that somehow prevent initiation of the Cdk1 and MAPK activation cascades (Dupre & Jessus, 2017; Dupre *et al*., 2017; Labbe *et al*., 2021). We reasoned that the lack of inhibitory activity of ClyARPP19 in the *Xenopus* oocyte could reflect its inability to recognise targets of the *Xenopus* protein and/or less efficient phosphorylation by PKA. To address the first hypothesis, we took advantage of phosphomimetic mutant proteins. The S109D-XeARPP19 mutant mimics constitutive phosphorylation on S109 and inhibits meiotic resumption (Dupre *et al*., 2014). We produced ClyARPP19 with the equivalent mutation, S81D, and compared the effects of progesterone following injection into *Xenopus* oocytes (Fig. 5D-F, Supp Fig. S5). While S109D-XeARPP19 fully abolished Pg-induced GVBD, S81D-ClyARPP19 did not, although the time course of meiosis resumption was delayed (1.5-fold increase in GVBD_50_ compared to uninjected oocytes or oocytes injected with wild-type ClyARPP19) (Fig. 5D-F, Supp Fig. S5). In all experiments, the observed effects were confirmed molecularly by monitoring the phosphorylation of MAPK and Cdk1 (Fig. 5G). Hence, ClyARPP19, when phosphorylated on S81, has a weaker inhibitory activity on meiosis resumption than its *Xenopus* counterpart, suggesting that it cannot effectively recognise XeARPP19 targets.

This lack of recognition could have two origins: either equivalent ARPP19 interactors are present in *Clytia* oocytes but their *Xenopus* counterparts are too divergent in sequence to be recognized by ClyARPP19, or they are absent in *Clytia* oocytes. To address this question, we injected the S81D phospho-mimetic mutant protein into *Clytia* oocytes, which were then stimulated with MIH. Neither this mutant protein nor wild-type ClyARPP19 affected meiotic resumption, as seen by GVBD in all cases (Fig. 5H). Furthermore, neither S109D-XeARPP19 nor wild-type XeARPP19 affected meiotic resumption in *Clytia* oocytes (Fig. 5H). Although we cannot rule out that the amount of protein injected was insufficient to compete with endogenous pools, these results support the hypothesis that specific interactors of PKA-phosphorylated ARPP19 that prevent entry of *Xenopus* oocytes into first meiotic M-phase are absent from *Clytia* oocytes. They do not exclude the possibility that ClyARPP19 is phosphorylated only weakly or not at all by PKA i*n vivo* in *Clytia* oocytes.

### Inefficient S81 phosphorylation of ClyARPP19 in *Xenopus* oocyte extracts

We addressed the efficiency of S81 phosphorylation by PKA of ClyARPP19 compared to S109 of XeARPP19 using extracts from prophase-arrested *Xenopus* oocytes, in which PKA is active. Purified GST-tagged ClyARPP19 or XeARPP19 proteins were incubated in extracts pre-incubated or not with PKI, a specific inhibitor of PKA. S81 phosphorylation of ClyARPP19 and S109 phosphorylation of XeARPP19 were monitored by western blot using anti-phospho-antibodies that specifically recognize phospho-S81-ClyARPP19 and phospho-S109-XeARPP19 (Fig. 6A, Supp Fig. S6). S109 phosphorylation of XeARPP19 was detected after only 5 minutes of incubation and then increased during the following 15 minutes (Fig. 6A). This phosphorylation was totally abolished in the presence of PKI (Fig. 6A), confirming that XeARPP19 is efficiently phosphorylated by PKA at the prophase stage (Dupre *et al*., 2014). In contrast, S81 phosphorylation of ClyARPP19 was not detected by the anti-phospho-S81 antibody in either condition (Fig. 6A; compare with the positive control of ClyARPP19 *in vitro* phosphorylated by PKA after a 2 hour-incubation). We conclude that in *Xenopus* prophase oocyte extracts, ClyARPP19 is not detectably phosphorylated on S81, whereas XeARPP19 shows phosphorylation on S109 strictly dependent of PKA.

**Figure 6.**
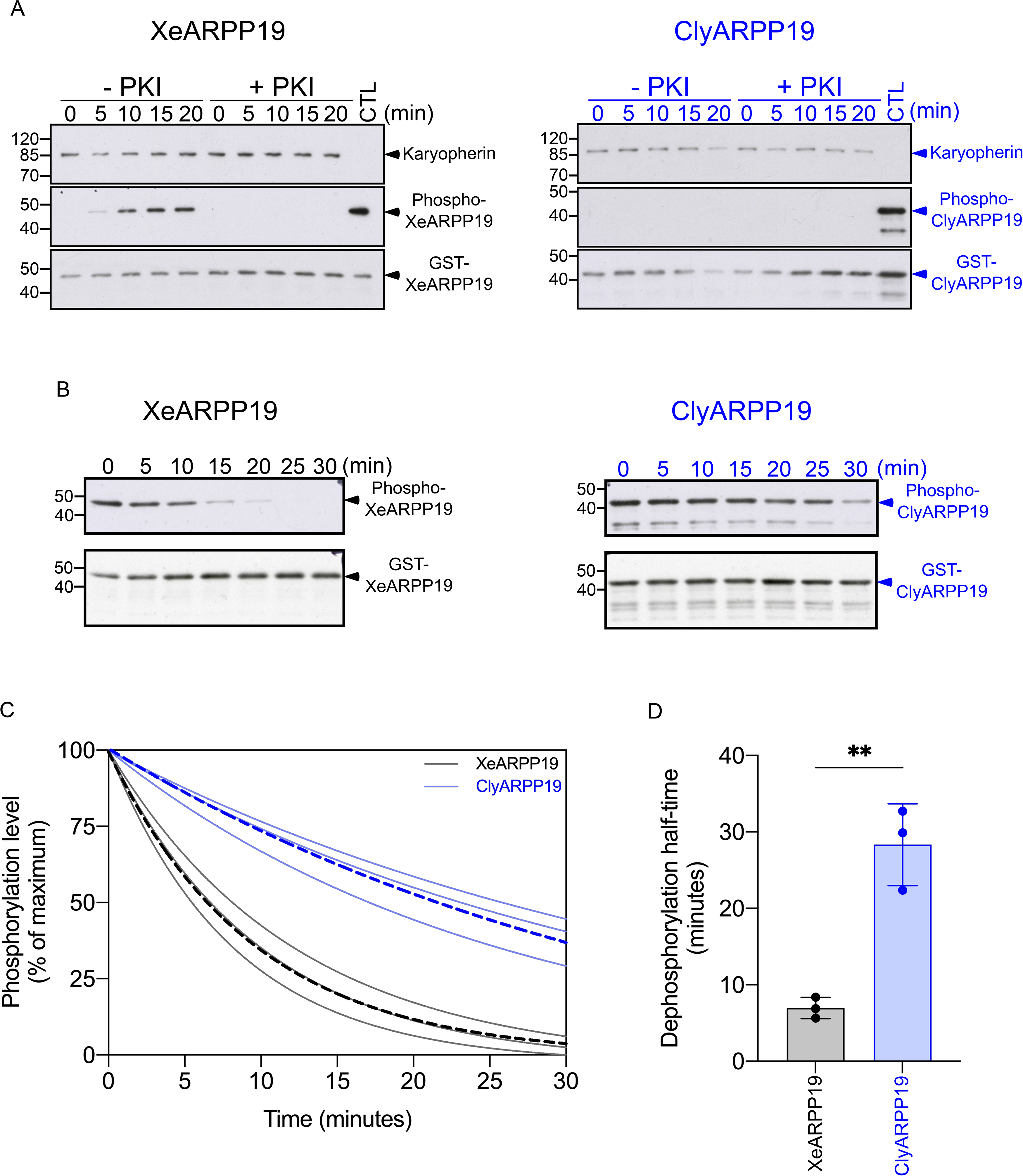
S81 phosphorylation and dephosphorylation of ClyARPP19 in *Xenopus* oocyte extracts. **(A)** After 30 minutes preincubation with or without PKI, *Xenopus* prophase oocyte extracts were supplemented with XeARPP19 (left panel) or ClyARPP19 (right panel). At indicated times, samples were collected and phosphorylation was analyzed by western blot with antibodies raised against phosphoS109-XeARPP19 or phosphoS81-ClyARPP19. The total ARPP19 amount was detected by an anti-GST antibody. Karyopherin was used as a loading control. A control (CTL) is represented by recombinant XeARPP19 and ClyARPP19 *in vitro* phosphorylated by PKA for 2 hours. One representative experiment from 3 independent experiments is shown. **(B)** *Xenopus* prophase oocyte extracts were incubated with hexokinase/glucose and PKI. They were supplemented with S109-phosphorylated XeARPP19 (left panel) or S81-phosphorylated ClyARPP19 (right panel). At indicated times, samples were collected and the phosphorylation level was followed by western blot using antibodies directed against phosphoS109-XeARPP19 or phosphoS81-ClyARPP19. The total ARPP19 amount was detected by an anti-GST antibody. One representative experiment from 3 independent experiments is shown. **(C)** Quantification of the phosphorylation level of 3 independent experiments performed as in (B). The dashed lines correspond to the mean of the 3 experiments. **(D)** Comparison of the half-time dephosphorylation of phosphoS81-ClyARPP19 and phosphoS109-XeARPP19 calculated from 3 independent experiments performed as in (B), each represented by a dot. Unpaired T-test XeARPP19/ClyARPP19 half-time Pvalue = 0.0026.

Two possible mechanisms can account for the absence of ClyARPP19 detectable phosphorylation on S81 in *Xenopus* oocyte extracts. Either S81 of ClyARPP19 is rapidly dephosphorylated by a phosphatase and/or the ability of PKA to phosphorylate ClyARPP19 is lower than for the *Xenopus* counterpart as already observed in the assays using purified PKA (see above). To address the first mechanism, we measured the dephosphorylation rate of ARPP19 proteins. We produced ClyARPP19 and XeARPP19 phosphorylated on S81 and S109 respectively by a 4 hours *in vitro* incubation with PKA. In order to compare the dephosphorylation of equivalent starting amounts of phosphorylated ClyARPP19 and XeARPP19, their phosphorylation levels were calibrated as shown in Supp Fig. S7. The prophase oocyte extracts were supplemented with PKI to inhibit PKA activity and with hexokinase/glucose to deplete ATP and thus eliminate any kinase activity. S81-phosphorylated-ClyARPP19 or S109-phosphorylated-XeARPP19 were added to these kinase-dead oocyte extracts, and aliquots were collected over time to estimate ARPP19 dephosphorylation using specific phospho-antibodies. As shown in Fig. 6B-D, 50% dephosphorylation was reached in 7 minutes for XeARPP19 and 28 minutes for ClyARPP19, meaning that XeARPP19 was dephosphorylated 4 times faster than ClyArpp19. Hence, for the same starting amount of phosphorylated proteins and in the absence of any kinase activity, the *Clytia* protein is dephosphorylated less efficiently than its *Xenopus* counterpart (Fig. 6A-D and Supp Fig. S8), indicating that ClyARPP19 is a poor substrate of the phosphatase specific of the S81 residue.

It has been shown that in *Xenopus,* ARPP19 is dephosphorylated on S109 by PP2A-B55δ (Lemonnier *et al*., 2021). To verify that this PP2A isoform also dephosphorylates ClyARPP19, the dephosphorylation assay was repeated in prophase oocyte extracts depleted in PP2A-B55δ. Extracts were incubated with beads coupled to both a specific anti-B55δ antibody (Supp Fig. S9) and the S67-thio-phosphorylated form of XeARPP19 (thio-S67-ARPP19), which acts as a specific inhibitor of the PP2A-B55δ isoform (Gharbi-Ayachi *et al*., 2010; Mochida *et al*., 2010). To avoid any side-effects from the S109 residue of the thio-S67-ARPP19 inhibitor, we produced a S109A mutant, called thio-S67-S109A-ARPP19. We assessed that the level of B55δ was reduced in the extracts after removal of the beads (Fig. 7A). Dephosphorylation of ClyARPP19 and XeARPP19 was estimated after a 45 min-incubation in these depleted extracts. As shown in Fig. 7, XeARPP19 and ClyARPP19 dephosphorylation on S109 and S81 respectively were strongly reduced.

**Figure 7.**
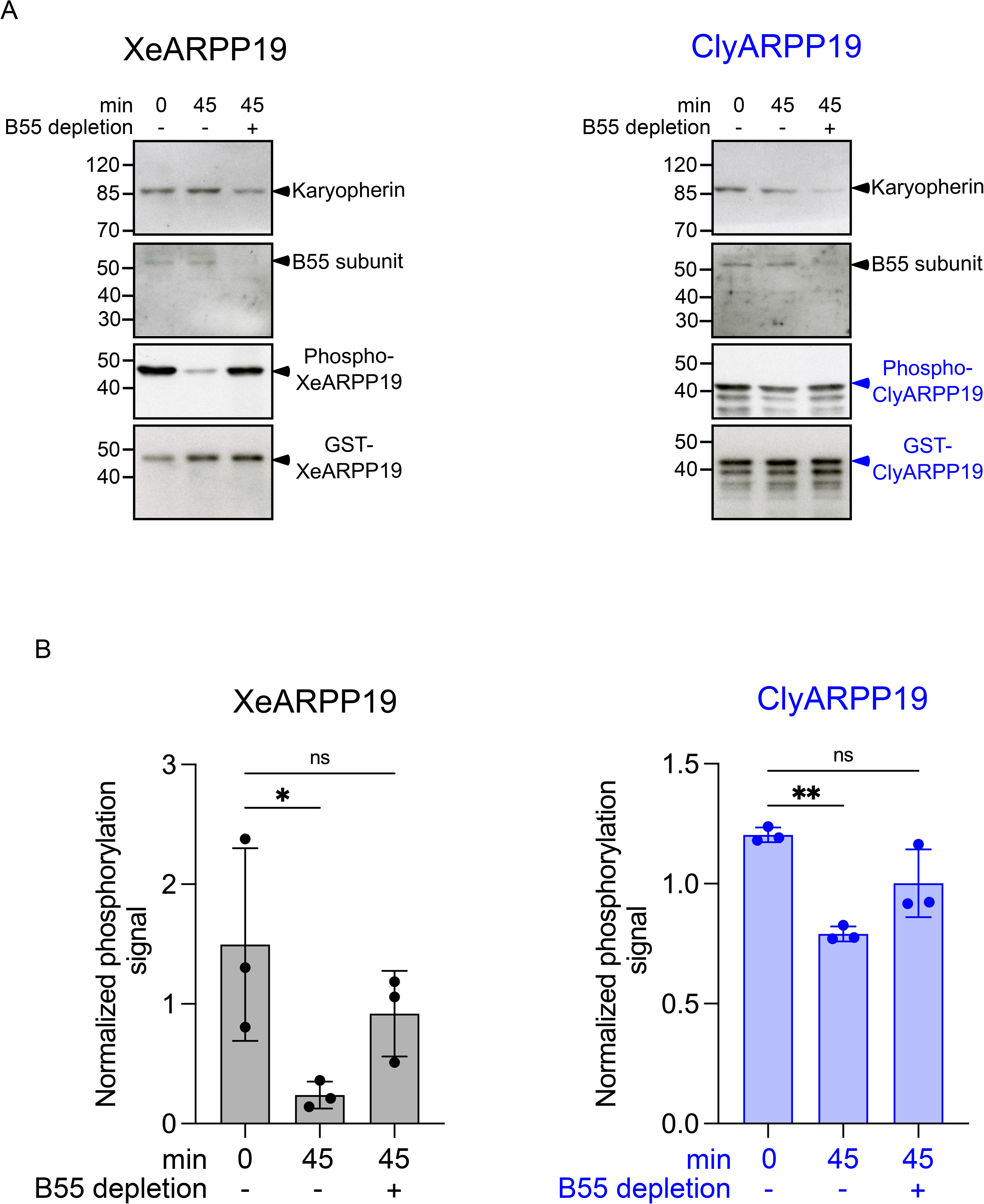
ClyARPP19 is dephosphorylated by PP2A-B55δ on S81. **(A)** *Xenopus* prophase oocyte extracts were incubated with hexokinase/glucose and PKI. They were depleted or not in B55δ by beads coated with S67thio-S109A-XeARPP19 and B55δ antibody. They were then supplemented with S109-phosphorylated XeARPP19 or S81-phosphorylated ClyARPP19 for 45 min. The phosphorylation level was estimated by western blot using antibodies directed against phosphoS109-XeARPP19 or phosphoS81-ClyARPP19. The total Arpp19 amount was detected by an anti-GST antibody and B55δ by a specific antibody. Karyopherin was used as a loading control. One representative experiment from 3 independent experiments is shown. **(B)** Quantification of the phosphorylation level at 45 min of 3 independent experiments performed as in (A), each of them represented by a dot. Data are shown as mean +/- SD. Ordinary one-way ANOVA has been applied. S109XeARPP19 dephosphorylation: no treatment 0’/no treatment 45’ Pvalue=0.0471, no treatment 0’/B55 depletion Pvalue=0.3873. S81ClyARPP19 dephosphorylation: no treatment 0’/no treatment 45’ Pvalue=0.0021, no treatment 0’/B55 depletion Pvalue=0.0537.

Taken together these results show that ClyARPP19 is dephosphorylated on S81 by PP2A-B55δ much less efficiently than the *Xenopus* protein. This implies that the absence of S81 phosphorylation in the prophase oocyte extract (Fig. 6A) is not due to hyper active dephosphorylation but rather to poor phosphorylation by PKA.

### ClyARPP19 is poorly phosphorylated by PKA *in vitro* and *in vivo*

We compared PKA phosphorylation of ClyARPP19 versus XeARPP19 first *in vitro* using recombinant bovine PKA, and then *in vivo* by injection into *Xenopus* and *Clytia* oocytes. To assess the *in vitro* phosphorylation rate of purified ClyARPP19 and XeARPP19, the proteins were thio-phosphorylated by using γS-ATP, enabling us to make direct comparisons using a single antibody targeting thio-phosphates rather than using separate species-specific phospho-antibodies. ClyARPP19 or XeARPP19 were incubated with recombinant PKA and γS-ATP and samples were collected at successive times up to 60 min (Fig. 8A). Thio-phosphorylation of XeARPP19 was detected after 5 min of incubation with PKA and increased until 60 min. In contrast, ClyARPP19 thiophosphorylation only started to be detected at 30 min. Quantification of 3 independent experiments determined that ClyARPP19 is indeed phosphorylated by PKA less efficiently than XeARPP19 (Fig. 8B-C, Supp Fig. S10). 50% of XeARPP19 was phosphorylated within 18 minutes whereas the extrapolation of the curve indicates that it would take 148 minutes to reach the same level for ClyARPP19, showing that thio-phosphorylation of XeARPP19 is 8 times faster than that of ClyARPP19 (Fig. 8C). ClyARPP19 is thus a relatively poor substrate of purified PKA.

**Figure 8.**
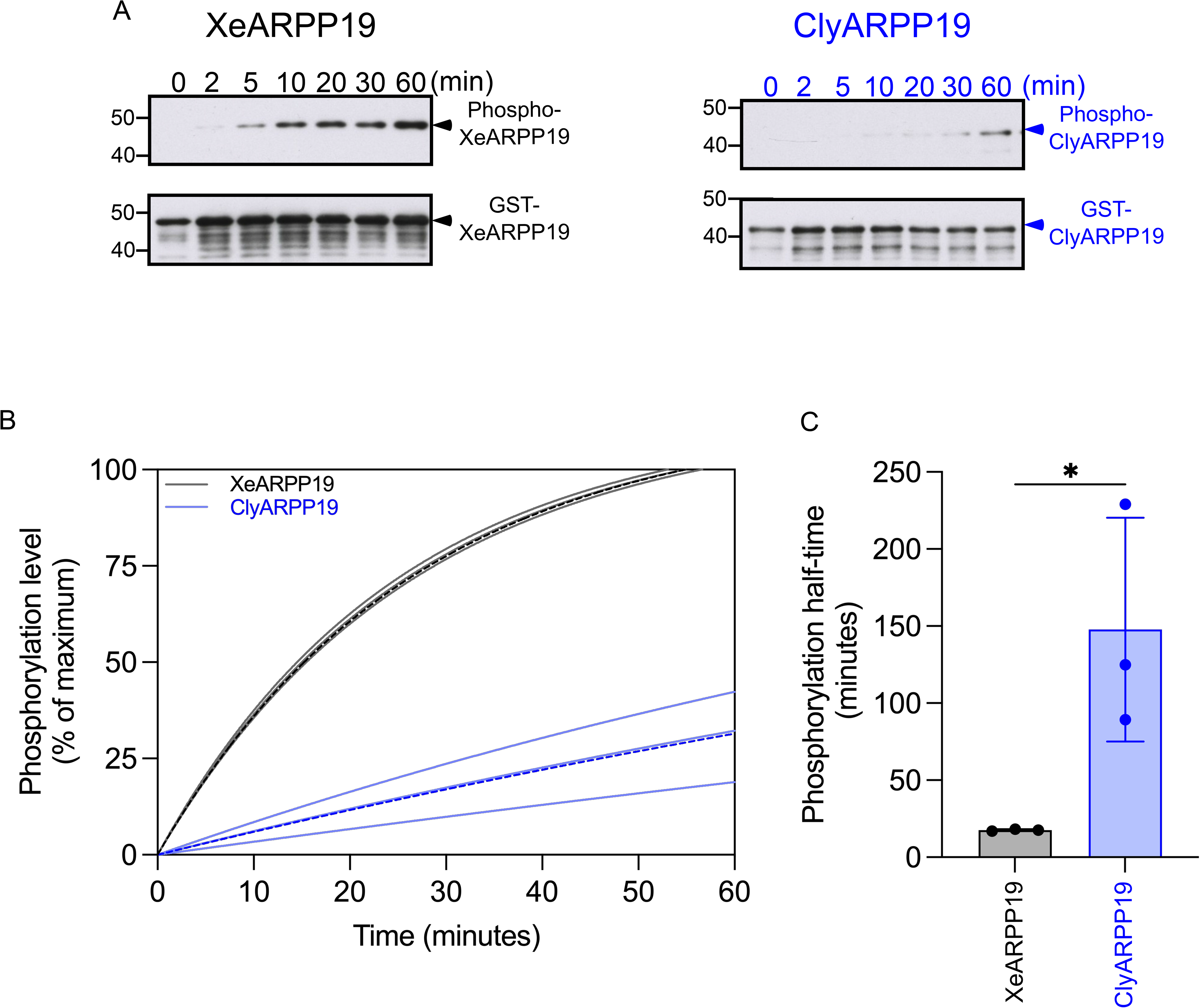
ClyARPP19 is phosphorylated *in vitro* by PKA at low rate compared to XeARPP19. **(A)** Purified bovine PKA was incubated with XeARPP19 or ClyARPP19 in the presence of γS-ATP. Samples were collected at various times as indicated. The phosphorylation of ARPP19 proteins was assessed by western blot with an anti-thiophosphate ester antibody. The total ARPP19 amount was detected by an anti-GST antibody. **(B)** Quantification of the phosphorylation level of 3 independent experiments performed as in (A). The dashed lines correspond to the mean of the 3 experiments. **(C)** The half-time of XeARPP19 and ClyARPP19 phosphorylation by PKA was determined from the regression curve of the phosphorylation level obtained for the 3 independent experiments, each represented by a dot. Unpaired T-test XeARPP19/ClyARPP19 half-time Pvalue=0.0361.

To assess phosphorylation of ARPP19 proteins in intact oocytes, ClyARPP19 and XeARPP19 were first injected into *Xenopus* prophase-arrested oocytes. Oocytes were collected 30 min later and phosphorylation of the PKA sites were estimated by western blot (Fig. 9A, Supp Fig. S11). For each of 3 experiments (A, B and C), 3 pools of 5 oocytes were analyzed, with *in vitro* thiophosphorylated forms of ClyARPP19 and XeARPP19 included as positive controls. Injected XeARPP19 became phosphorylated on S109 in the oocyte, and this phosphorylation was prevented by prior injection of PKI, confirming that PKA is entirely responsible for it (Fig. 9). In contrast, no phosphorylation on S81 of injected ClyARPP19 was detected, despite the fact that the protein remained stable in the oocytes and detectable by anti-GST antibodies (Fig. 9). These results show that, while XeARPP19 is actively phosphorylated by PKA in prophase-arrested *Xenopus* oocytes, ClyARPP19 is not.

**Figure 9.**
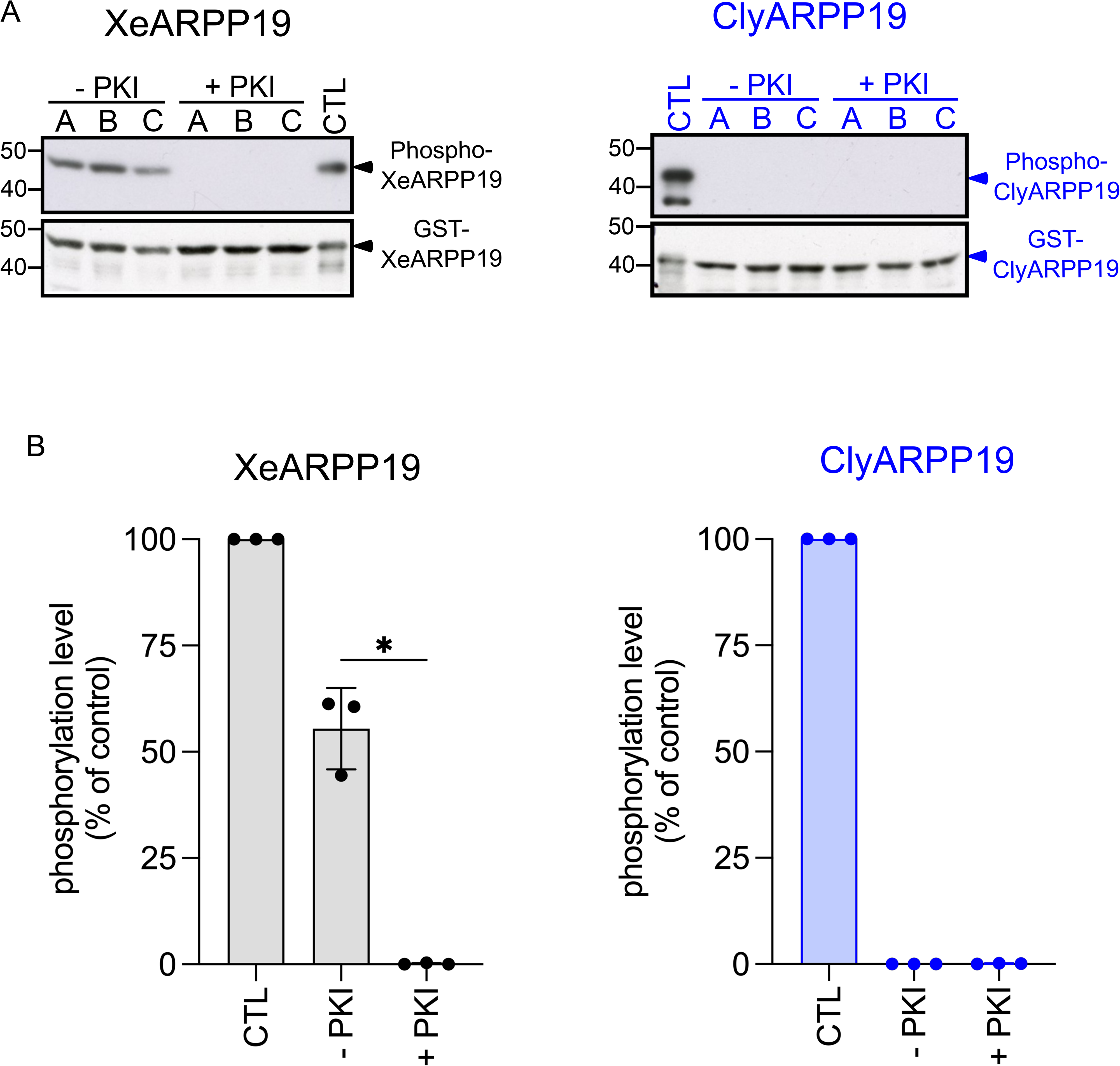
Phosphorylation of XeARPP19 and ClyARPP19 by PKA in *Xenopus* oocytes. **(A)** XeARPP19 or ClyARPP19 were injected into *Xenopus* oocytes previously injected or not with PKI. 30 min after ARPP19 injection, injected ARPP19 proteins were recovered by GST pull-down and their phosphorylation was monitored by western blot using antibodies against phosphoS109-XeARPP19 or phosphoS81-ClyARPP19. The total ARPP19 amount was detected with an anti-GST antibody. Three groups of 5 oocytes (A, B and C) were analyzed in both conditions. An *in vitro* phosphorylated form of ARPP19 was used as a positive control of phosphorylation (CTL). One representative experiment from 3 independent experiments is shown. **(B)** Quantification of the phosphorylation level of S109-XeARPP19 or S81-ClyARPP19 expressed as the % of the phosphorylation of the positive control. The quantifications were made from 3 independent experiments performed as in (A). Data are shown as mean +/- SD. Each dot represents one experiment. Ordinary one-way ANOVA: +PKI/-PKI Pvalue<0.0001.

The catalytic subunit of PKA (PKA-C) is highly conserved in the animal kingdom, the sequence being almost identical across species (Supp Fig. S12). The few differences in the sequences lie outside the critical motifs of PKA (catalytic loop, activation loop, Mg^2+^-binding loop or the two P-loops, Supp Fig. S12). Hence, it is unlikely that divergence between the catalytic subunits of the kinase prevents *Xenopus* or bovine PKA from phosphorylating ClyARPP19 in the experiments reported above, or *Clytia* PKA from phosphorylating XeARPP19. PKA activation in hydrozoan oocytes is induced by MIH (Amiel & Houliston, 2009; Deguchi *et al*., 2011; Takeda *et al*., 2006). We confirmed the increased PKA activity following MIH stimulation in *Clytia* oocytes using an antibody recognizing the phosphorylated PKA consensus site, which detected a set of protein bands more strongly in *Clytia* oocytes harvested 15 minutes post-MIH than in non-stimulated oocytes (Fig. 3A, B). Furthermore, a specific phospho-S109-XeARPP19 antibody detected phosphorylation of XeARPP19 injected into *Clytia* oocytes following MIH stimulation (Fig. 3A). In contrast, a distinct phospho-S81-ClyARPP19 antibody detected weak phosphorylation of exogenous ClyARPP19 injected to *Clytia* both before and after MIH, with no increase in signal intensity (Fig. 3B).

These experiments together demonstrate that *Clytia* ARPP19 is inherently a poorer PKA substrate than *Xenopus* ARPP109 both *in vivo* and *in vitro*, despite the presence of a functional PKA site. The endogenous rise in PKA activity in *Clytia* in MIH-stimulated oocytes likely generates little or no phosphorylation on this site.

## DISCUSSION

Maintenance and release of the prophase arrest of oocytes play a critical role in metazoan sexual reproduction. Correct regulation of these processes is of prime importance for the subsequent transformation of ovarian oocytes into fertilisable haploid eggs equipped with the maternal reserves necessary for early embryogenesis. cAMP and PKA activity play a central role in controlling the prophase arrest in many species, but with opposite roles in vertebrates versus many invertebrates (Deguchi *et al*., 2011; Jessus *et al*., 2020). Since ARPP19 is one of the substrates mediating the negative role of PKA on meiosis resumption in *Xenopus* (Dupre *et al*., 2014; Lemonnier *et al*., 2021), we conducted a comparative study of the phosphorylation and function of ARPP19 homologues between *Xenopus* and a species where PKA plays an opposite role, the hydrozoan jellyfish *Clytia* (Amiel & Houliston, 2009; Freeman & Ridgway, 1988; Takeda *et al*., 2006). Although a consensus sequence for phosphorylation by PKA can be recognised in both species, we show here by *in vitro* and *in vivo* approaches that ClyARPP19 is a weak substrate of the two enzymes that regulate its phosphorylation in *Xenopus*, PKA and PP2A-B55δ. Furthermore, the targets of XeARPP19 that allow it to exert its negative control of meiosis resumption are not present in the *Clytia* oocyte. As discussed below, these findings provide an evolutionary perspective on the control of the release of the oocyte prophase arrest by PKA.

The evolutionary history of the ARPP19 PKA site contrasts with that of the site phosphorylated by the kinase, Gwl (Gharbi-Ayachi *et al*., 2010; Mochida *et al*., 2010). The Gwl phosphorylation motif is highly conserved across eukaryotes and can be used as a hallmark of the ARPP19/ENSA family (Labandera *et al*., 2015). It is well established that across a very wide range of eukaryotic species including yeast and man, the serine of this short sequence is phosphorylated by Gwl, converting ARPP19 into an inhibitor of PP2A-B55δ that is essential for Cdk1 activation and M-phase completion (Castro & Lorca, 2018; Haccard & Jessus, 2011). Our work confirms that phosphorylation by Gwl of the central serine of this motif in ClyARPP19, S49, inhibits PP2A-B55δ activity. This property can be correlated with the conservation of several basic (K or R) residues flanking the Gwl site across eukaryotic species, previously reported to serve as a recognition signal for PP2A-B55 (Cundell *et al*, 2016; Labbe *et al*., 2021). Other conserved features of the ARPP19 proteins may have roles in modulating their phosphorylation by Gwl and thereby PP2A inhibition in different physiological contexts. Common sequences conserved across animal groups and even more widely, are candidates for such roles. A GxxxPTPxxϕP sequence is present across the opisthokonts, although it has degenerated in some animal lineages including ctenophores and sponges, while a C-terminal SxL sequence can be detected in species across all metazoan groups and amongst their closest unicellular relatives. It would be interesting in the future to address the significance of these sites by functional studies.

The XeARPP19 sequence contains only one site fitting the PKA phosphorylation consensus (Dupre *et al*., 2014). In contrast to the Gwl site, this site of PKA phosphorylation is not an ancient eukaryotic feature. We found the central serine in ARPP19 sequences from a range metazoan species, including from the most basally branching clade *Ctenophora* (Schultz *et al*., 2023), but no phosphorylatable residues at this position in ARPP19 proteins from non-metazoan species including choanoflagellates, plant and fungi. Under the likely scenario that ctenophores are the sister to all other animals (Schultz *et al*., 2023), this suggests that PKA phosphorylation of ARPP19 arose in a common metazoan ancestor, with rapid evolution of the C-ter half of the protein within the sponge clade leading to loss of this site as well as the GxxxPTPxxϕP motif (but not the C-terminal SxL). We have shown that this site is indeed phosphorylatable by PKA in a cnidarian ARPP19, from *Clytia hemisphaerica*. The emergence of a functional PKA phosphorylation site as a feature of ARPP19 proteins early during the emergence of metazoans and its subsequent maintenance across the clade of Eumetazoans (Cnidaria and Bilateria), suggests that phosphorylation on this site is involved in an important biological process common to these animals. One possibility is that it relates to the emergence of neural cell types in animals. ARPP19 and its splice-variant, ARPP16, were initially discovered in mammals as substrates of PKA in dopaminergic neurons of the striatum (Dulubova *et al*., 2001; Horiuchi *et al*., 1990). Since then, it has been shown that a complex antagonistic interplay between the control of ARPP16 by Gwl and PKA regulates key components of the striatal signaling by a mechanism whereby cAMP mediates PP2A disinhibition (Andrade *et al*, 2017; Musante *et al*, 2017). In the *Clytia* medusa, ARPP19 is expressed in the nerve rings that run around the bell margin as well as in ovarian oocytes. Thus, the appearance of the PKA site in the ARPP19 proteins at the time of the emergence of metazoans could have participated, thanks to its interaction with the pre-existing Gwl site, in the establishment of neural-type communication systems, which have been largely lost in sponges but have diversified in parallel in ctenophores (Burkhardt *et al*, 2023).

We have addressed another critical physiological feature common to all metazoans, the oocyte prophase arrest, which is regulated by PKA through the phosphorylation of ARPP19 in *Xenopus* (Lemonnier *et al*, 2020). In *Clytia*, PKA activation is required for release from prophase and Cdk1 activation (Amiel & Houliston, 2009; Freeman & Ridgway, 1988; Takeda *et al*., 2006). If ClyARPP19 was phosphorylated by PKA as in *Xenopus*, and if the targets of the PKA-phosphorylated form were expressed in the *Clytia* oocyte, how to explain that the protein does not block meiosis resumption in the *Clytia* oocyte as it does in *Xenopus*? In this paper we uncover two main explanations for this apparent paradox. First, the ability of PKA to phosphorylate ClyARPP19 is lower than for the *Xenopus* orthologue, both *in vitro*, in oocyte extracts and in the oocyte. *Xenopus* and *Clytia* catalytic subunits of PKA being almost identical, the differences in the ability of *Xenopus* and *Clytia* ARPP19 to become phosphorylated should reflect the ARPP19 sequences. Similarly, ClyARPP19 is a poor substrate of the phosphatase(s) acting on the PKA phosphosite, mainly PP2A-B55δ. Hence, in contrast to *Xenopus*, the two antagonistic enzymes controlling S81 phosphorylation are not very effective towards *Clytia* ARPP19. Unsurprisingly, the activity of the two enzymes is balanced, preventing their substrate from being in a permanent state of phosphorylation or dephosphorylation. On the other hand, the affinity of the two antagonistic enzymes for ARPP19 is very different in these species despite the presence of a similar motif, endowed with the characteristics of a PKA consensus site. This highlights the importance of studying in more detail variations within the consensus motif and in their surrounding sequences that may modulate its affinity for PKA and PP2A-B55δ, as was done for Gwl flanking sequences (Cundell *et al*., 2016; Labbe *et al*., 2021). It would also be instructive to analyse PKA phosphorylatability of ARPP19 proteins from other species previously demonstrated to show cAMP-induced oocyte maturation, such as the annelid *Pseudopotamilla* and the brittle star *Amphipholis* (Deguchi *et al*., 2011). Another avenue of investigation would be to address the impact of PKA site regulation on the access of the Gwl site to its two regulatory enzymes, Gwl and PP2A-B55δ. It is clear that feedback occurs between the two sites, but the sequence elements responsible for these interactions remain to be identified (Dupre *et al*., 2017; Labbe *et al*., 2021). Thus, a comparison between *Clytia* and *Xenopus* ARPP19, respectively offering a site of low or high affinity for the PKA/PP2A-B55δ tandem, could shed light on the sequences outside the PKA and Gwl sites that regulate both their phosphorylation level and their reciprocal influences.

The low affinity of PKA for ClyARPP19 can explain why the protein does not interfere with the prophase release of the *Clytia* oocyte when PKA is activated by MIH. However, another mechanism relying on the ARPP19 interactors also protects the *Clytia* oocyte from a potentially negative action of S81-phosphorylated ARPP19. The nature of the proteins controlled by the S109 phosphorylation of XeARPP19 responsible for maintaining the prophase block in *Xenopus* oocytes remain unknown but their presence can be deduced experimentally. First, our results indicate that ClyARPP19 interacts only weakly with these *Xenopus* effectors since wild-type ClyARPP19 or a ClyARPP19 mutant mimicking a constitutive S81-phosphorylation are not as efficient as their *Xenopus* counterparts in blocking meiotic resumption of *Xenopus* oocytes. Second, they suggest that their *Clytia* homologs are not expressed or have divergent structures in *Clytia* oocytes since neither the injection of S81-phosphomimetic ClyARPP19 nor the injection of S109-phosphomimetic XeARPP19 prevented MIH-induced oocyte maturation in *Clytia*. In addition, Gwl-phosphorylated forms of *Xenopus* (S67) or of *Clytia* (S49) ARPP19 were not able to trigger meiotic resumption of *Clytia* oocytes, supporting the hypothesis that additional mechanisms lock the resting *Clytia* oocyte in a prophase state. OA, which has a broader spectrum towards PP2A isoforms than ARPP19 (Dounay & Forsyth, 2002), was also unable to induce GVBD when injected into *Clytia* oocytes, but did provoke cortical contractions and lysis likely reflecting PP2A regulation of the actin cytoskeleton as reported in other cells (Basu, 2011; Hoffman *et al*, 2017). Taken together, our results suggest that the low capacity of *Clytia* ARPP19 to be phosphorylated by PKA combined with the absence of functional interactors mediating its negative effects on Cdk1 activation may provide a double security allowing induction of meiosis resumption in *Clytia* by elevated PKA activity despite the presence of ARPP19, while additional as yet unidentified mechanisms ensure the *Clytia* oocyte prophase arrest.

What can we conclude from this study about the evolution of oocyte maturation initiation mechanisms? It is likely that the ancestral ARPP19 was expressed in oocytes of early metazoans to ensure MPF activation through its highly conserved eukaryotic role in the Gwl/PP2A regulatory circuit, and this ancestral ARPP19 had a “prototype” PKA phosphorylation site (Fig. 10), but it remains to be established how strongly this was regulated and its role in the oocyte. One scenario consistent with our findings is that in the earliest metazoan, the ARPP19 proteins present in the oocyte had no role in the maturation initiation step; cAMP-PKA would have played a positive role in initiating oocyte maturation, perhaps relating to an ancient involvement of a GPCR related to the *Clytia* MIH receptor (Quiroga Artigas *et al*., 2020). Such ARPP19-independent cAMP/PKA-activated oocyte maturation initiation would have been maintained in hydrozoan cnidarians and in some protostomes (eg *Spisula* and *Pseudopotamilla*) and non-vertebrate deuterostomes (eg *Amphipholis* and *Boltenia*). Maturation initiation is tightly linked to spawning/ovulation and so its regulation would have been modulated repeatedly during evolution in relation to the ecology and ethology of each species (for instance integrating the influence of daily and lunar cycles, nutritional availability, etc.) The emergence of the vertebrate lineage was accompanied by major changes in reproductive biology linked to the complexification of the hormone systems regulating ovulation (Deguchi *et al*., 2011; Quiroga Artigas *et al*., 2020). If cAMP-PKA activation of oocyte maturation was indeed ancestral, this ancient initiation mechanism would have become over-ridden by an alternative regulation in which cAMP-PKA inhibits meiotic maturation. Under this scenario, the biochemical pathways within the oocyte linking the initiation trigger to Cdk1 activation would have undergone modifications during the emergence of the vertebrate lineage, including the co-option of ARPP19 for an inhibitory role under the control of cAMP-PKA. Whatever the ancestral situation for oocyte maturation control, it is highly likely that cAMP-PKA of ARPP19 initially arose during metazoan evolution for other functions, for instance in the nervous system. Consistent with this, we detected expression of ARPP19 in both the oocytes and the nerve ring of *Clytia* jellyfish.

**Figure 10.**
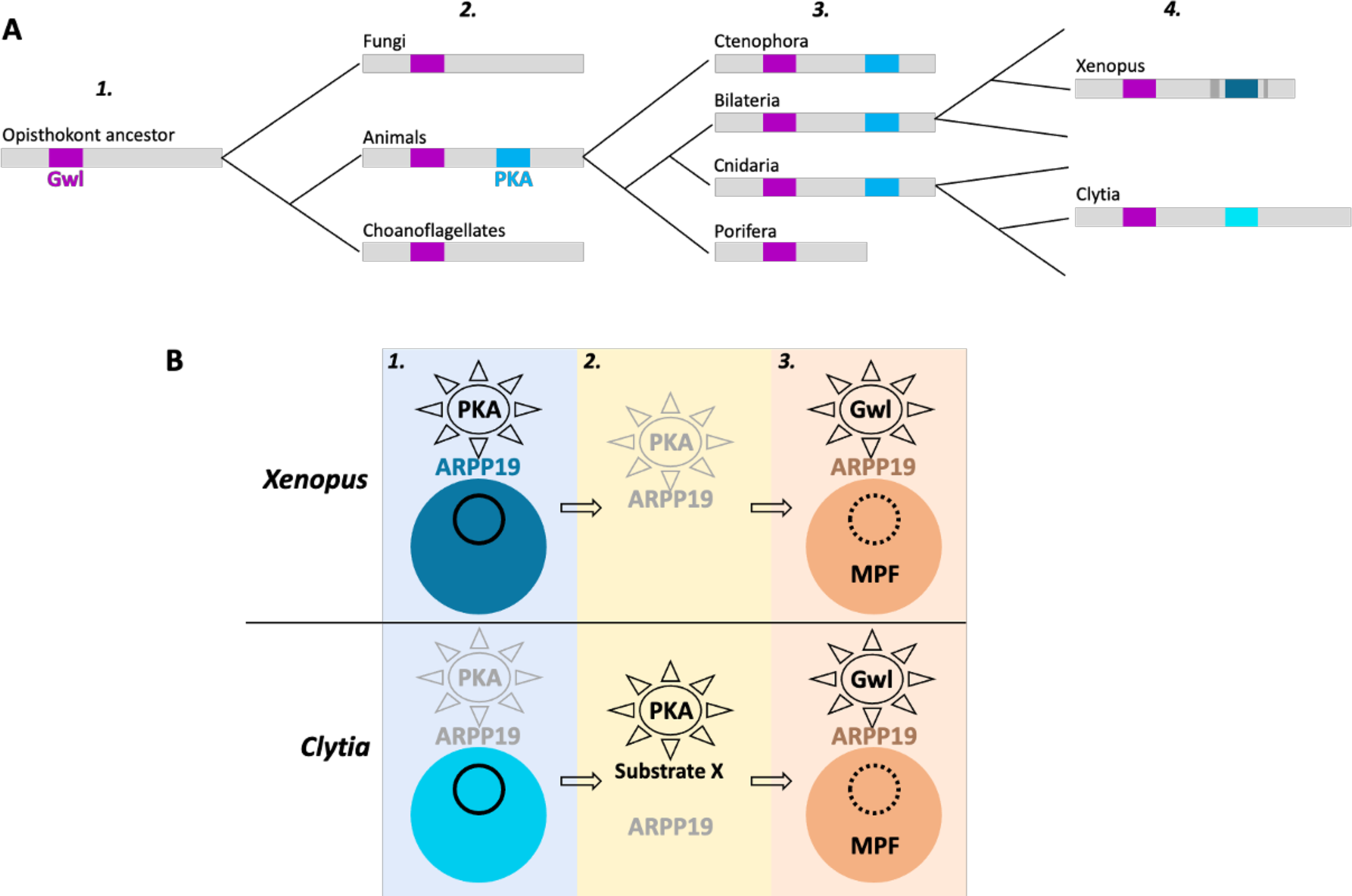
Complexification of ARPP19 phosphorylation sites and functions in oocyte maturation during animal evolution. **(A)** Evolutionary history of the PKA and Gwl sites of ARPP19 among Opisthokonts. ***1.*** A widely conserved Gwl phosphorylation site among eukaryotes, hence present in the opisthokont ancestor. ***2.*** Arising of a consensus PKA phosphorylation site early in the clade Metazoa. It could play a biological role common to these animals, but not related to oocyte meiotic maturation. ***3.*** Loss of the PKA site in *Porifera*. ***4.*** Species specific changes in the phosphorability of the PKA site: strong in *Xenopus*, weak in *Clytia*. **(B)** Functions of ARPP19 in oocyte meiotic maturation. ***1.*** *Prophase arrest.* In *Xenopus*, PKA is active and phosphorylates ARPP19 that indirectly inhibits MPF activation by unknown mechanisms. In *Clytia*, PKA is inactive and ARPP19, even if its phosphorylation by PKA is artificially mimicked, is unable to inhibit MPF. ***2.*** *MIH stimulation and meiotic maturation initiation.* In *Xenopus*, progesterone causes a drop of PKA activity, ARPP19 is dephosphorylated, releasing the activatory pathway of MPF. In *Clytia*, MIH activates PKA whose hypothetical substrate X activates MPF. ***3.*** *Entry into M-phase*. In both species, Gwl phosphorylates ARPP19 that plays its highly conserved action as a PP2A-B55 inhibitor, essential for MPF activity.

To address this hypothesis much remains to be clarified about the phylogenetic distribution of cAMP regulation modes of oocyte maturation initiation. From the relatively sparse available data it is already clear that this is quite patchy: not all protostome oocytes use cAMP-PKA to trigger meiosis resumption and not all deuterostome use cAMP-PKA to maintain the prophase block. Thus, whatever the ancestral situation, there were likely multiple acquisitions or losses of the activatory role of cAMP-PKA in oocyte meiotic maturation during the evolution of metazoans. We propose that the new and critical function in oocytes of vertebrates and of some other protostome lineages complemented existing roles for ARPP19 under the control of PKA, that emerged during early metazoan evolution (Fig. 10). More knowledge of the sequence modifications around the PKA site of ARPP19 and the identification of its binding partners will be important to understand fully these evolutionary transitions. This will increase our understanding of the function of ARPP19 in the control of oocyte meiosis, and more widely in the control of cell cycle progression and its dysfunction in pathologies related to fertility and cancer.

## MATERIAL AND METHODS

### Material

*Xenopus laevis* adult females (Centre de Ressources Biologiques Xenopes, CNRS, France) were bred and maintained according to current French guidelines in the IBPS aquatic animal facility, with authorization: Animal Facility Agreement: #A75-05-25. All *Xenopus* experiments were subject to ethical review and approved by the French Ministry of Higher Education and Research (reference APAFIS#14127-2018031614373133v2). Sexually mature jellyfish were generated from laboratory-maintained *Clytia hemisphaerica* polyp colonies (“A strains” or “Z strains”) (Houliston *et al*, 2010). All reagents, unless otherwise specified, were from Roth.

### *Xenopus* oocyte handling

*Xenopus laevis* stage VI oocytes (Dumont, 1972) were obtained from unprimed female. The edges of the ovarian lobes were collected from anesthetized *Xenopus* females (30 minutes bath in 1 g/l MS222 (Sigma). To obtain defolliculated fully-grown oocytes, pieces of ovaries were incubated for 3 hours in buffer M (10 mM HEPES pH 7.8, 88 mM NaCl, 1 mM KCl, 0.33 mM Ca(NO3)_2_, 0.41 mM CaCl_2_, 0.82 mM MgSO_4_) in the presence of dispase (0.4 mg/ml) to weaken ovary connective tissues without altering oocyte integrity and potentiate the effect of collagenase used in the sub-sequent step. Oocytes were then washed in 1 liter of buffer M and incubated for 1 hour in the presence of collagenase (0.4 mg/ml) diluted in buffer M. After washing with 2 liters of buffer M to eliminate collagenase, the oocytes were sorted by size to collect stage VI (≍ 1200 µm of diameter). They were kept in buffer M at 16°C for experiments and then immediately lysed or stored at -80°C.

Prophase-arrested oocytes were micro-injected with the following recombinant proteins (injection volume: 50 nL per oocyte): 100 ng or 800 ng of GST-XeARPP19, GST-ClyARPP19, GST-S109D-XeARPP19 or GST-S81D-ClyARPP19; 200 ng of GST-S49thio-ClyARPP19 or GST-S67thio-XeARPP19; 75 ng of PKI.

Meiotic maturation was induced by 2 µM progesterone. Oocytes were referred to as GVBD when the first pigment rearrangement was detected at the animal pole. The percentage of oocytes at GVBD over the time, calculated for group of 20 to 30 injected oocytes, was then fitted with a four-parameter logistic regression whose the equation is the following:

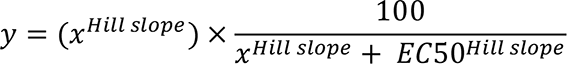

### *Clytia* oocyte handling

*Clytia hemisphaerica* fully grown (stage III) oocytes were obtained from manually isolated ovaries of jellyfish cultured overnight in MFSW (Millipore filtrated seawater). The harvested gonads in were opened lengthwise using fine tweezers. Oocytes were then recovered using 0.3 µm diameter tungsten wire loops mounted on glass capillaries. Meiotic maturation was induced by MIH treatment (100 nM WPRP-NH_2_(Takeda *et al*., 2018). GVBD was scored every 5 minutes under Zeiss dissecting microscopes, or from time lapse films recorded using a Zeiss Axiobserver with DIC optics. Protein injection into *Clytia* isolated oocytes was performed using Nanoject compressed air microinjection systems (Eppendorf) (Momose & Houliston, 2007) in continuous flow mode. Recombinant proteins and OA (Enzo Life Sciences) were diluted in PBS and centrifuged at 14,000 rpm at 4°C for 5 minutes before injection. Solutions of 2-4 mg/ml of each recombinant ARRP19 protein or 2 µM OA were injected, aiming at a delivering 3-5% oocyte volume based on the size of the cloud of injected liquid. ADD

### Cloning and recombinant protein purification

Construction of plasmids encoding GST-tagged XeARPP19, S109D-XeARPP19, S67A-XeARPP19 and PKI were previously described (Dupre *et al*., 2014). A cDNA fragment encoding full ORF of *Clytia* ARPP19, identified from gene models in the *Clytia* genome assembly and associated transcriptome data (Leclere *et al*., 2019) (Supp Fig. S1A and S1C) was amplified by PCR from medusa gonad cDNA and subcloned into pGex4-T1 vector to express GST-tagged ClyARPP19. S49A, S81D, S82D and S81D-S82D-ClyARPP19 were generated according to the manufacturer protocol (Stratagene) and sequences were verified by DNA sequencing (Genewiz-Azenta, Germany). The cDNA of 6XHIS-tagged S109A-XeARPP19 has been subcloned into pET14b vector. Both vectors were used to transform BL21 strain of *E. coli* and GST-tagged or 6XHIS-tagged recombinant proteins were produced by autoinduction (Studier, 2005). Bacteria were lysed by sonification in PBS pH 7,4 (13.7 mM NaCl, 2.7 mM KCl, 4.3 mM KH_2_PO_4_, 1.4 mM Na_2_HPO_4_) with 1 mg/ml lysozyme in the presence of 10% Triton. After a brief centrifugation step, supernatants were loaded either onto an equilibrated glutathione-agarose column (Sigma) for GST-tagged protein or on Nickel beads column (Qiagen) for 6XHIS-tagged protein. Columns were then washed and eluted with respectively 10 mM glutathione in PBS or 160 mM imidazole diluted in 1 M of β-glycerophosphate. The purified proteins were concentrated against polyethyleneglycol, dialyzed overnight against PBS and stored at −80°C.

### Clytia in situ hybridization

For *in situ* hybridization, female jellyfish were fixed in 3.7% formaldehyde, 0.2% glutaraldehyde in PBS on ice for 60-120 min. They were then washed 5 times with PBST (PBS + 0.1% Tween20), dehydrated stepwise in methanol and stored in 100% methanol at -20°C. Hybridization (at 62°C for 72h) and washing steps were performed in a robot (Intavis AG, Bioanalytical Instruments) using 20X SSC pH adjusted to 4.7. Acetylation steps using 0.1M triethanolamine in PBST (2×5min), then 0.25% acetic anhydride in 0.1M triethanolamine (2×5min) followed by PBST washes (3×10min) were included before pre-hybridization to reduce probe non-specific binding. Incubation with Anti-DIG AP, 1:2000 in 1X blocking solution was performed for 3h and the NCB-BCIP performed at pH 9, monitoring colour development under the binocular microscope. Following post-fixation, washing and equilibration of samples in 50% glycerol/PBS, Images were acquired on an Olympus BX51 microscope.

### ARPP19 *in vitro* phosphorylation or thiophosphorylation by Gwl and PKA

Active Gwl was obtained by injecting *Xenopus* prophase oocytes with mRNA encoding Histidine-tagged *Xenopus* K71M-Gwl (Dupre *et al*., 2013). Oocytes were collected at metaphase II and K71M-Gwl was recovered by incubating the oocyte extract with Nickel beads in the presence of 1 μM OA (Enzo Life Sciences). Nickel beads were washed in kinase buffer (20 mM HEPES pH 7.4, 2 mM 2-Mercaptoethanol). Recombinant 400 μM GST-ARPP19 or 6XHIS-S109A-XeARPP19 was added to the beads in the presence of 1 mM γS-ATP. The reaction was performed in a final volume of 60 μL for 60 min at 30°C under stirring (750 RPM, Thermomixer, Eppendorf). γS-ATP was removed by dialyzing the reaction mix against kinase buffer. For PKA thiophosphorylation, 3 nM of either XeARPP19 or various forms of ClyARPP19 (stock solutions at 1 μg/μL) were incubated in the presence of 62.5 units of recombinant bovine PKA (Promega) and 1 mM γS-ATP in a final volume of 30 μL of PKA Buffer (20 mM HEPES pH 7.4, 20 mM MgCl_2_), at 37°C under stirring (750 RPM, Thermomixer, Eppendorf). At indicated times, 5 μL of reaction were sampled, supplemented with Laemmli buffer (Laemmli, 1970) and heated for 1 minute at 90°C. To be detected by the thiophosphate-ester antibody, thiophosphates incorporated in proteins were then subjected to alkylation by incubation with 1 mM P-nitrobenzyl mesylate (PNBM) (Cayman Chemical) for 1 hour at room temperature. Samples were then stored at -20°C. Positive controls of XeARPP19 and ClyARPP19 phosphorylation used in various experiments were obtained by incubating 3 nM of XeARPP19 and ClyARPP19 for respectively 30 and 180 minutes to reach a similar thiophosphorylation level for both proteins. In some experiments, XeARPP19 and ClyARPP19 were phosphorylated by PKA. 100 ng of either XeARPP19 or ClyARPP19 were incubated with 200 μM ATP and 25 units of recombinant bovine PKA (Promega) in PKA Buffer for 3 hours at 37°C under 1200 RPM stirring. The reaction was stopped by Laemmli buffer (Laemmli, 1970) and heating for 1 minute at 90°C.

### ARPP19 phosphorylation and dephosphorylation in *Xenopus* oocyte extract

Prophase-arrested oocytes were lysed in 10 volumes of Extraction buffer (80 mM β-glycerophosphate pH 7.3, 20 mM EGTA, 15 mM MgCl_2_). Lysates were centrifuged for 15 minutes at 12,000 rpm and 4°C and the supernatant was then used as the “oocyte extract”. For PKA phosphorylation, 50 ng (25 µM) of either GST-XeARPP19 or GST-ClyARPP19 (stock solution of 1 μg/μL) were added in 50 μL of prophase extract previously incubated 30 min or not with 75 ng PKI, and further incubated at 30°C, under 750 RPM stirring. 5 μL of extract were sampled every 5 minutes, supplemented with Laemmli buffer (Laemmli, 1970), heated at 90°C for 1 minute and stored at -20°C. To measure ARPP19 dephosphorylation, “kinase-dead” oocyte extracts, were generated by addition of 75 ng PKI, 0,1 units/ml hexokinase (Sigma) and 10 mM glucose to deplete endogenous ATP (Newmeyer *et al*, 1986) followed by a 1 hour incubation at 18°C. In some experiments, kinase-dead extracts were incubated 3 consecutive times with protein A-beads (Bio-Rad) previously coated with the anti-B55δ antibody (1 hour incubation at 4°C, followed by extensive washes) and Nickel beads (Qiagen) coated with 400 μg of 6XHIS-S67thio-S109A-XeARPP19. The extract was then incubated with protein A-beads to deplete free IgG against B55δ. 250 ng of GST-phosphoS109-XeARPP19 or GST-phosphoS81-ClyARPP19 (stock solution of 1 μg/μL) were then added in 4 μL of kinase-dead oocyte extract and incubated at 30°C under 1250 RPM stirring. 3 μL of extract were collected every 5 minutes, supplemented with Laemmli buffer (Laemmli, 1970), heated and stored at -20°C. All western blot signals were quantified using Image J software. In assays using recombinant GST-ARPP19, each phosphorylation or thiophosphorylation signal was divided by the corresponding GST signal to obtain a ratio (normalized signals).

*Dephosphorylation assays (Figs 2 and 6)*: the normalized signals were standardized on the time 0 minute that was set as 100% of phosphorylation. Standardized signals were fitted with a one-phase decay equation (Supp Fig. S8):

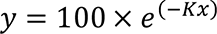

The half-time for each experiment, defined as the time of 50% of dephosphorylation, was calculated with the following equation (Supp Fig. S8):

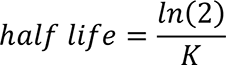

Comparison of the half-times of dephosphorylation between XeARPP19 and ClyARPP19 was done by applying a paired T-test.

*Phosphorylation assays (Fig. 8)*: the normalized signals were standardized on the time 60 minutes of XeARPP19 that was considered as 100% phosphorylation. Standardized signals were fitted with a one-phase association equation (Supp Fig. S10):

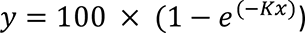

For each experiment, the half-time is defined as the time of 50% of phosphorylation and was calculated with the following equation (Supp Fig. S10):

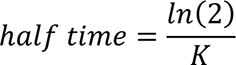

Comparison of the half-times of phosphorylation between XeARPP19 and ClyARPP19 was done by applying a paired T-test. *: P≤0.05; **: P≤0.01; ***: P≤0.001; ****: P≤0.0001; P>0.05: not significant (ns).

### Antibodies and western blots

The equivalent of 0.5 oocyte or 50 ng of Xe-ARPP19 and ClyARPP19 (*in vitro* and *in vivo* phosphorylation/dephosphorylation assays) were loaded on 12% acrylamide gel, subjected to SDS gel electrophoresis (Laemmli, 1970) and then transferred onto nitrocellulose membranes (Dupre *et al*, 2002). The antibodies directed against the following proteins were used: phosphoS109-XeARPP19 (Dupre *et al*., 2014) (1:100,000 for *Xenopus* proteins or 1:500 for *Clytia* proteins), phosphoS81-ClyARPP19 (1:160,000 for *Xenopus* proteins or 1:500 after retro-elution for *Clytia* proteins, see Supp Fig. S6 for validation), B55δ (see Supp Fig. S9 for validation), thiophosphate ester (1:60,000, Abcam ab92570), GST (1:5,000, Sigma A-7340), karyopherin (1:1,000, Santa-Cruz biotechnology sc-1863), phospho-MAPK (1:1,000, Cell Signaling 9106), phosphoY15-Cdk1 (1:1,000, Cell Signaling 9111) and phosphorylated-PKA substrates (1:1000, Cell Signaling 9624). After overnight incubation at 4°C, nitrocellulose membranes were incubated 1 hour at room temperature in the appropriate horse-radish peroxidase-labeled secondary antibodies (Jackson Immunoresearch) and revealed by chemiluminescence (Pierce). All western blots are representative of at least three different experiments.

### Bioinformatics and sequence alignments

ARPP19 protein sequences were retrieved from public databases. Accession numbers along with the *Clytia* ARPP19 protein and transcript sequence are provided in Supp Fig. S1. Alignments of different subsets of sequences were made using Clustal Omega on the EBI portal (www.ebi.ac.uk) and then compared and adjusted by eye.

## ACKNOWLEDGMENTS

We thank Julie Uviera and Gonzalo Quiroga-Artigas for their invaluable contributions to the *Clytia* oocyte injection experiments, and all members of the team “Oocyte Biology” for helpful discussions. This work was supported by the National Center for Scientific Research (CNRS), Sorbonne University and the National Research Agency (ANR grants 13-BSV2-0008-01 to C.J. and E.H. and 18-CE13-0013-01 to C.J.). F.M. received a PhD grant from the CNRS and from the ARC foundation (Association pour la Recherche sur le Cancer, grant ARCDOC42021120004303).

## REFERENCES

Amiel A, Houliston E (2009) Three distinct RNA localization mechanisms contribute to oocyte polarity establishment in the cnidarian Clytia hemisphaerica. Dev Biol 327: 191–203

Amiel A, Leclere L, Robert L, Chevalier S, Houliston E (2009) Conserved functions for Mos in eumetazoan oocyte maturation revealed by studies in a cnidarian. Curr Biol 19: 305–311

Andrade EC, Musante V, Horiuchi A, Matsuzaki H, Brody AH, Wu T, Greengard P, Taylor JR, Nairn AC (2017) ARPP-16 Is a Striatal-Enriched Inhibitor of Protein Phosphatase 2A Regulated by Microtubule-Associated Serine/Threonine Kinase 3 (Mast 3 Kinase). J Neurosci 37: 2709–2722

Basu S (2011) PP2A in the regulation of cell motility and invasion. Curr Protein Pept Sci 12: 3–11

Burkhardt P, Colgren J, Medhus A, Digel L, Naumann B, Soto-Angel JJ, Nordmann EL, Sachkova MY, Kittelmann M (2023) Syncytial nerve net in a ctenophore adds insights on the evolution of nervous systems. Science 380: 293–297

Castro A, Lorca T (2018) Greatwall kinase at a glance. J Cell Sci 131

Conti M, Andersen CB, Richard F, Mehats C, Chun SY, Horner K, Jin C, Tsafriri A (2002) Role of cyclic nucleotide signaling in oocyte maturation. Mol Cell Endocrinol 187: 153–159

Cundell MJ, Hutter LH, Nunes Bastos R, Poser E, Holder J, Mohammed S, Novak B, Barr FA (2016) A PP2A-B55 recognition signal controls substrate dephosphorylation kinetics during mitotic exit. J Cell Biol 214: 539–554

Deguchi R, Takeda N, Stricker SA (2011) Comparative biology of cAMP-induced germinal vesicle breakdown in marine invertebrate oocytes. Mol Reprod Dev 78: 708–725

Dounay AB, Forsyth CJ (2002) Okadaic acid: the archetypal serine/threonine protein phosphatase inhibitor. Curr Med Chem 9: 1939–1980

Dulubova I, Horiuchi A, Snyder GL, Girault JA, Czernik AJ, Shao L, Ramabhadran R, Greengard P, Nairn AC (2001) ARPP-16/ARPP-19: a highly conserved family of cAMP-regulated phosphoproteins. J Neurochem 77: 229–238

Dumont JN (1972) Oogenesis in Xenopus laevis (Daudin). I. Stages of oocyte development in laboratory maintained animals. J Morphol 136: 153–179

Dupre A, Buffin E, Roustan C, Nairn AC, Jessus C, Haccard O (2013) The phosphorylation of ARPP19 by Greatwall renders the auto-amplification of MPF independently of PKA in Xenopus oocytes. J Cell Sci 126: 3916–3926

Dupre A, Daldello EM, Nairn AC, Jessus C, Haccard O (2014) Phosphorylation of ARPP19 by protein kinase A prevents meiosis resumption in Xenopus oocytes. Nature communications 5: 3318

Dupre A, Jessus C (2017) ARPP19 Phosphorylations by PKA and Greatwall: The Yin and the Yang of the Cell Decision to Divide. In: Protein Phosphorylation, Prigent C. (ed.) pp. 3-29. IntechOpen: Rijeka, Croatia

Dupre A, Jessus C, Ozon R, Haccard O (2002) Mos is not required for the initiation of meiotic maturation in Xenopus oocytes. EMBO J 21: 4026–4036

Dupre AI, Haccard O, Jessus C (2017) The greatwall kinase is dominant over PKA in controlling the antagonistic function of ARPP19 in Xenopus oocytes. Cell Cycle 16: 1440–1452

Eyers PA, Liu J, Hayashi NR, Lewellyn AL, Gautier J, Maller JL (2005) Regulation of the G(2)/M transition in Xenopus oocytes by the cAMP-dependent protein kinase. J Biol Chem 280: 24339–24346

Freeman G, Ridgway EB (1988) The role of cAMP in oocyte maturation and the role of the germinal vesicle contents in mediating maturation and subsequent developmental events in hydrozoans. Rouxs Arch Dev Biol 197: 197–211

Gharbi-Ayachi A, Labbé J-C, Burgess A, Vigneron S, Strub J-M, Brioudes E, Van-Dorsselaer A, Castro A, Lorca T (2010) The Substrate of Greatwall Kinase, Arpp19, Controls Mitosis by Inhibiting Protein Phosphatase 2A. Science 330: 1673-1677

Goris J, Hermann J, Hendrix P, Ozon R, Merlevede W (1989) Okadaic acid, a specific protein phosphatase inhibitor, induces maturation and MPF formation in Xenopus laevis oocytes. FEBS Lett 245: 91–94

Grau-Bove X, Torruella G, Donachie S, Suga H, Leonard G, Richards TA, Ruiz-Trillo I (2017) Dynamics of genomic innovation in the unicellular ancestry of animals. eLife 6

Haccard O, Jessus C (2011) Greatwall Kinase, ARPP-19 and Protein Phosphatase 2A: Shifting the Mitosis Paradigm. Results Probl Cell Differ 53: 219-234

Hoffman A, Taleski G, Sontag E (2017) The protein serine/threonine phosphatases PP2A, PP1 and calcineurin: A triple threat in the regulation of the neuronal cytoskeleton. Mol Cell Neurosci 84: 119-131

Horiuchi A, Williams KR, Kurihara T, Nairn AC, Greengard P (1990) Purification and cDNA cloning of ARPP-16, a cAMP-regulated phosphoprotein enriched in basal ganglia, and of a related phosphoprotein, ARPP-19. The Journal of biological chemistry 265: 9476-9484

Houliston E, Momose T, Manuel M (2010) Clytia hemisphaerica: a jellyfish cousin joins the laboratory. Trends Genet 26: 159–167

Huchon D, Ozon R, Fischer EH, Demaille JG (1981) The pure inhibitor of cAMP-dependent protein kinase initiates Xenopus laevis meiotic maturation. A 4-step scheme for meiotic maturation. Mol Cell Endocrinol 22: 211-222

Jessus C, Munro C, Houliston E (2020) Managing the Oocyte Meiotic Arrest-Lessons from Frogs and Jellyfish. Cells 9

Jessus C, Rime H, Haccard O, Van Lint J, Goris J, Merlevede W, Ozon R (1991) Tyrosine phosphorylation of p34cdc2 and p42 during meiotic maturation of Xenopus oocyte. Antagonistic action of okadaic acid and 6-DMAP. Development 111: 813-820

Juanes MA, Khoueiry R, Kupka T, Castro A, Mudrak I, Ogris E, Lorca T, Piatti S (2013) Budding yeast greatwall and endosulfines control activity and spatial regulation of PP2A(Cdc55) for timely mitotic progression. PLoS genetics 9: e1003575

Kishimoto T (2018) MPF-based meiotic cell cycle control: Half a century of lessons from starfish oocytes. Proc Jpn Acad Ser B Phys Biol Sci 94: 180–203

Kovo M, Kandli-Cohen M, Ben-Haim M, Galiani D, Carr DW, Dekel N (2006) An active protein kinase A (PKA) is involved in meiotic arrest of rat growing oocytes. Reproduction (Cambridge, England) 132: 33–43

Labandera AM, Vahab AR, Chaudhuri S, Kerk D, Moorhead GB (2015) The mitotic PP2A regulator ENSA/ARPP-19 is remarkably conserved across plants and most eukaryotes. Biochem Biophys Res Commun 458: 739–744

Labbe JC, Vigneron S, Mechali F, Robert P, Roque S, Genoud C, Goguet-Rubio P, Barthe P, Labesse G, Cohen-Gonsaud M et al (2021) The study of the determinants controlling Arpp19 phosphatase-inhibitory activity reveals an Arpp19/PP2A-B55 feedback loop. Nature communications 12: 3565

Laemmli UK (1970) Cleavage of structural proteins during the assembly of the head of bacteriophage T4. Nature 227: 680–685

Lambert CC (2011) Signaling pathways in ascidian oocyte maturation: the roles of cAMP/Epac, intracellular calcium levels, and calmodulin kinase in regulating GVBD. Mol Reprod Dev 78: 726–733

Lechable M, Jan A, Duchene A, Uveira J, Weissbourd B, Gissat L, Collet S, Gilletta L, Chevalier S, Leclere L et al (2020) An improved whole life cycle culture protocol for the hydrozoan genetic model Clytia hemisphaerica. Biol Open 9

Leclere L, Horin C, Chevalier S, Lapebie P, Dru P, Peron S, Jager M, Condamine T, Pottin K, Romano S et al (2019) The genome of the jellyfish Clytia hemisphaerica and the evolution of the cnidarian life-cycle. Nature ecology & evolution 3: 801–810

Lemonnier T, Daldello EM, Poulhe R, Le T, Miot M, Lignieres L, Jessus C, Dupre A (2021) The M-phase regulatory phosphatase PP2A-B55delta opposes protein kinase A on Arpp19 to initiate meiotic division. Nature communications 12: 1837

Lemonnier T, Dupre A, Jessus C (2020) The G2-to-M transition from a phosphatase perspective: a new vision of the meiotic division. Cell Div 15: 9

Maller JL, Butcher FR, Krebs EG (1979) Early effect of progesterone on levels of cyclic adenosine 3’:5’-monophosphate in Xenopus oocytes. J Biol Chem 254: 579–582

Maller JL, Krebs EG (1977) Progesterone-stimulated meiotic cell division in Xenopus oocytes. Induction by regulatory subunit and inhibition by catalytic subunit of adenosine 3’:5’-monophosphate-dependent protein kinase. J Biol Chem 252: 1712-1718

Masui Y (2001) From oocyte maturation to the in vitro cell cycle: the history of discoveries of Maturation-Promoting Factor (MPF) and Cytostatic Factor (CSF). Differentiation 69: 1–17

Mochida S, Ikeo S, Gannon J, Hunt T (2009) Regulated activity of PP2A-B55 delta is crucial for controlling entry into and exit from mitosis in Xenopus egg extracts. EMBO J 28: 2777–2785

Mochida S, Maslen SL, Skehel M, Hunt T (2010) Greatwall phosphorylates an inhibitor of protein phosphatase 2A that is essential for mitosis. Science 330: 1670–1673

Momose T, Houliston E (2007) Two oppositely localised frizzled RNAs as axis determinants in a cnidarian embryo. PLoS biology 5: e70

Munro C, Cadis H, Pagnotta S, Houliston E, Huynh JR (2023) Conserved meiotic mechanisms in the cnidarian Clytia hemisphaerica revealed by Spo11 knockout. Sci Adv 9: eadd2873

Musante V, Li L, Kanyo J, Lam TT, Colangelo CM, Cheng SK, Brody AH, Greengard P, Le Novere N, Nairn AC (2017) Reciprocal regulation of ARPP-16 by PKA and MAST3 kinases provides a cAMP-regulated switch in protein phosphatase 2A inhibition. eLife 6

Nader N, Courjaret R, Dib M, Kulkarni RP, Machaca K (2016) Release from Xenopus oocyte prophase I meiotic arrest is independent of a decrease in cAMP levels or PKA activity. Development 143: 1926–1936

Newmeyer DD, Lucocq JM, Burglin TR, De Robertis EM (1986) Assembly in vitro of nuclei active in nuclear protein transport: ATP is required for nucleoplasmin accumulation. EMBO J 5: 501–510

Ozon R, Belle R, Huchon D, Mulner O (1979) Roles of cyclic AMP and calcium in maturation of Xenopus laevis oocytes. J Steroid Biochem 11: 709–713

Quiroga Artigas G, Lapebie P, Leclere L, Bauknecht P, Uveira J, Chevalier S, Jekely G, Momose T, Houliston E (2020) A G protein-coupled receptor mediates neuropeptide-induced oocyte maturation in the jellyfish Clytia. PLoS biology 18: e3000614

Quiroga Artigas G, Lapebie P, Leclere L, Takeda N, Deguchi R, Jekely G, Momose T, Houliston E (2018) A gonad-expressed opsin mediates light-induced spawning in the jellyfish Clytia. eLife 7

Schultz DT, Haddock SHD, Bredeson JV, Green RE, Simakov O, Rokhsar DS (2023) Ancient gene linkages support ctenophores as sister to other animals. Nature 618: 110–117

Stricker SA, Smythe TL (2001) 5-HT causes an increase in cAMP that stimulates, rather than inhibits, oocyte maturation in marine nemertean worms. Development 128: 1415–1427

Studier FW (2005) Protein production by auto-induction in high density shaking cultures. Protein Expr Purif 41: 207–234

Takai A, Bialojan C, Troschka M, Ruegg JC (1987) Smooth muscle myosin phosphatase inhibition and force enhancement by black sponge toxin. FEBS Lett 217: 81–84

Takeda N, Kon Y, Quiroga Artigas G, Lapebie P, Barreau C, Koizumi O, Kishimoto T, Tachibana K, Houliston E, Deguchi R (2018) Identification of jellyfish neuropeptides that act directly as oocyte maturation-inducing hormones. Development 145

Takeda N, Kyozuka K, Deguchi R (2006) Increase in intracellular cAMP is a prerequisite signal for initiation of physiological oocyte meiotic maturation in the hydrozoan Cytaeis uchidae. Dev Biol 298: 248–258

Vigneron S, Brioudes E, Burgess A, Labbe JC, Lorca T, Castro A (2009) Greatwall maintains mitosis through regulation of PP2A. EMBO J 28: 2786–2793

Von Stetina JR, Orr-Weaver TL (2011) Developmental control of oocyte maturation and egg activation in metazoan models. Cold Spring Harb Perspect Biol 3: a005553

Voronina E, Wessel GM (2003) The regulation of oocyte maturation. Curr Top Dev Biol 58: 53–110

Wang J, Liu XJ (2004) Progesterone inhibits protein kinase A (PKA) in Xenopus oocytes: demonstration of endogenous PKA activities using an expressed substrate. J Cell Sci 117: 5107–5116

Yamashita M (1988) Involvement of cAMP in initiating maturation of the brittle-star Amphipholis kochii oocytes: induction of oocyte maturation by inhibitors of cyclic nucleotide phosphodiesterase and activators of adenylate cyclase. Dev Biol 125: 109–114

Yi JH, Lefievre L, Gagnon C, Anctil M, Dube F (2002) Increase of cAMP upon release from prophase arrest in surf clam oocytes. J Cell Sci 115: 311–320

