## Supplementary data for "Cross-species analysis of ARPP19 phosphorylation during oocyte meiotic maturation charts the emergence of a new cAMP-dependent role in vertebrates"

#### Figure S1. ARPP19/ENSA protein sequences used in the study and their accession identifiers

##### A. Sequences used for the alignments in Figure 1

| | |
|---|---|
| *Amphimedon queenslandica* | XP_003389546.1 |
| *Arabidopsis thaliana* | NP_564967.1 |
| *Aspergillus niger* | XP_001392163.1 |
| *Branchiostoma lanceolatum* | CAH1254632.1 |
| *Caenorhabditis elegans* | NP_492609.1 |
| *Capitella teleta* | ELT98737.1 |
| *Ciona intestinalis* | XP_002129373.1 |
| *Choanoeca perplexa* | GGOP01002813.1 (TSA translation) |
| *Clytia hemisphaerica* | XLOC_041555 from *Clytia* genome assembly GCA_902728285.1 |
| *Drosophila melanogaster* | NP_001246745.1 |
| *Homo sapiens* (ARPP19) | KAI4057880.1 |
| *Hydra vulgaris* | XP_047140150.1 |
| *Lottia gigantea* | XP_009048785.1 |
| *Nematostella vectensis* | XP_032227676.1 |
| *Mnemiopsis leidyi* | ML210012a (from https://research.nhgri.nih.gov/mnemiopsis/) |
| *Patiria miniata* | XP_038072705.1 |
| *Saccoglossus kowalevskii* | XP_002732849.1 |
| *Sphaeroforma arctica* | EC804984. 1 (TSA translation) |
| *Xenopus laevis* (ARPP19) | NP_001086634.1 |

##### B. Additional sequences used for alignments used in Supp Fig. S2

| | |
|---|---|
| *Acropora digitifera* | XP_015773650.1 |
| *Actinia tenebrosa* | XP_031564726.1 |
| *Amphiprion ocellaris* | XP_023139216.1 |
| *Aplysina aerophoba* | HANI01464700.1 (TSA translation) |
| *Asterias rubens* | XP_033633809.1 |
| *Aurelia aurita* | GHAG01094477.1 (TSA translation) |
| *Bufo bufo* | XP_040269877.1 |
| *Beroe forskalii* | GHXY01160637.1 (TSA translation) |
| *Byssothecium circinans* | KAF1955593.1 |
| *Cassiopea andromeda* | GJJJ01041283.1 (TSA translation) |
| *Danio rerio* | Q7ZUT5 |
| *Dendronephthya gigantean* | XP_028399175.1 |
| *Dynamena pumila* | GHMC01017442.1 (TSA translation) |
| *Ephydatia muelleri* | AM760832.1 (cDNA translation) |
| *Exaiptasia diaphana* | XP_020907908.1 |
| *Haplosporangium sp. Z 27* | KAF9202459.1 |
| *Hartaetosiga balthica* | GGOO01015587.1 (TSA translation) |
| *Hormiphora californensis* | GGLO01016586.1 (TSA translation) |
| *Hydractinia symbiolongicarpus* | GAWH01050646.1 (TSA translation) |
| *Hymeraphia stellifera* | GKDX01076453.1 (TSA translation) |
| *Limulus polyphemus* | XP_013774917.1 |
| *Lupinus albus* | KAE9595631.1 |
| *Lytechinus variegatus* | XP_041465673 |
| *Mytilus galloprovincialis* | VDI17929.1 |
| *Paracentrotus lividus* | 27929.1 |
| *Podila clonocystis* | KAG0011881.1 |
| *Podocoryna carnea* | GCHV01001733.1 (TSA translation) |
| *Penicillium griseofulvum* | KXG51649.1 |
| *Physalia physalis* | GHBB01030377.1 (TSA translation) |
| *Quercus robur* | XP_050277191.1 |
| *Rana temporaria* | XP_040198643.1 |

| | |
|---|---|
| *Rhopilema esculentum* | GEMS01057539.1 (TSA translation) |
| *Salpingoeca rosetta* | XP_004988287 |
| *Saccharomyces cerevisiae* | KZV10868.1 |
| *Salmo salar* | ACN10327.1 |
| *Schizosaccharomyces pombe* | NP_593267.1 |
| *Stylophora pistillata* | PFX34499.1 |
| *Takifugu rubripes* | XP_003969615.1 |
| *Velella velella* | GHAZ01100014.1 (TSA translation) |

#### C. *Clytia hemisphaerica* ARPP19 sequences

gene identifier XLOC_041555 from genome assembly GCA_902728285.1[1]

**Nucleotide sequence of transcript**

```
GTTTTGTTCAGGAAGCGCTCGCACTCGCAGGCAAGAAGATTCAACCAAGGCCAAGGTTAAAGCCAAGAGTGAGGA
AATCAAAGAAAATCAACATCATTTTAATTTAAATCGAGAAGAAAAGCAACACAAGGGTATCGAAGACGATGATTT
AGATTGAGAATTTAACCTTACAAATTCATAAGGAGCGTTTGTTACTCAACGTTGAATCAACAAAACAACTTTGTC
TCGTCTGAGAAGCTGTCAAGCAAAAAGAAGTTTTTTAAGCATATTGTTGTTTTCTCTTGACTATTATCACCCAAA
TGTCTACAATGGCAACATTTAAAGGCCCACAAGAGTTGCCAGAAGCAGAACGCTTCAAACAGAAGTTCCCACAAG
GAGCACCAAGGCAGTCTGATTTTCTAAGAAAACGCTTACAAGGCAAAGGAGGGGCAAAGTATTTTGATTCTGGGG
ATTACAACATGGCCAAGAGCACTCATGGTAGGAATGTTCCAACTGGGAAAGCCATTCCAACGCCTGAAAATTTAC
CTCAAAGGAAAATCTCTTCTACTAAACAGAGTAACCTGATAGAAGGCGGCACGAGTCCCCCGCAGAGCGCTCTAC
CAGAGGAAAAAGCTGTGACTGAAGGTGAGCAGCATAAATCAAAACAACATATCACTGAACAAGAATAATGTTTAT
ATATACCCCCAATTAATATTTTTAGCCAGAAAAGTTGTAGTCACTTGACTTGAATTTATTAAGTTTATTAGGAAT
GGATCCATTTTTATTTTGTTCCATTTCTCACATTAACACTGTACATAGCTCATATCTGTAGATAGGAATTAGATC
TTCATTGGGTCAAGCTTCTGAATTCCTTTCTGAAGCTTCTTTTGGACCTATGATTGGGCTATAGTTTTATCTACT
TTTTAAAGACACTTTCCTCTAATACCCCCACCGTCAATATTTTGTTATTTAAGGGTTTCTTTCCCTACGATCTTA
TCTCTGTTGATATGAAACCTTTTTTACACAATCGTCAGCAATTTAATCTACCTTGATATACCAAATCCTTGCCCC
TTCCCCTCCCCCCTCAGGTTCTATATTATGCCACAGCACGTATAAATGAACTCTGGCTATGCCCATGGTTGCTGT
AGACTACACCTGTTATCTTCGTTTTATGTAGTAACTTTTTTGTTTATGTTTTATTTTTTGGTAAAACCGAAAA
```

**Predicted protein sequence**

```
MSTMATFKGPQELPEAERFKQKFPQGAPRQSDFLRKRLQGKGGAKYFDSGDYNMAKSTHGRNVPTGKAIPTPENL
PQRKISSTKQSNLIEGGTSPPQSALPEEKAVTEGEQHKSKQHITEQE
```

1. Leclere L, *et al.* The genome of the jellyfish Clytia hemisphaerica and the evolution of the cnidarian life-cycle. *Nature ecology & evolution* **3**, 801-810 (2019).

#### Figure S2. Alignments of ARPP19 protein sequences from various eukaryotic species.

##### A. Relationships between the main eukaryotic clades

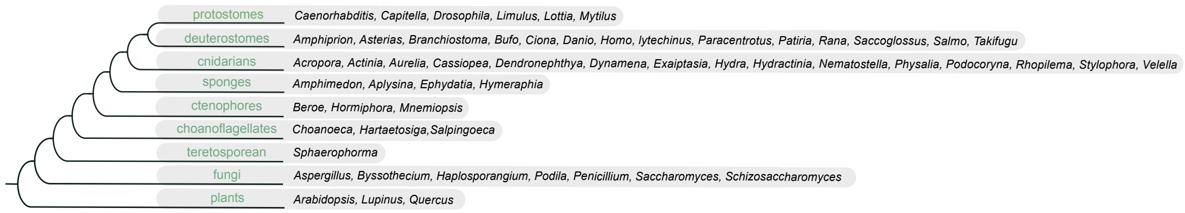

Phylogenetic tree indicating relationships between the main eukaryotic clades to which the species in B to H belong, adapted from[1]. All accession numbers are provided in Supp Fig. S1.

[1]Grau-Bove X, Torruella G, Donachie S, Suga H, Leonard G, Richards TA, Ruiz-Trillo I (2017) Dynamics of genomic innovation in the unicellular ancestry of animals. *eLife* 6

##### B. Alignment of an extended selection of ARPP19 protein sequences

```
Caenorhabditis      -------------------------------------MRGEAGELA------------VS
Drosophila          MSS------------------------AEENSN-------SPAT-----------TPQDT
Capitella           MSQE------------------------EQRIASAGSKASDEGSEKVDTGS--EKDGEFK
Mytilus             MSSE-----------------------GTIDRKEPEFVAPE-------------------
Lottia              --------------------------------------------MSMASETGNDVPTGDA
Limulus             MSS------------------------EEYSE---------------SP----IKVEKCE
Saccoglossus        ------------------------------------------------------------
Patiria             MSDNSVKAEKPQATEQEPKPEAQ-VTAEEETTQEPEVIEDKPTPLPLLGGKENQQAASFK
Branchiostoma       ----------------------------------------------MAT----DTARPPS
Ciona               --------------------------------------------MSVSPDVVPTMEPVNK
Takifugu            MSG------------------------DNEDP---------------QP----AEESPVD
Xenopus             MSG------------------------ENQET--------------KAQ----EESSALE
Homo                MSA------------------------EVPEAASAEEQKSEHNMLPWSL----QPSIPNS
Aurelia             --------------------------------------------------------MSAP
Clytia              ------------------------------------------------------------
Hydra               ------------------------------------------------------------
Dendronephthya      -----------------------------------------------------MADTSIP
Nematostella        -----------------------------------------------------------M
Stylophora          -----------------------------------------------------------M
Amphimedon          MANKEE-------------A-----------------------TVP----------EVKE
Ephydatia           MTTEGT-------------ETTILVTEPEENT-----HQDKPVIEPV--------TVSAE
Aplysina            MSAEEK-------------------------------------NPPA----------TEE
Hymeraphia          MSDEEK-------------PRA----EQEERE------KANGASLPA----------TKD
Mnemiopsis          MASC-----------QE----DQ-IP----SNTAGSDVNASNTSNEVIS--EVASDSVPV
Beroe               MASC-----------QE----DN-VH----SEPSPPVIENIPQQESEVT--VEAEDVPCV
Choanoeca           ------------------------------------------------------------
Hartaetosiga        ------------------------------------------------------------
Sphaeroforma        ------------------------------------------------------------
S.cerevisiae IGO2   -------------MSEDLSPT------------SSRVDL-----------------SNPH
Aspergillus         --------------------------------------------------------MNP-
Arabidopsis         -----MEDVKGKEIIDD-------------------------------------------
Lupinus             MSDTNVEDVKKQESLHDPDPKDDVGS-----DNNSEDDK-----------------KDLN
Quercus             MSGANNEDIKEQELTDN-TLNNQVDG-----D-SSVDD------------------LNKA

Caenorhabditis      SGEIATGALSPEKQQEQELMGKLAATGKLPARPASSFLQKKLQ-----QRKFFDSGDYAM
Drosophila          ETTEQANLTDLEKIEEEKLKSKYPSG-MRVPGGHSAFLQKRLQ----KGQKFFDSGDYQM
Capitella           ---VPLSKVDIEREQENKLKARYPGV---KSGGGSALLQKRIA----KPQKFFDSGDYAM
Mytilus             --PTTMTIKEKMMMEEAKLKAKYPNV---KPGGPSGLLAKRLQ----RGQKYFDSGDYNM
Lottia              DVFKQPQPKTVEQNEEARLRSKYPNL----KGGGSAILQKRLI----NKGKYFDSGDYNM
Limulus             ENEKKEELRKAELEEEAKLRAKYPQA---MRPGGSAFLQKRLH----KGQKYFDSGDYNM
Saccoglossus        MSIETSQKSQLEQMEEAKLKSKYPGM---QRPGGSDFLRKRLQ----KGPKYFDSGDYNM
Patiria             KPLDVKPQLSPEKMEEAKLSAKYGKL----KPGGSDFLRKRLN----KGVKYFDSGDYQM
Branchiostoma       MESGEAKELTPEQAEEAKLKAKYPALGHQQRPGGSDFLRKRLQ----KGPKYFDSGDYNM
Ciona               PYTETVPSISVEQQQEQILKAKYGNL---AKKGGSSLLQKRLAQ--KGGNKYFDSGDYNM
Takifugu            EKEIQDKVISPEKAEEAKLKARYPNLG--NKPGGSDLLRKRLQ----KGQKYFDSGDYNM
Xenopus             QKEIDDKVVSPEKSEEIKLKARYPNLG--PKPGGSDFLRKRLQ----KGQKYFDSGDYNM
Homo                LEEMEDKVTSPEKAEEAKLKARYPHLG--QKPGGSDFLRKRLQ----KGQKYFDSGDYNM
Aurelia             NAGLPPHLKAAELSEDQKYKNKFPGK---KAPANTDFLRKRLQ----KGVKYFDSGDYNM
Clytia              MSTMATFKGPQELPEAERFKQKFP-Q---GAPRQSDFLRKRLQG--KGGAKYFDSGDYNM
Hydra               --MEATLKIQPDSSEADHFKNKFPGK---GAPKQTDFLRKRLQG--KSGGKYFDSGDYNV
Dendronephthya      TVENPVEEKPKELSEEQKLKAKYPG----GLKKNFAQSRLQNR----GGVKYFDSGDYNM
Nematostella        SAE-EPATNATELSEEEKFKRKFPN----KKPMTTDVMRKRLQ----KGVKYFDSGDYMM
```

```
Stylophora           SEETGLEEKPSELTEEQKLALKFPG----KKPVATDILKKRMQ----RGVKYFDSGDYAM
Amphimedon           PSSEEKDTIDPIKKSEQELMSRYPKH---LQPKQKSVAQRKISS---GGVQYFDSGDYNA
Ephydatia            PGKATLDP--------QQADPKYGGL-----VKKAPFAQRKISE---GGRQYFDSGDYQT
Aplysina             DSSTEPDPSSDIAKAEAEYMAKFGKN-----QKKSNFAQRKMMM---EGKRYFDSGDHST
Hymeraphia           --AGHLDPKSADAVAQAEYQKKYGFV-----KKPPAFAQRKISQ---GGRQYFDSGDYNM
Mnemiopsis           NAPANAPLAQAIQEEEKKAAMKYGN----MKKKQSDVMRKRMMG---GGKQYFDSGDYNV
Beroe                KNSVPTPLAKAIADDEQKAAAKYGN----MKKKQTDVMRKRMMG---GGKQYFDSGDYNV
Choanoeca            ------MADAGLTPEQIMFKKKYGRL-----PPKKKVMKGPLGRVGGGERTFFDSADHAM
Hartaetosiga         -------MDPQMQAQEAVLKAKFGGM-----MGK--KKKGHLSQFKNKDRAYFDSADHAL
Sphaeroforma         ----MADAPSNLSAQDQEFLKKFGRL-----PPKNKLAQKRMN--RGGERKYFDSGDYAL
S.cerevisiae IGO2    GFTKEGVDLSKLSPQELKLYKMYGKL-----PSKKDLLRHKM-----QDRQYFDSGDYAL
Aspergillus          -HQQNKIDTSKMSPDEQRLFRLYGKM-----PTKKDLLQNKL-----KERKYFDSGDYAL
Arabidopsis          -APIDNKVSDEMESEENAIKKKYGGL-----LPKKIPLISKD-----HERAFFDSADWAL
Lupinus              EHKNDGNPMPSSHQEEEVIKKKYGGL-----IPKKPPLISKD-----HERAYFDSADWAL
Quercus              VDRNDENPMPSPQQEEEIVKKKYGGL-----LPRKHPLISKD-----HERAFFDSADWAL

Caenorhabditis       DKSKAGTGLGSKPHPLAGGPPPAAPPVVAQRSPAPAATTPSPSASPISQQTNRPSSDRN
Drosophila           AKQKGGGVK------------------------QVF---------------------AN
Capitella            AKAKKQPVT------------------VGVKLPYNA---------------------PA
Mytilus              AKAKMNNPK----TPLAP--------------TEKM---------------------IL
Lottia               AKAKLNNPK----KPLNQ--------------SEKF---------------------LI
Limulus              AKAKSQKAR----VP------------------AAP---------------------IV
Saccoglossus         AKQTGQLLR----KPNNA--------GNGEKMPPPP---------------------AR
Patiria              EQQSGKMKL----RPNSGKMPGLAGARVGGAMPAPL---------------------PR
Branchiostoma        ALAKTKKVP----IR-----------GNGARL-QQP---------------------NP
Ciona                ARAKVGGKQ----KI------------------PIE---------------------VK
Takifugu             AKAKIKNKQ----LP-----------------TAAP---------------------EK
Xenopus              AKAKMKNKQ----LP-----------------TAAS---------------------DK
Homo                 AKAKMKNKQ----LP-----------------TAAP---------------------DK
Aurelia              AKSTNRTGS--------------------------------------------------
Clytia               AKSTHGR----------------------------------------------------
Hydra                AKSTKKKDA--------------------------------------------------
Dendronephthya       AKTTGNKSR--------------------------------------------------
Nematostella         AKSRDKNPR-------------------GPVN-------------------------PA
Stylophora           AKSEKKAGPAAA--------------AIGNRG-------------------------KL
Amphimedon           AAHKKEKQKMLDLP---------------------------------------------
Ephydatia            ATG-KDRTAIASHPHAMAPEALQAAKVPPKPVAG-----------------------KA
Aplysina             AGSKSRKEEVASRPHLPLMNQQKAKRPSVTS----------------------------
Hymeraphia           HASKKDRQEIASHPHVMNPHLAQLRAKAVISKSS-----------------------PS
Mnemiopsis           GKSRPTAQMM-------------------------------------------------
Beroe                GKQC--TPTV-------------------------------------------------
Choanoeca            SKAGISVD---------------------------------------------------
Hartaetosiga         STAGVKVD---------------------------------------------------
Sphaeroforma         SQAGKSQNK--------------------------------------------------
S.cerevisiaeIGO2     KKAGVIKS-------------------------------------------------DD
Aspergillus          SKAGKASD-------------------------------------------------VG
Arabidopsis          GKQKGQKPKG-----------------------------------------------PL
Lupinus              GKQGAQKPKG-----------------------------------------------PL
Quercus              GKQGAQKPKG-----------------------------------------------PL

Caenorhabditis       SDDDNLQIPRPDTVP-QRKASI--I-NPSVHCKLSPAPHVQHHDAASPNATSE
Drosophila           KVTTGEAIPTPETVP-ARKTSI--I-QPCNKFPATS
Capitella            LAPTGDEHPTPDTVP-ARKASL--V-QNKLLTRLT
Mytilus              QESIGDAIPTPESLP-PRKPSI--I-QSRLAEGELS
Lottia               EESVGEAIPTPENLP-ARKASL----PSKLVNPLLEQAKQ
Limulus              RESTGDAIPTPDVLP-TRKTSL--V-QSKLATELS
Saccoglossus         VPLTGDSIPTPEDLP-QRKSSI--T-ASKLAAM
Patiria              GPATGKTIPTPASIP-HRKQST--E-ISKLAV
Branchiostoma        KEVTGDTIPTPDNVP-ARKPSL--Q-TSKLVS--DQPQITKPEAVVGLKKGDGDAGEEKKDEEK--
Ciona                KEVTGDHMPTPDELPVRRKAS---Q--SSLVI--PQLN
Takifugu             AEITGDHIPTPQDLP-QRKPSL--V-ASKLAG
Xenopus              TEVTGDHIPTPQDLP-QRKPSL--V-ASKLAG
Homo                 TEVTGDHIPTPQDLP-QRKPSL--V-ASKLAG
Aurelia              NPLLGKAIPTPEGVT-QRKHST--ASKSKLVDG----AEKSPSPNDEKAFFEKMDEKENNAEEEEKQ
Clytia               NVPTGKAIPTPENLP-QRKISS--TKQSNLIEG--GTSPPQSALPEEKAVTEGEQHKSKQHITEQE
Hydra                TIITGKAIPTPEVIP-HRKPSS--S-QSNLLGV-------PDHTLEEKKVEVENEEAKNAE
Dendronephthya       LPGLGAAIPKPEDIP-HRKQSV----PSKLASP-TSVSVQD
Nematostella         VLAVGKGIPTPDKIP-HRKTSV----PMTEHPV-TQTVPTHPHQPHHTNVEVEN
Stylophora           PMGVGKAIPTPDNIP-HRKTSV----PSKLVEG-QEQPPVAVTR
Amphimedon           ----GRPAINPAMAR------LQGRPPSKLTAPKSGLASVARSEQ
Ephydatia            MAERGFPVPTRRAPA--------ERGVSKLASPSKRVSVPSPLAQ
Aplysina             KLVD
Hymeraphia           KLASGSP------------------PSKLAAS----PRQSPLAQ
Mnemiopsis           PQAINPEQPAPEKLR-NRKASIPNKTFSRLQS
Beroe                PQTISPEQPAIEKLR-NRKISIPNKHLTRLS
Choanoeca            ---NGKVIPNPENIS-AK----PHAKPSGMTKGAVVDEDAPAADESAAAPATDTPA—AEPEEKPAQ
Hartaetosiga         -RTPGNIIATPENIP-KK----THAKPSELVDHTKEA---LAEDE
Sphaeroforma         ---VGRALPTPETIP-HHSDNVQPKKQSHLSEKSGSLTNPDK---------------PSDSEKKA
S.cerevisiae IGO2    VIVNNSSNNLPVTNP-SGLRESIIRRRMSSSSGGDSISRQGS------ISSGP-----PPRSPNK
Aspergillus          VTNIGSQHPVPENIP-HLTATSPGANNPAAASNGGSISAQGGQQIPGSISGHPGSIGFQSRSPVK—
Arabidopsis          EALRPKLQPTPQQQPRARRMAYSSGETEDTEIDNNE-APDDQACASAVDSTNLKDDGGAKDNIKS
Lupinus              EALRPKLQPT-QQHARSRRSAYAPADDS-EVDGCNNHASSEDQSAAEDVANDTAQDQSSHQ
```

```
Quercus             EALRPKLQPTPHQQVRSRRSAYAPADEGGEVDGSNNHSISSEDQSCMLDAGEHNDTTSDGDHNDKTGSNDQKCHE

Branchiostoma       PEEQSTENQENQEHH

Aspergillus         EASYLQRETSADETEEAEKKEDDSVSPPPARGGVPIRQ
```

Conserved amino acids of the Gwl phosphorylation motif are highlighted in green, of the PKA site in yellow, and of other conserved sequences discussed in the text in blue and magenta.
Text colours indicate species belonging to different taxa as following: Red=Deuterostomes; Black=Protostomes; Dark blue=Anthozoan cnidarians; Light blue=Medusazoans; Purple=Sponges; Red=Ctenophores; Orange=Choanoflagellates. Pale orange=Teretosporeans; Grey=Fungi; Green=Plants.

#### C. Alignment of *Clytia* and *Xenopus* ARPP19 protein sequences with those from a selection of other deuterostomes

```
Clytia          MST------------------------------------------------------------   3
Xenopus         MSGENQET-------------------------------------K----------AQEES--  14
Lytechinus      MS-SAEETQSTPAEAPGTEVQAPAEE-ESMDTTPAAPAATDTTESAPSQTTTPPPETTPSQPV  61
Paracentrotus   METPAETTQTTPAEATAAEVEAPAQEEESMDTTPAPAP-------APTTTESPPVNVAPAQPV  56
Asterias        MSE--DTVKTDSPTVPE------SEPEVAMETTSQAESATVEEKSSPLPSVE-DEDKPTPLPT  54
Patiria         MSDNSVKAEKPQATEQEPKPEAQVTAEEETTQEPEVIEDKPTPLPLLGG------KENQQA--  55
Saccoglossus    M-SIETS--------------------------------------------------------   6
Branchiostoma   M-ATDTA--------------------------------------------------------   6
Ciona           MSVSP---------------------------------------------------------D   6
Takifugu        MSG-----DNEDP-------------------------------------------QPAEE--  13
Salmo           MSEEIEG-------------------------------------------------TRPAT--  12
Danio           MS------------------------------------------------------SEVET--   7
Amphiprion      MSEEVEG----------------------------------------------------TR--   9
Bufo            MSADNQET-------------------------------------K----------AHEEN--  14
Rana            MSAENLES-------------------------------------K----------TPEEA--  14
Homo            MSAEVPEAAS---------------AE-EQKSEH-------NMLPW----------SLQPS--  28

Clytia          ---------------------------------MATFKGPQ-ELPEAERFKQKFP-QG---AP  28
Xenopus         -------------------SA------LEQ-KEIDDK-VVSPEKSEEIKLKARYPNLG--PKP  48
Lytechinus      MTESNQNIPKTDTGKFIRPAQPVIKPPVPKTIQTDRPSKQSIEQMEEAKLKAKFGGLS---KP  120
Paracentrotus   MTESSQNIPKSDSGKFIRPANPLIKPPVPKTIQ-DRPSKQSIEQMEEAKLKAKFGGLS---KP  114
Asterias        VTEDVKPAPLPTATLTATEKP-----PFKK-PMDIKKPQLSPEKAEEAKLNAKYGKV----KP  107
Patiria         -------------------AS------FKK-PLDVK-PQLSPEKMEEAKLSAKYGKL----KP  87
Saccoglossus    -------------------------------------QKSQLEQMEEAKLKSKYPGM---QRP  29
Branchiostoma   -------------------RP------PSM-ESGEA-KELTPEQAEEAKLKAKYPALGHQQRP  42
Ciona           ------------------VVP-TMEPVNKP-YTETV-PSISVEQQQEQILKAKYGNL---AKK  45
Takifugu        -------------------SP------VDE-KEIQD-KVISPEKAEEAKLKARYPNLG--NKP  47
Salmo           -------------------GE------ATR-EEMDD-TVLSPEKAEEVKLKARFPHLG--AKP  46
Danio           -------------------ST------EEQ-QEMQD-TVVSPEKAEEIKLKARYPHLG--ARP  41
Amphiprion      -------------------TL------DEQ-QEMED-KVISPEKAEEAKLKARYPNLA--PKH  43
Bufo            -------------------SS------DEQ-KEMED-KVISPEKAEEAKLKARYPHLG--PKP  48
Rana            -------------------TT------DEQ-KEVDD-KVVSPEKAEEAKLKARYPHLG--PKP  48
Homo            -------------------IP------NSL-EEMED-KVTSPEKAEEAKLKARYPHLG--QKP  62

Clytia          -RQSDFLRKRLQGKGGAKYFDSGDYNMAKSTHGR----------------------------N  62
Xenopus         -GGSDFLRKRLQ-KGQ-KYFDSGDYNMAKAKMKNK-QLP-----------------TAASDKT  90
Lytechinus      -GGSQFLQKRLN-KG-MKYFDSGDYNMAKQSGKLG-RRP-------LSGKPG-GIP--SA--P  168
Paracentrotus   -GGSQFLQKRLN-KGQMKYFDSGDYNMAKQSGKLK-MRP-------LSGKPG-GIA--PVPVP  165
Asterias        -GGSDFLRKKLN-KGV-KYFDSGDYNMEQQSGKLR-NR---GKPPGLAGPPG-VRLPVQSPLG  162
Patiria         -GGSDFLRKRLN-KGV-KYFDSGDYQMEQQSGKMK-LRPNSGKPMGLAGARVGGAMPAPLPRG  146
Saccoglossus    -GGSDFLRKRLQ-KGP-KYFDSGDYNMAKQTGQLL-RKPNNA--------GNGEKMPPPPARV  80
Branchiostoma   -GGSDFLRKRLQ-KGP-KYFDSGDYNMALAKTKKV-PIRG-----------NGAR-LQQPNPK  89
Ciona           -GGSDLLQKRLAQKGGNKYFDSGDYNMARAKVGGK-QKI------------------PIEVKK  88
Takifugu        -GGSDLLRKRLQ-KGQ-KYFDSGDYNMAKAKIKNK-QLP-----------------TAAPEKA  89
Salmo           -GGSDLLRKRLQ-KGQ-KFFDSGDYNMAKAKVKNKQQLP----------------AVTPAEKA  90
Danio           -GGSDLLRKRLQ-KGQQKYFDSGDYNMAKAKMKNK-QLP-----------------AAAAEKT  84
Amphiprion      -GGSDFLRKRLQ-KGQ-KYFDSGDYNMAKAKMKNK-QLP-----------------SAPTEKT  85
Bufo            -GGSDFLRKRLQ-KGQ-KYFDSGDYNMAKAKIKNK-QLP-----------------AAAPDKT  90
Rana            -GGSDFLRKRLT-KGQ-KYFDSGDYNMAKAKMKNK-QLP----------------TTTAADKA  91
Homo            -GGSDFLRKRLQ-KGQ-KYFDSGDYNMAKAKMKNK-QLP-----------------TAAPDKT  104

Clytia          VPTGKAIPTPENLPQ-RKISSTKQSNLIEGG-TSPPQSALP---------------EEKAVTE  108
Xenopus         EVTGDHIPTPQDLPQ-RKPSL-VASKLAG                                   117
Lytechinus      KPTGEAIPTPDSIHH-RKQSS-EISKLAV                                   195
Paracentrotus   KPTGEAIPTPDSIHH-RKQSS-EISKLAV                                   192
Asterias        QATGKTIPTPASIPH-RKQST-EISKLAV                                   189
```

```
Patiria          PATGKTIPTPASIPH-RKQST-EISKLAV                                       173
Saccoglossus     PLTGDSIPTPEDLPQ-RKSSI-TASKLAAM                                      108
Branchiostoma    EVTGDTIPTPDNVPA-RKPSL-QTSKLAS----DQPQITKPEAVVGLKKGDGDAGEEKKDEEK     146
Ciona            EVTGDHMPTPDELPRVRKAS---QSSLVI------PQLN                             118
Takifugu         EITGDHIPTPQDLPQ-RKPSL-VASKLAG                                       116
Salmo            EITGDHIPTPQDIPQ-RKPSI-MASKLAG                                       117
Danio            EITGDHIPTPQDLPQ-RKQSL-EASKLAF                                       111
Amphiprion       EITGDHIPTPQDLPQ-RKTSI-VASKLAG                                       112
Bufo             EVTGDHIPTPQDLPQ-RKPSL-VASKLAG                                       117
Rana             EVTGDHIPTPQDLPQ-RKPSL-VASKLAG                                       118
Homo             EVTGDHIPTPQDLPQ-RKPSL-VASKLAG                                       131

Clytia           GEQHKSKQHITEQE      122
Branchiostoma    PEEQSTENQENQEHHS    105
```

Conserved amino acids of the Gwl phosphorylation motif are highlighted in green, of the PKA site in yellow, and of other conserved sequences discussed in the text in blue and magenta.
Text colours indicate species belonging to different taxa as following: Dark blue=Fish; Light blue=Tunicates; Purple=Echinoderms; Orange=Hemichordate. Grey=Cephalochordate; Green=Amphibians; Red=Mammals.

#### D. Alignment of *Clytia* and *Xenopus* ARPP19 protein sequences with those from a selection of other cnidarians

```
Clytia           MST------------------MATFKGPQ-ELPEAERFKQKFP-QG-AP-RQSDFLRKRLQGK     41
Xenopus          MSGENQETKAQEESSALEQKEIDDK-VVSPEKSEEIKLKARYPNLGPKP-GGSDFLRKRLQ-K     60
Physalia         ME----------------------SGKLKEMSETE-RFKAKYGAHE-AP-RQSAFLRKRLQG-     37
Velella          MEA-----------------ETFKVPHKVQMNEAE-TFKKKFPGHG-AP-KQSDFLRKKLHGK     43
Hydra            MEA------------------TLKIQP---DSSEADHFKNKFPGKG-AP-KQTDFLRKRLQGK     40
Dynamena         MD-------------------TFKAPH---EMSEAELFKKKFPGQG-VP-RQSDFLRKRMQGK     39
Podocoryna       MA-------------------TFKAPH---EISEADRFKQKFPGQG-VP-KQSDFLRKRLQGK     39
Hydractinia      MA-------------------TFKAPQ---EISEADRFKQKFPGHG-VP-KQSDFLRKRLQGK     39
Aurelia          MSA--------------PNAGLPPHLKAA-ELSEDQKYKNKFPGKK-AP-ANTDFLRKRLQ-K     45
Cassiopea        MTEM----------N-APKPAIPPHLKAS-ELSEEQKFRNKYP--GKKP-ASTDFLRKRIQ-K     47
Rhopilema        MTEM----------N-APKLGLPPHLAAS-EVSEEQKFKNKYP--GKKP-ASTDFLRKRIQ-K     47
Dendronephthya   MADT----------S-IPTVENPVEEKPK-ELSEEQKLAKY-----KPGGLKNFAQSRLQNR     46
Acropora         MS-----------------DGTNLEEKPA-ELSEDQKFALKYP--GKKP-VATDILKKR-MQR     41
Stylophora       MS-----------------EETGLEEKPS-ELTEEQKLALKFP--GKKP-VATDILKKR-MQR     41
Nematostella     MSA-----------------EE-PATNAT-ELSEEEKFKRKFP--NKKP-MTTDVMRKRLQ-K     40
Exaiptasia       MA-----------------TES-EPKNLS-EMSEEEKFRAKFP--GRKP-TTTDVMRKRLQ-K     40
Actinia          MSGEN------------------EPMDKP-QLTEEEKFKAKFP--NRKP-NTTDLMRKRLQ-K     40

Clytia           GGAKYFDSGDYNMAKSTHG--RN---------------VPTGKAIPTPENLPQRKISSTKQSN     87
Xenopus          GQ-KYFDSGDYNMAKAKMKNKQLPTA-------ASDKTEVTGDHIPTPQDLPQRKPSLV-ASK     114
Physalia         G-NKYFDSGDYNVAKSNKRKDPT---------------MVIGKAIPTVDKLPQRKPSSSKLSN     94
Velella          GGNKYFDSGDYMMAKSTKKKDAT---------------IVTGKAIPTPEALPQRKPSATKQSN     91
Hydra            GGGKYFDSGDYNVAKSTKKKDAT---------------IITGKAIPTPEVIPHRKPSSS-QSN     88
Dynamena         GGAKYFDSGDYNMAKSK-TKNAN---------------VVTGKAIPTPDTLPQRKTSSTKQSN     86
Podocoryna       GGAKYFDSGDYNVAKSTKKKDAT---------------IVIGKAIPTPDNLPQRKPSCSKQST     87
Hydractinia      GGAKYFDSGDYNVAKSTKKKDAT---------------IVIGKAIPTPDNLPQRKPSCSKQSS     87
Aurelia          G-VKYFDSGDYNMAKSTNRTGSN---------------PLLGKAIPTPEGVTQRKHSTASKSK     92
Cassiopea        G-VKYFDSGDYNMAKSNSRTGPN---------------PLLGKAIPTAEAVTQRKHSTACKSK     94
Rhopilema        G-VKYFDSGDYNMAKSTNRTGPN---------------PLLGKAIPTPEAVTQRKHSTACKSK     94
Dendronephthya   GGVKYFDSGDYNMAKTTGNKS---------------RLPGLGAAIPKPEDIPHRKQSVP--SK     92
Acropora         G-VKYFDSGDYAMAKSEKKGGPAA--AINRGKPMPCMPLGVGKAIPTPENIPHRKTSVP--TE     99
Stylophora       G-VKYFDSGDYAMAKSEKKAGPAA--AAIGN--RGKLPMGVGKAIPTPDNIPHRKTSVP--SK     97
Nematostella     G-VKYFDSGDYMMAKSRDKNP-----RGP----VNPAVLAVGKGIPTPDKIPHRKTSVP--MT     91
Exaiptasia       GGVKYFDSGDYAMAKSSEKKGPIP---HR----PNLDVLGVGRGIPTPDKIPHRKSSVP--IK     94
Actinia          GGVKYFDSGDYAMAKSTEKKGPIA--KPR----PNVDVLGVGRGIPTPDKIPHRKTSVP--IQ     95

Clytia           LIEGGTSPPQSALPEEKAVTEGE------Q----HKSKQHITEQE          122
Xenopus          LAG                                                    117
Physalia         LLEGDNPPEKTNNSTEEEVKDEKS-----ELPDEKNAKLDLADGK          124
Velella          LLSCEERDPNPAESSNDNSEVEK------IEPD-EKCK--LAG---------N  126
Hydra            LLGV----PDHTLEEKKVEVENE------EAKNAE                    113
Dynamena         LLEGTSPPPMNATAEERASVQEV------ASSDEQKTKQLITESDTTTTNTSN  133
Podocoryna       LLEGDAPHPHPSDEKETTTAAEQ------A----VMTE                 115
Hydractinia      LLEGDTPHPHPSDEKETPTVDSD------E----QKSNQAVMD            120
Aurelia          LVDGAEKSPSP--NDEKAFFEKMDEKENNAEEEEKQSHQMWIQLNAAHSNISNNNV 146
Cassiopea        LADGEEHSP--SAVTEPVFFADAEGVQTDENQDQTESR                 130
Rhopilema        LADGEEQLSPPCCGDEKVFFENVEQKDAQTENEEKN                   130
Dendronephthya   LASPTSVSVQD-----------------------                     103
```

```
Acropora           YHSRTRSDEFNLVMDQ------------------           115
Stylophora         LVEGQEQPPVAVTR--------------------           111
Nematostella       EHPVTQTVPTHPHQPHHTNVEVEN----------           115
Exaiptasia         EHEHQDTVH--HTHTPVEN---------------           111
Actinia            EHGVQSSTH--PEHPHTS----------------           111
```

Conserved amino acids of the Gwl phosphorylation motif are highlighted in green, of the PKA site in yellow, and of other conserved sequences discussed in the text in blue and magenta.
Text colours indicate species belonging to different taxa as following: Light blue=Hydrozoans; Green= Scyphozoans; Purple=Octocoralian; Dark blue=Hexacoralians.

#### E. Alignment of *Clytia* and *Xenopus* ARPP19 protein sequences with those from a selection of sponges

```
Clytia             MST-------------------------------------MATFKGPQ-ELPEAERFKQKFPQ    25
Xenopus            MSGENQET-------------------KAQEESSALEQKEIDDK-VVSPEKSEEIKLKARYPN    43
Amphimedon         MANKEEA------------------TVP------EVKEPSSEEKDTIDPIKKSEQELMSRYPK    39
Ephydatia          MTTEGTETTILVTEPEENTHQDKPVIEPV----TVSAEPGKATLDP--------QQADPKY--    49
Aplysina           MSAEEK-------------------NPPA------TEEDSSTEPDPSSDIAKAEAEYMAKF--    38
Hymeraphia         MSDEEKPRA----EQEERE-KANGASLPA------TKD--AGHLDPKSADAVAQAEYQKKY--    48

Clytia             -G-APRQSDFL-RKRLQGKGGAKYFDSGDYNMAKSTHGR------------------------    61
Xenopus            LGPKPGGSDFL-RKRLQ-K-GQKYFDSGDYNMAKAKMKNKQ---LP-----------------    83
Amphimedon         H-LQPKQKSVAQRK-ISS-GGVQYFDSGDYNAAAHKKEKQKMLDLP-----------------    82
Ephydatia          -GGLVKKAPFAQRK-ISE-GGRQYFDSGDYQTATG-KDRTAIASHPHAMAPEALQAAKVPPKP    108
Aplysina           -GKNQKKSNFAQRKMMME--GKRYFDSGDHSTAGSKSRKEEVASRPHLPLMNQQKAKRPSVT-    98
Hymeraphia         -GFVKKPPAFAQRK-ISQ-GGRQYFDSGDYNMHASKKDRQEIASHPHVMNPHLAQLRAKAVIS    109

Clytia             -------NVPTGKAIPTPENLPQRKISS--TKQSNLIEG--GTSPPQSALPEEKAVTEGEQHK    113
Xenopus            -TAASDKTEVTGDHIPTPQDLPQRKPSL--V-ASKLAG                             117
Amphimedon         -----------GRPAINPAMAR-----LQGRPPSKLTAPKSGLASVARSEQ                117
Ephydatia          VAG--KAMAERGFPVPTRRAPA-------ERGVSKLASPSKRVSVPSPLAQ                150
Aplysina           ---------------------------------SKLVD                             103
Hymeraphia         KSS--PSKLASGSP------------------PSKLAAS----PRQSPLAQ                136

Clytia             SKQHITEQE 122
```

Conserved amino acids of the Gwl phosphorylation motif are highlighted in green, of the PKA site in yellow, and of other conserved sequences discussed in the text in blue and magenta.

#### F. Alignment of *Clytia* and *Xenopus* ARPP19 protein sequences with those from a selection of ctenophores

```
Clytia             MST-------------------------------------MATFKGPQ-ELPEAERFKQKFP-    25
Xenopus            MSGENQET-----------------KAQ--EESSALEQKEIDDK-VVSPEKSEEIKLKARYPN    43
Mnemiopsis         MASCQEDQIPSNTAGSDVNASNTSNEVISEVASDSVPVNAPANAPLAQAIQEEEKKAAMKYGN    63
Beroe              MASCQEDNVHSEPSPPVIENIPQQESEVTVEAEDVPCVKNSVPTPLAKAIADDEQKAAAKYGN    63
Hormiphora         MASCQDTVTADTPAESAVPVIAA--------EAVAPDSSAPSPDTLAKAIEEDQKKAVSRYGN    55

Clytia             QG-APRQSDFLRKRLQGKGGAKYFDSGDYNMAKSTHG--RN--------VPTGKAIPTPENLP    76
Xenopus            LGPKPGGSDFLRKRLQ-KG-QKYFDSGDYNMAKAKMKNKQLPTAASDKTEVTGDHIPTPQDLP    104
Mnemiopsis         --MKKKQSDVMRKRMMG-GGKQYFDSGDYNVGKSRPTAQMM-------PQAINPEQPAPEKLR    116
Beroe              --MKKKQTDVMRKRMMG-GGKQYFDSGDYNVGKQC--TPTV-------PQTISPEQPAIEKLR    114
Hormiphora         --IKKKQTDIMRTRMMG-GGKQYFDSGDHALGRNRPNAPTI-------PSTLEPA--SVDILR    106

Clytia             QRKISS-TK-QSNLIEGGTSPPQSALPEEKAVTEGEQHKSKQHITEQE           122
Xenopus            QRKPSL---VASKLAG                                           117
Mnemiopsis         NRKASIPNKTFSRLQS                                           132
Beroe              NRKISIPNKHLTRLS                                            129
Hormiphora         NRKPSIPNKIVTRLS                                            121
```

Conserved amino acids of the Gwl phosphorylation motif are highlighted in green, of the PKA site in yellow, and of other conserved sequences discussed in the text in blue and magenta.

#### G. Alignment of *Clytia* and *Xenopus* ARPP19 protein sequences with those from a selection of sequences from Choanoflagellates

```
Clytia           MST------------------MATFKGPQ-ELPEAERFKQKFP-QG-APRQSDFLRKRLQG--    40
Xenopus          MSGENQETKAQEESSALEQKEIDDK-VVSPEKSEEIKLKARYPNLGPKPGGSDFLRKRLQ---    59
Choanoeca        MA-------------------------DAGLTPEQIMFKKKYGRLPPK---KKVMKGPL-GRV    34
Hartaetosiga     MDP-------------------------QMQAQEAVLKAKFGGMMG-K---KK--KGHL-SQF    31
Salpingoeca      MDP-------------------------TIQAQEE-KMRKFGRAPR-------PRKGPL-GKL    29

Clytia           --KGGAKYFDSGDYNMAKSTHG--RN--------VPTGKAIPTPENLPQRKISSTKQSNLIEG    91
Xenopus          --KG-QKYFDSGDYNMAKAKMKNKQLPTAASDKTEVTGDHIPTPQDLPQRKPSLVA-SKLAG    117
Choanoeca        GG-GERTFFDSADHAMSKAGISVD------------NGKVIPNPENIS-AKPHAKP-SGMTKG    82
Hartaetosiga     KNK-DRAYFDSADHALSTAGVKVD----------RTPGNIIATPENIP-KKTHAKP-SELVDH    81
Salpingoeca      SGR-ERKYFDSADHALSSAGVKT------------QTGKVIATAQNIP-RRTQSKP-TGMVDH    77

Clytia           GTSPPQSALPEEKAVTEGEQHKSKQHITEQE        122
Choanoeca        AVVDEDAPAADESAAAPATDTPAAEPEEKPAQ       114
Hartaetosiga     TKEA---LAEDE                           90
Salpingoeca      TQEVIETDGKSGDEQETTK                    96
```

Conserved amino acids of the Gwl phosphorylation motif are highlighted in green, of the PKA site in yellow, and of other conserved sequences discussed in the text in blue and magenta.

#### H. Fungi

```
Clytia           MST---------------------------------MATFKGPQ-ELPEAERFKQKFP-QG-A    27
Xenopus          MSGENQET-------------KAQ--EESSALEQKEIDDK-VVSPEKSEEIKLKARYPNLGPK    47
Aspergillus      MNP--------------------------------HQQNKIDTSKMSPDEQRLFRLYGKM---    28
S.cerevisiae     MSEDLSPTSSRVDL----------------SNPHGFTKEGVDLSKLSPQELKLYKMYGKL---    44
Podila           MASPL-------------------------SVA--TQPAFSPAQTLTEQEQKLQRMYGKL---    33
Haplosporangium  MSSPLNPS------------------HNSGGTA--TEHNFNPTPTLTEQEQKLQRLYGNR--L    41
S.pombe          MV-----------RTRKWMLS---TTIIAMSSS--NSEQKVDVAKLSPEEQKLFRLYGRL---    44
Byssothecium     MNP--------------------------------HQANKVDISKMSPEEAKLFRLYGKL---    28
Penicillium      MNP--------------------------------HQQNKVDINSLSPEEQRLLRLYGKM---    28

Clytia           PRQSDFLRKRLQGKGGAKYFDSGDYNMAKSTHG--RN----------VPTGKAIPTPENLP-Q    77
Xenopus          PGGSDFLRKRL-QKG-QKYFDSGDYNMAKAKMKNKQ--LPTAASDKTEVTGDHIPTPQDLP-Q    105
Aspergillus      PTKKDLLQNKL--K-ERKYFDSGDYALSKAGKASD---------VGVTNIGSQHPVPENIP-H    78
S.cerevisiae     PSKKDLLRHKM---QDRQYFDSGDYALKKAGVIKS---------DDVIVNNSSNNLPVTNP-S    94
Podila           PQPKDLLGKLR--SPDRKYFDSGDYALSKAGKIN-------------IPIGLKHPSPEAIP-H    80
Haplosporangium  PTAKGLLGQKL--K-ERKYFDSGDYALSKAGKTN-------------APVGSQHPQPENIP-H    87
S.pombe          PQRKDLLVQKL--QQGRKYFDSGDYALNKAGKASDSGI---------TCIGKEIPSPDTIP-H    95
Byssothecium     PNKKDLLQNKL--K-ERKYFDSGDYALSKAGKASDIGV---------TSIGREHPVPEKIP-H    78
Penicillium      PTKKDVLQNKL--K-ERKYFDSGDYALSKAGKASDVGV---------TNIGSRHPVPENIP-H    78

Clytia           RKISS--TKQSNLIEGGTSPPQSALPEEKAVTEGE------Q----HKSKQHITEQE 122
Xenopus          RKPSL--V-ASKLAG                                                   117
Aspergillus      LTATSPGANNPAAASN-GGSISAQGGQQIPGSISGHPGSIGFQSRSPVKEASY-LQRETSADE    139
S.cerevisiae     GLRESIIRRRMSSSSGGDSISRQGS------ISSGP-----PPRSPNK                  125
Podila           TNPHQSSSGS----------------------------------ISPIRESPLLHQTEFSQTR    109
Haplosporangium  STPAPSTQLI---YH--------------------------HHGSPPTKESSLIHESESAPAS    121
S.pombe          RVVSAGSPNKEPSLHTKRPSESSPSG---------A------SSR-----------RE-SVT-    130
Byssothecium     IAPPTQN------------------GQQQNGHDKGL-------SGSPVKEH-TFLHRETSLNR    115
Penicillium      LTSTSPGVNNSA---N-NGSVSSQGGQQIPGSISGHPGSIGFQSRSPIKDGGSFLHRGSSLSE    137

Aspergillus      TE--------EAEKKEDDSVSPPPARGGVPIRQ      164
Podila           --AQ                                   111
Haplosporangium  --HHHTIAAQPNV                          132
S.pombe          ---RHDLESNEN                           139
Byssothecium     ETSAEELENEEPKEENKA                     133
Penicillium      GEASSGLASGSVQGEQELSVSPPAAREGVPIRK      170
```

Conserved amino acids of the Gwl phosphorylation motif are highlighted in green, of the PKA site in yellow, and of other conserved sequences discussed in the text in blue and magenta.

### Supplementary Fig. S3
### Thiophosphorylation of ClyARPP19 at S49 and XeARPP19 at S67 by Gwl.

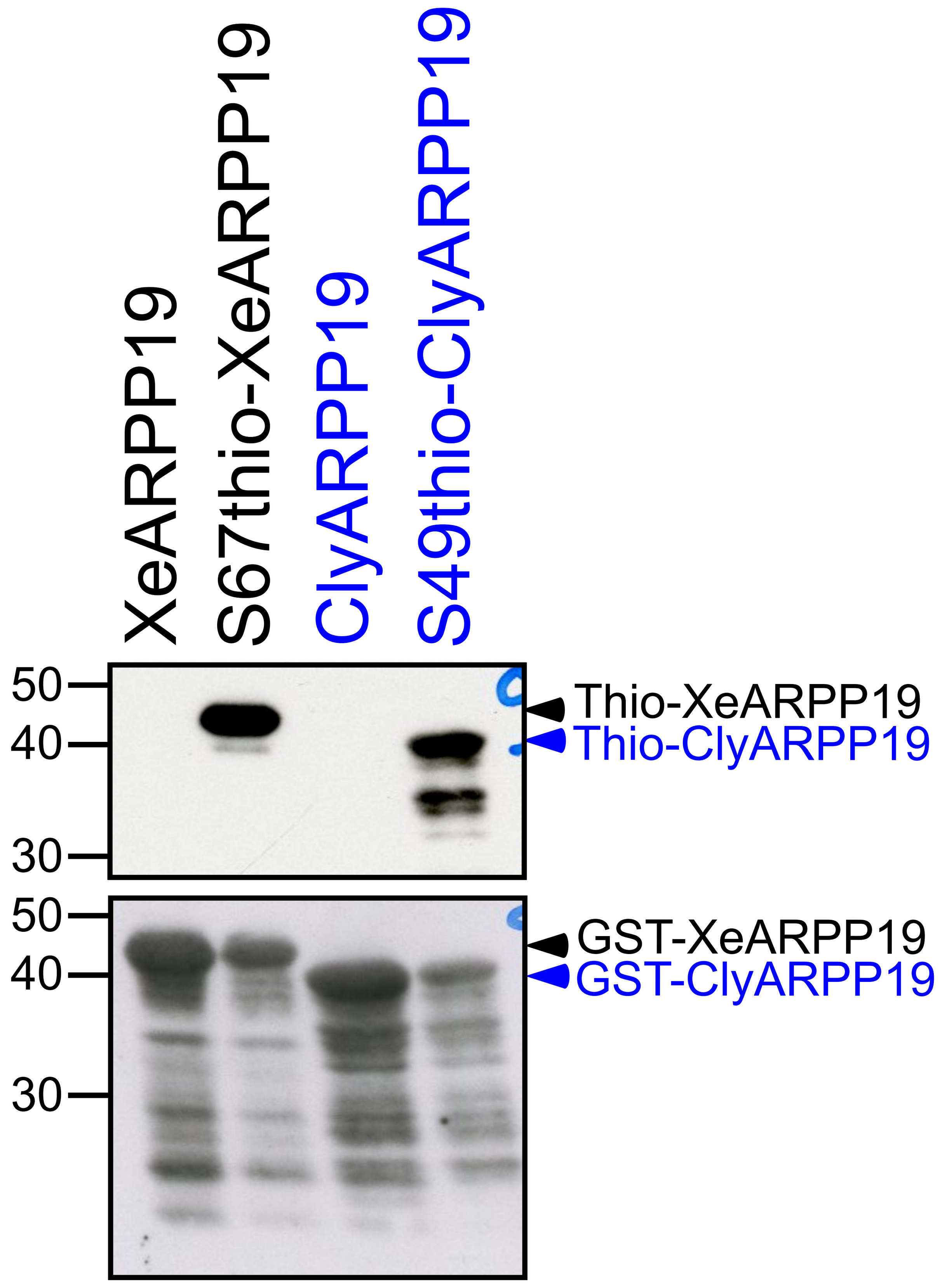

400 μM GST-XeARPP19 or GST-ClyARPP19 were incubated (S67thio-XeARPP19 and S49thio-ClyARPP19) or not (XeARPP19 and ClyARPP19) in the presence of K71M-Gwl bound nickel beads and 1 mM γS-ATP in kinase buffer (20 mM HEPES pH 7.4, 2 mM 2-Mercaptoethanol) in a final volume of 60 μL for 60 min at 30°C under stirring (750 rpm, Thermomixer, Eppendorf). γS-ATP was removed by dialyzing the reaction mix against kinase buffer. After addition of Laemmli buffer[1] and heating for 1 minute at 90°C, the samples were incubated with 1 mM P-nitrobenzyl mesylate (PNBM) (Cayman Chemical) for 1 hour at room temperature. The level of alkylated thiophosphates incorporated by Gwl in ClyARPP19 and XeARPP19 was analyzed by western blot using an antibody directed against thiophosphate ester (1:60,000, Abcam ab92570). Total ClyARPP19 and XeARPP19 were detected by an anti-GST antibody (1:5,000, Sigma A-7340). Note that XeARPP19 and ClyARPP19 exhibit different electrophoretic migration due to their slightly different molecular weights.

[1]Laemmli UK. Cleavage of structural proteins during the assembly of the head of bacteriophage T4. Nature 227, 680-685 (1970).

### Supplementary Fig. S4
### Identification of the serine phosphorylated by PKA in ClyARPP19

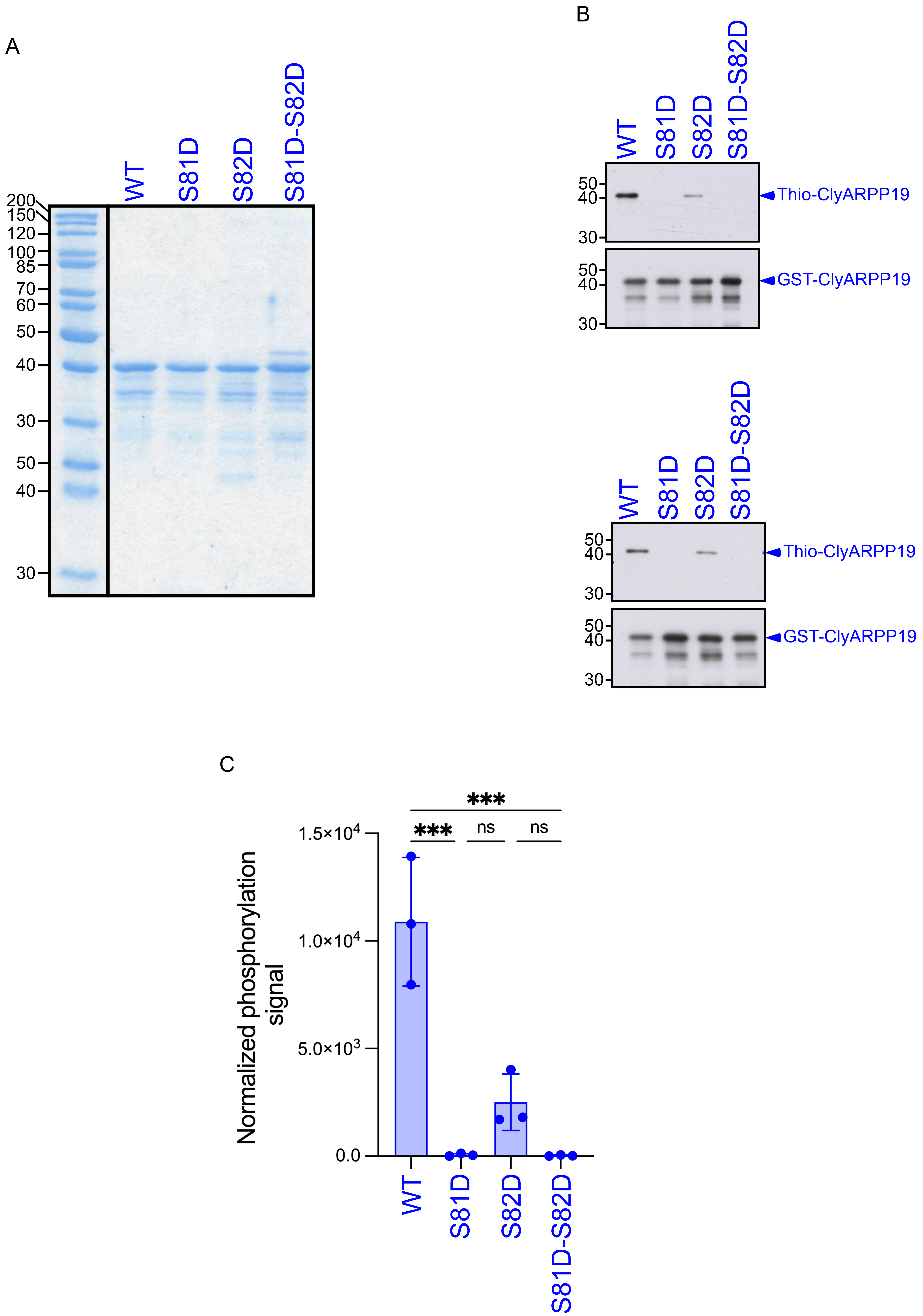

**(A)** Wild-type (WT) and three non-phosphorylable mutants of ClyARPP19 (S81D, S82D, S81D-S82D) were produced to identify the serine phosphorylated by PKA. The purified proteins were loaded and electrophoresed on an acrylamide-SDS gel that was stained by Coomassie blue. **(B)** WT and ClyARPP19 mutants were in vitro thiophosphorylated by recombinant bovine PKA catalytic subunit in the presence of 1mM γS-ATP for 2 hours. The thiophosphorylation was analyzed by western-blot with an anti-thiophosphate ester antibody (1:60,000, Abcam ab92570). The levels of ClyARPP19 proteins were detected with an anti-GST antibody (1:5,000, Sigma A-7340). Two of the three replicates performed are presented here, the third one is presented in Fig. 4. **(C)** The thiophosphorylation level of the four ClyARPP19 forms was normalized by the total quantity of ARPP19 (GST signal). Quantification and statistical analysis of the in vitro kinase assay of the three replicates are represented. Ordinary one-way ANOVA: WT/S81D Pvalue=0.0001, WT/S81D-S82D Pvalue<0.0001, S82D/S81D Pvalue=0.3524 and S82D/S81D-S82D Pvalue=0.3420. Each dot represents an independent experiment.

### Supplementary Fig. S5
### Statistical analysis of GVBD time-courses of oocytes injected with ClyARPP19 and XeARPP19 phosphorylated by PKA

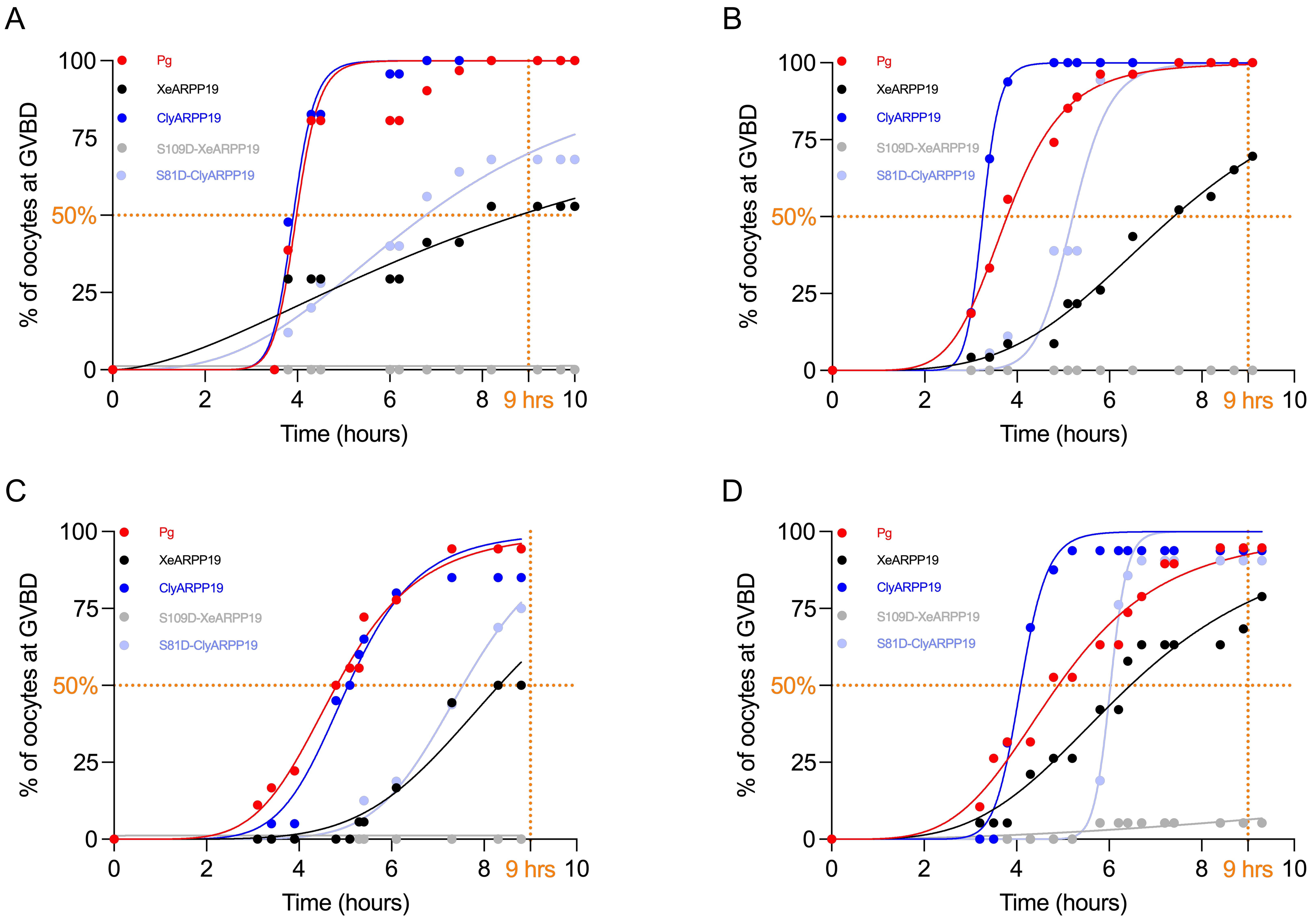

E

| Percentage of GVBD after 9 hours | | | | | |
|---|---|---|---|---|---|
| | Progesterone | XeARPP19 | ClyARPP19 | S109D-XeARPP19 | S81D-ClyARPP19 |
| EXP N°1 | 100 | 52,9 | 100 | 0 | 68 |
| EXP N°2 | 100 | 69,6 | 100 | 0 | 100 |
| EXP N°3 | 94,4 | 50 | 85 | 0 | 75 |
| EXP N°4 | 94,7 | 78,9 | 93,8 | 5,3 | 90,5 |
| Average | 97,3 | 62,9 | 94,7 | 1,3 | 83,4 |
| SD | 3,1 | 13,8 | 7,1 | 2,7 | 14,5 |

F

| Time at GVBD 50% (hours) | | | | | |
|---|---|---|---|---|---|
| | Progesterone | XeARPP19 | ClyARPP19 | S109D-XeARPP19 | S81D-ClyARPP19 |
| EXP N°1 | 3,980 | 8,799 | 3,912 | NA | 6,774 |
| EXP N°2 | 3,783 | 7,391 | 3,254 | NA | 5,207 |
| EXP N°3 | 4,818 | 8,325 | 5,063 | NA | 7,541 |
| EXP N°4 | 4,9 | 6,458 | 4,089 | NA | 6,029 |
| Average | 4,369 | 7,743 | 4,080 | NA | 6,388 |
| SD | 0,569 | 1,037 | 0,748 | NA | 1,000 |

**(A-D)** Oocytes were injected or not (red) with 800 ng/oocyte of either XeARPP19 (black), or ClyARPP19 (dark blue), or S109D-XeARPP19 (grey) or S81D-ClyARPP19 (light blue) and then stimulated by progesterone (Pg). GVBD time-course was recorded. Four independent experiments are illustrated. For each experiment, curves were subjected to a non-linear regression (sigmoid) with the following constrains: bottom=0%, top=100%. The time at 50% of GVBD and the percentage of GVBD after 9 hours are indicated by orange dashed lines and reported respectively in **(E)** and **(F)**. SD: standard deviation.

### Supplementary Fig. S6

### Specificity of the phosphoS109-XeARPP19 and phosphoS81-ClyARPP19 antibodies

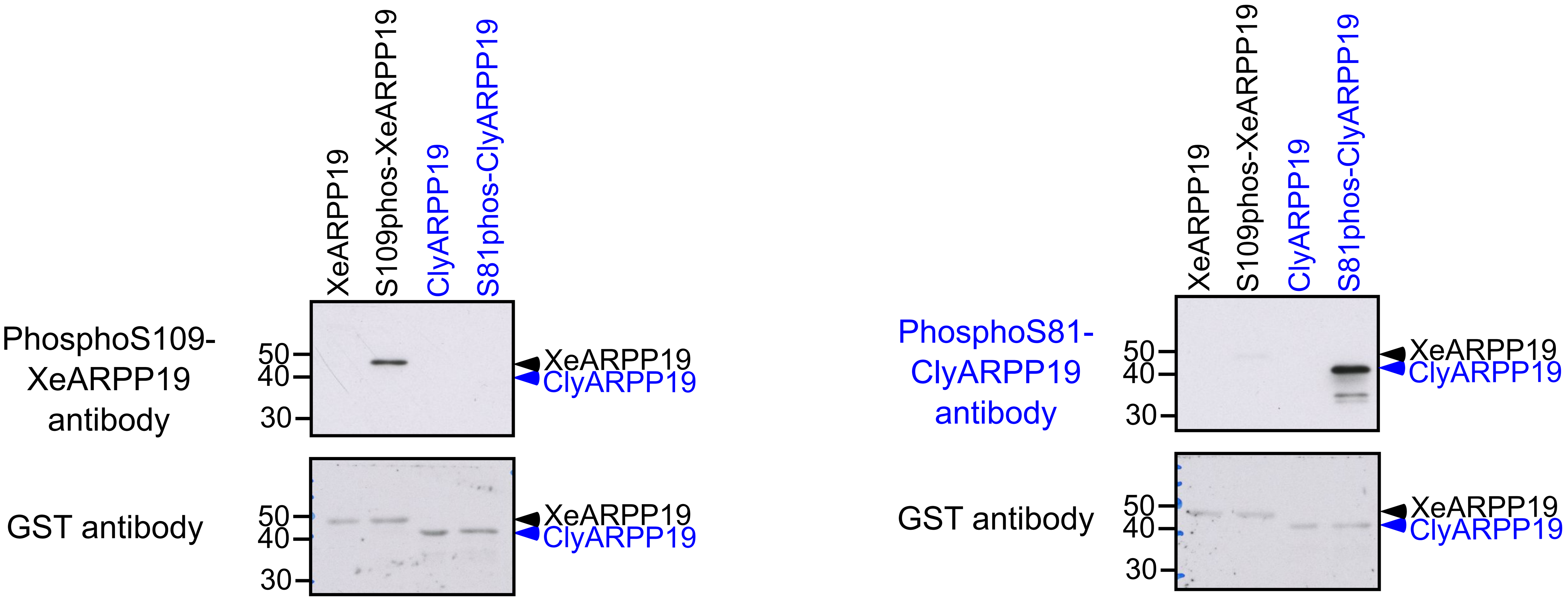

XeARPP19 and ClyARPP19 were in vitro thiophosphorylated or not by recombinant bovine PKA catalytic subunit in the presence of 1mM γS-ATP for 2 hours. The phosphorylation was analyzed by western blot with antibodies raised against phosphoS109-XeARPP19[1] (left panel) or phosphoS81-ClyARPP19 (Covalab) (right panel). The total ARPP19 amount was detected by an anti-GST antibody (1:5,000, Sigma A-7340). Note that XeARPP19 and ClyARPP19 exhibit different electrophoretic migration due to their slightly different molecular weights.

[1]Dupre A, Daldello EM, Nairn AC, Jessus C, Haccard O. Phosphorylation of ARPP19 by protein kinase A prevents meiosis resumption in Xenopus oocytes. Nature communications 5, 3318 (2014).

### Supplementary Fig. S7
### Calibration of phosphorylated S109-XeARPP19 and S81-ClyARPP19 amounts for dephosphorylation analysis

A

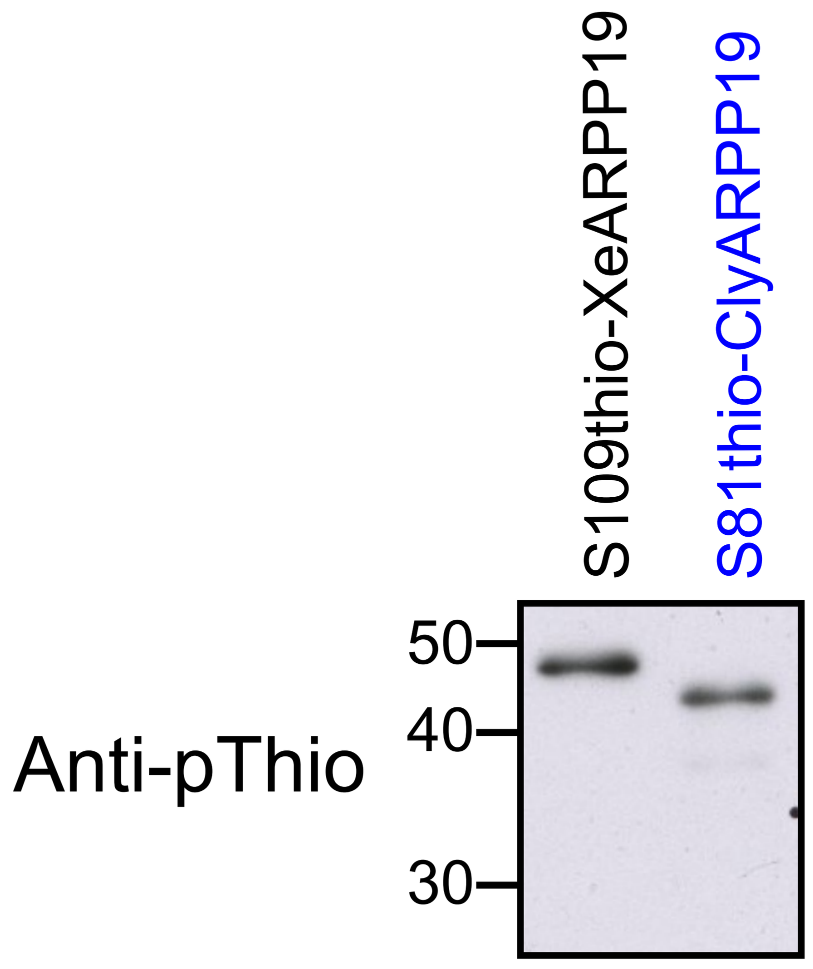

| | signal | Percentage conversion with S109thio-XeARPP19 setted as 100% |
|---|---|---|
| S109thio-XeARPP19 | 11656,912 | 100 |
| S81thio-XeARPP19 | 7981,548 | 68 |

B

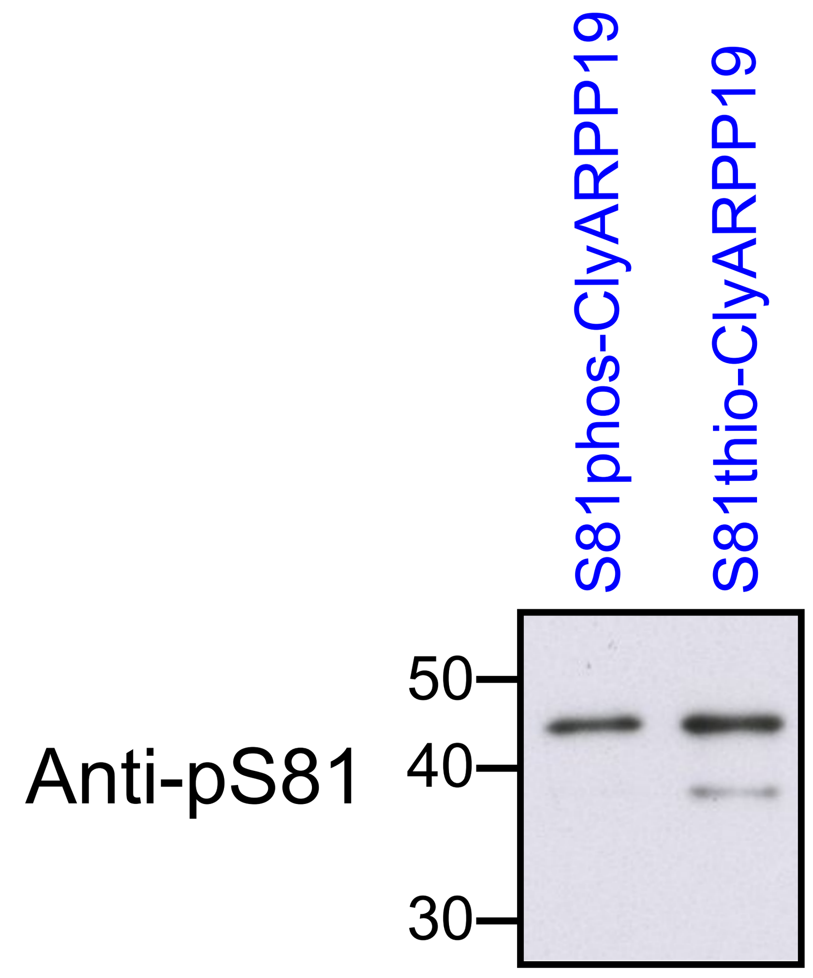

| | | |
|---|---|---|
| S81thio-ClyARPP19 | 11801,548 | 68 |
| S81phos-ClyARPP19 | 8442,012 | 49 |

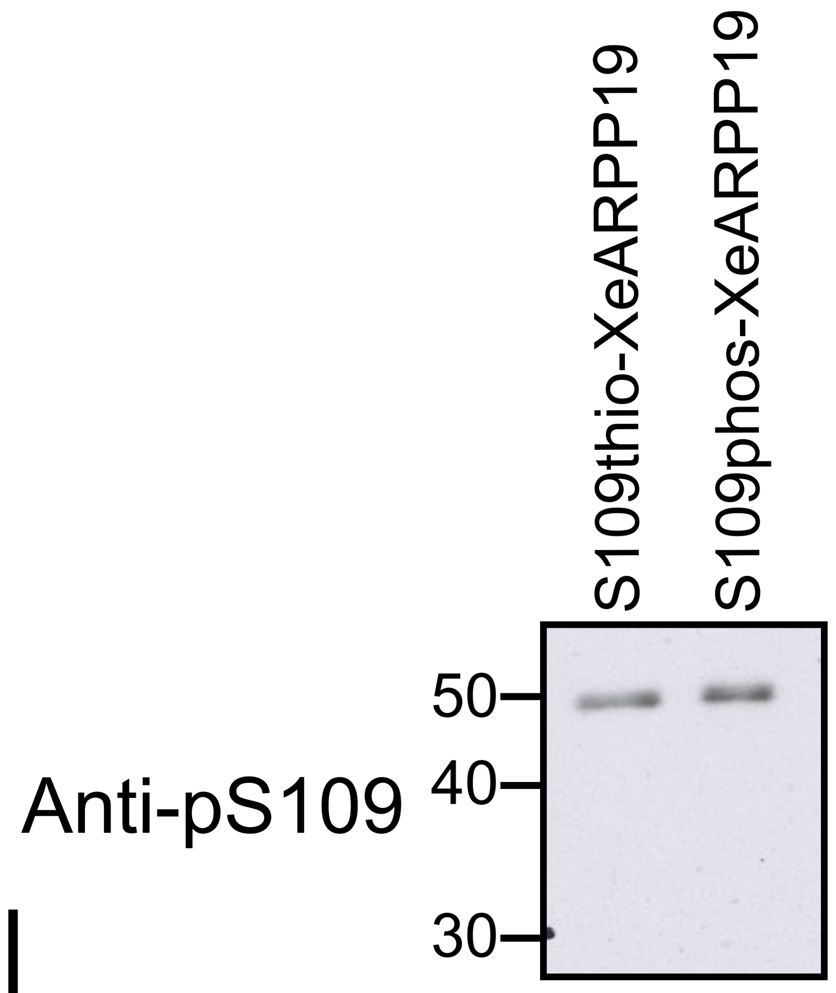

| | | |
|---|---|---|
| S109thio-XeARPP19 | 7868,477 | 100 |
| S109phos-XeARPP19 | 8610,355 | 109 |

C

| | Percentage |
|---|---|
| S81phos-ClyARPP19 | 49 |
| S109phos-XeARPP19 | 109 |

x 2,2

To start the dephosphorylation assays at time 0 **(Fig. 6)** with an identical amount of phosphorylated S109-XeARPP19 and S81-ClyARPP19 proteins, XeARPP19 and ClyARPP19 were in vitro incubated with recombinant bovine PKA catalytic subunit for 2 hours, either with 1mM γS-ATP (thiophosphorylation) or 200 µM ATP (phosphorylation). **(A)** Thiophosphorylated S109-XeARPP19 and S81-ClyARPP19 were analysed by western blot with a same antibody for both proteins, the anti-thiophosphate ester antibody, anti-pThio (1:60,000, Abcam ab92570). The signal intensities obtained with this antibody were quantified using Image J software and converted in % of phosphorylation, 100% being affected to the highest signal. **(B)** Then, thiophosphorylated and phosphorylated proteins were western blotted using antibodies directed against the phosphorylated proteins, the anti-phoshoS81, anti-pS81, for ClyARPP19 (Covalab, Supp Fig. S6) and the anti-phoshoS109, anti-pS109, for XeARPP19[1]. The respective levels of phosphorylated and thiophosphorylated proteins of each species were quantified using Image J software. Based on the % of thiophosphorylation determined in **(A)**, the signals obtained for the phosphorylated forms were converted as a %. **(C)** The phosphorylation level of XeARPP19 and S81-ClyARPP19 differs by 2.2-fold. This ratio was taken in account for the dephosphorylation assay.

[1]Dupre A, Daldello EM, Nairn AC, Jessus C, Haccard O. Phosphorylation of ARPP19 by protein kinase A prevents meiosis resumption in Xenopus oocytes. Nature communications 5, 3318 (2014).

### Supp Fig. S8
### Statistical analysis of XeARPP19 and ClyARPP19 dephosphorylation by PP2A-B55δ

A

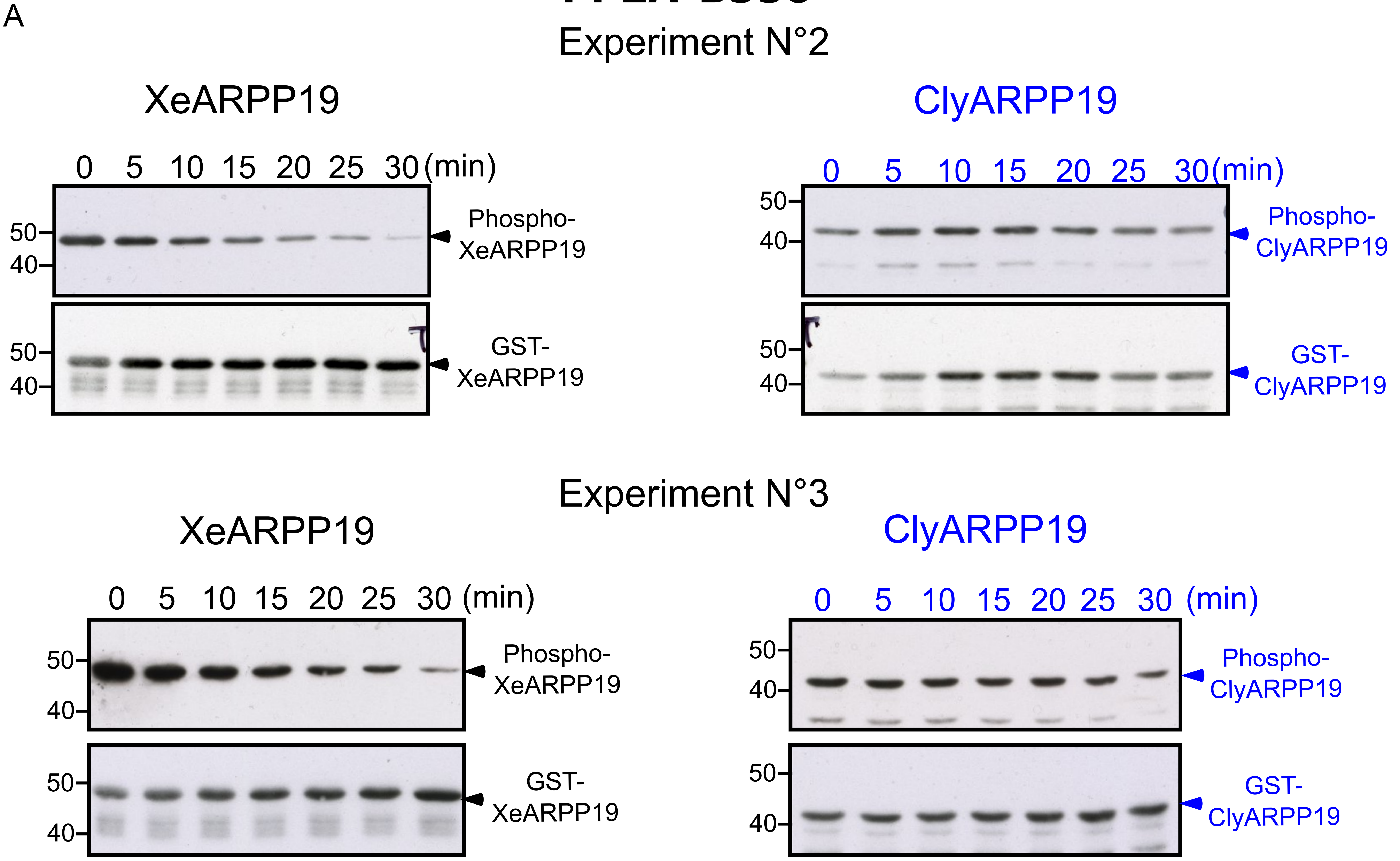

B

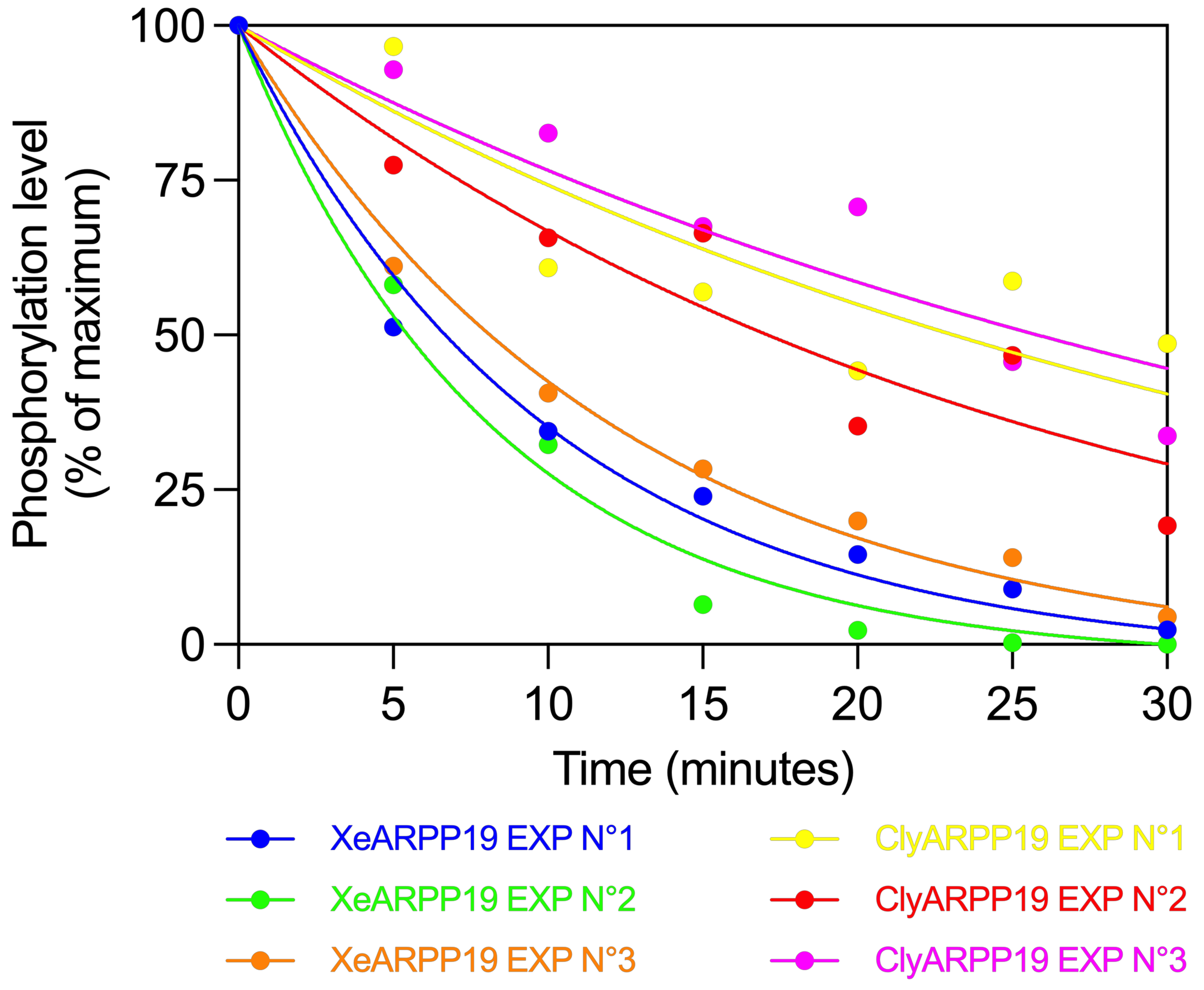

C

| Dephosphorylation half-time (minutes) | | | | | |
|---|---|---|---|---|---|
| | EXP N°1 | EXP N°2 | EXP N°3 | Average | SD |
| XeARPP19 | 6,88 | 5,64 | 8,38 | 6,97 | 1,37 |
| ClyARPP19 | 29,89 | 22,38 | 32,73 | 28,33 | 5,35 |

**(A)** Xenopus prophase oocyte extracts were incubated with hexokinase/glucose and PKI. They were supplemented with S109-phosphorylated XeARPP19 (left panels) or S81-phosphorylated ClyARPP19 (right panels). At indicated times, samples were collected and the phosphorylation level was followed by western blot using antibodies directed against phosphoS109-XeARPP19[1] or phosphoS81-ClyARPP19 (Covalab, Supp Fig. S6). The total ARPP19 amount was detected by an anti-GST antibody (1:5,000, Sigma A-7340). Two independent experiments are presented. A third identical experiment is illustrated in Fig. 6B-D. **(B)** The levels of phosphorylated proteins of the three replicates in **(A)** and Fig. 6B were quantified by Image J software. For each replicate, the percentage of phosphorylation was plotted and a non-linear regression (exponential: One-phase decay) was applied (constraints: Y0=0%, plateau=same for each replicate). **(C)** Dephosphorylation half-time of XeARPP19 and ClyARPP19, averages and standards deviations (SD).

[1]Dupre A, Daldello EM, Nairn AC, Jessus C, Haccard O. Phosphorylation of ARPP19 by protein kinase A prevents meiosis resumption in Xenopus oocytes. Nature communications 5, 3318 (2014).

### Supplementary Fig. S9
### Specificity of the antibody directed against the B55 subunit of PP2A

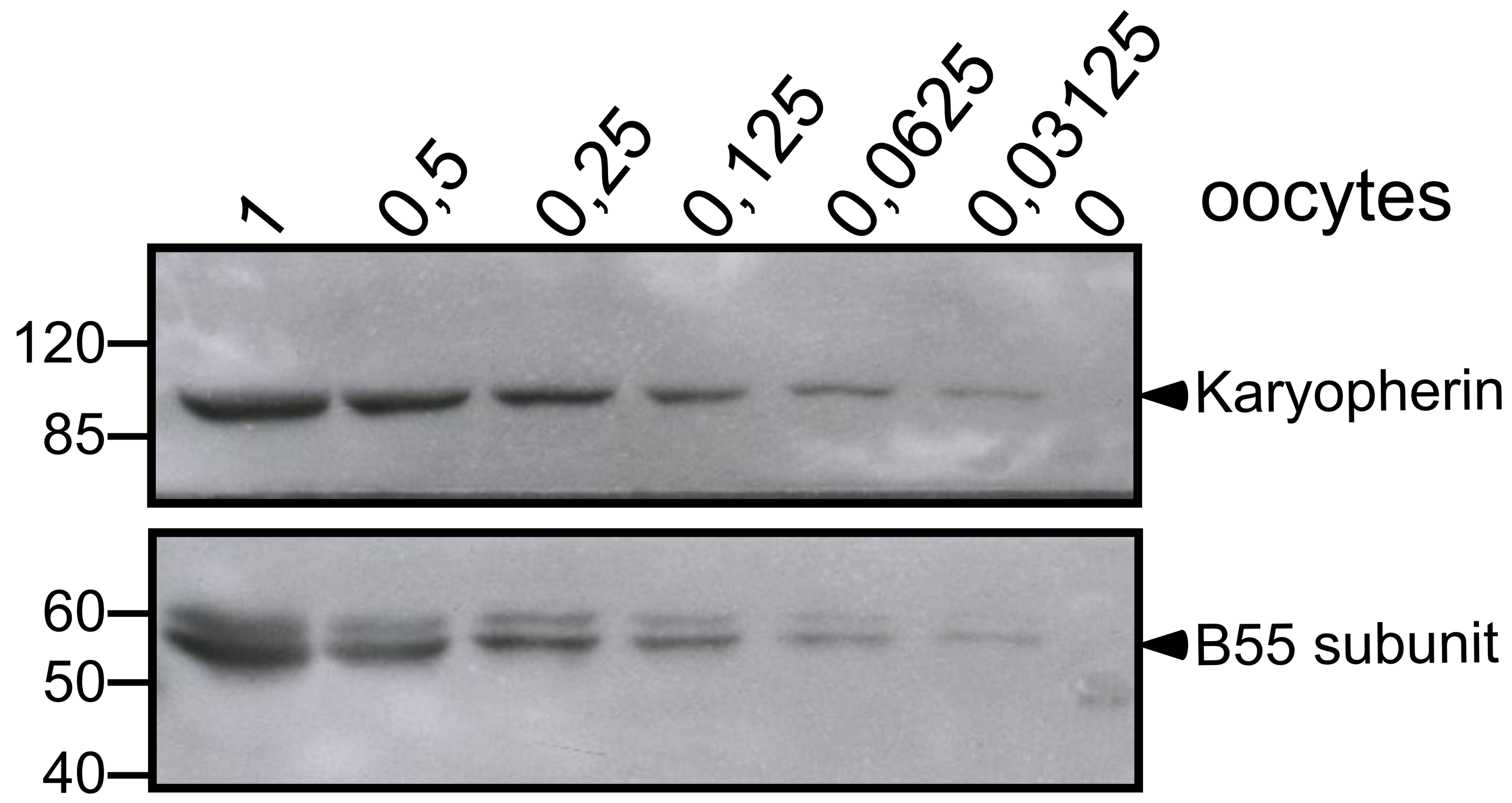

Xenopus prophase-arrested oocyte extracts were gradually diluted as indicated by the equivalent amounts of oocytes loaded per lane and then submitted to western blot with a polyclonal rabbit antibody directed against Xenopus B55δ (Covalab). Karyopherin was used as an internal control (anti-karyopherin antibody: 1:1,000, Santa-Cruz biotechnology sc-1863).

### Supplementary Fig. S10
### Statistical analysis of XeARPP19 and ClyARPP19 phosphorylation by recombinant PKA

A

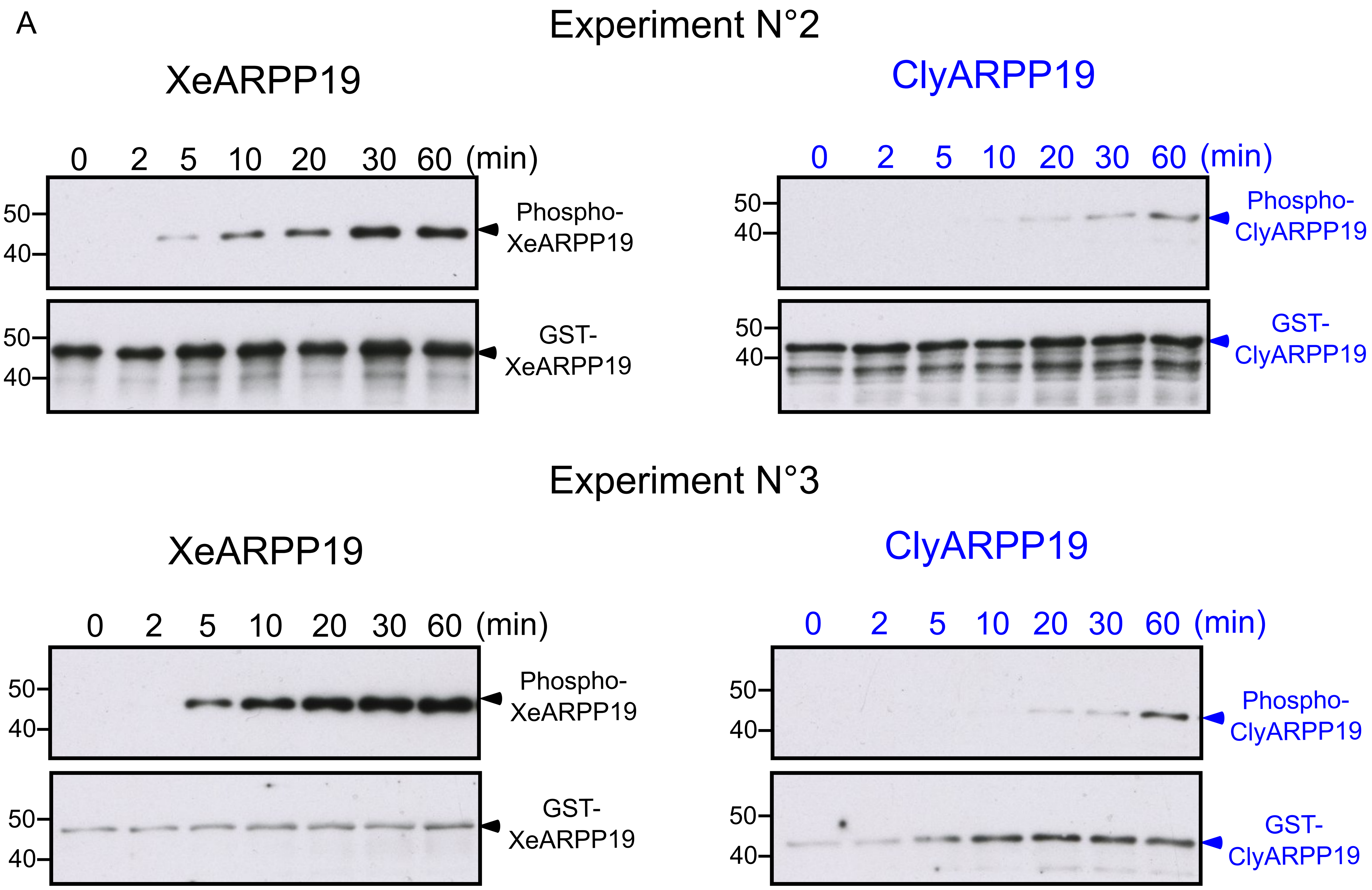

B

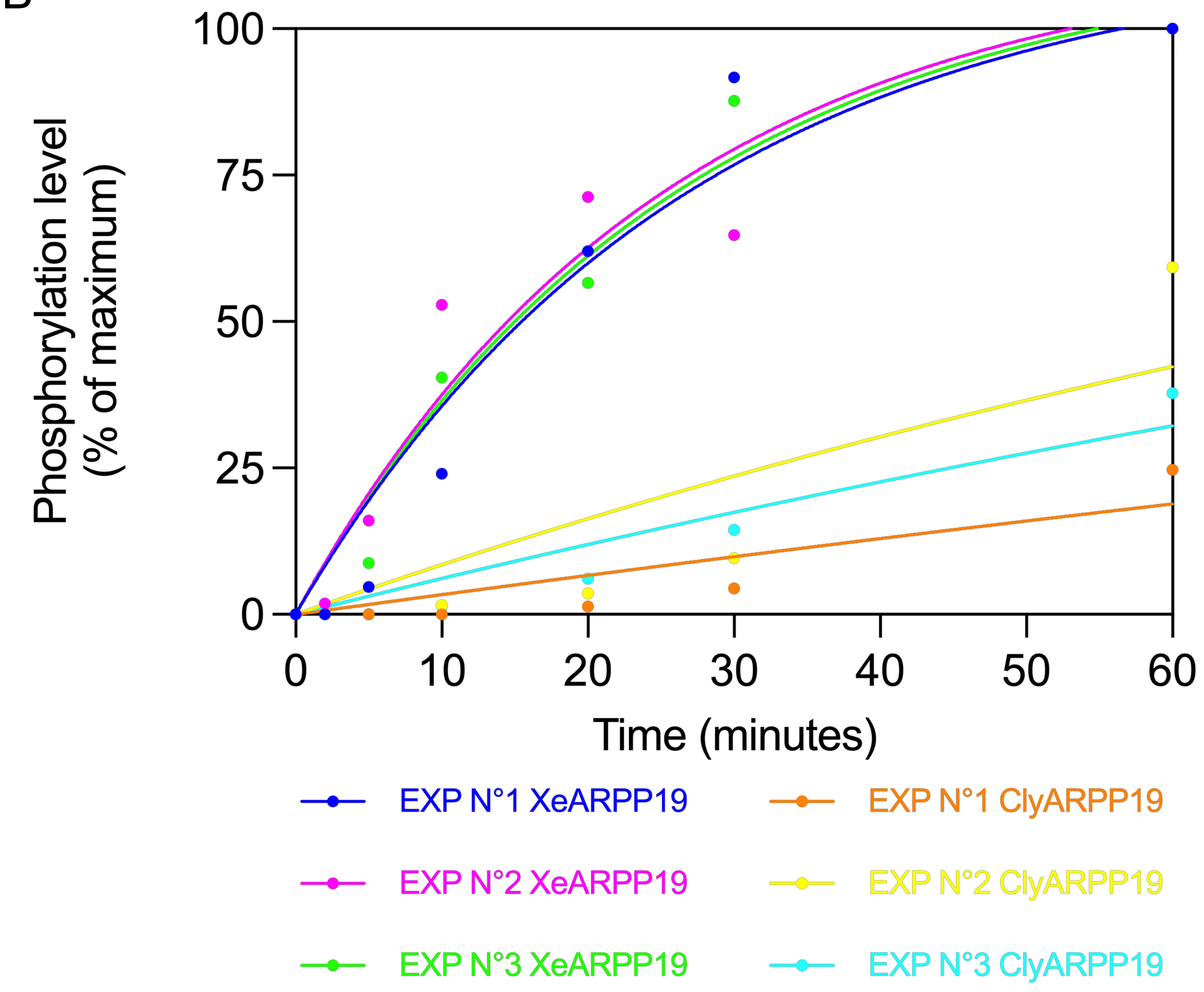

C

| Phosphorylation half-time (minutes) | | | | | |
|---|---|---|---|---|---|
| | EXP N°1 | EXP N°2 | EXP N°3 | Average | SD |
| XeARPP19 | 18,18 | 16,98 | 17,61 | 17,59 | 0,60 |
| ClyARPP19 | 229,1 | 89,33 | 124,9 | 147,78 | 72,64 |

**(A)** XeARPP19 (left panels) or ClyARPP19 (right panels) were incubated with purified bovine PKA in the presence of 1mM γS-ATP. Samples were collected at various times as indicated. The phosphorylation of ARPP19 proteins was assessed by western blot with an anti-thiophosphate ester antibody (1:60,000, Abcam ab92570). The total ARPP19 amount was detected by an anti-GST antibody (1:5,000, Sigma A-7340). Two independent experiments are presented. A third identical experiment is illustrated in Fig. 8. **(B)** The levels of phosphorylated proteins of the three replicates in **(A)** and Fig. 9 were quantified by Image J software. For each replicate, the percentage of phosphorylation was plotted and a non-linear regression (exponential: One-phase association) was applied (constraints: Y0=100%, plateau=same for each replicate). **(C)** Phosphorylation half-time of XeARPP19 and ClyARPP19, averages and standards deviations (SD).

### Supplementary Fig. S11
### Replicates of the in vivo PKA phosphorylation experiments of XeARPP19 and ClyARPP19

#### Experiment N°2

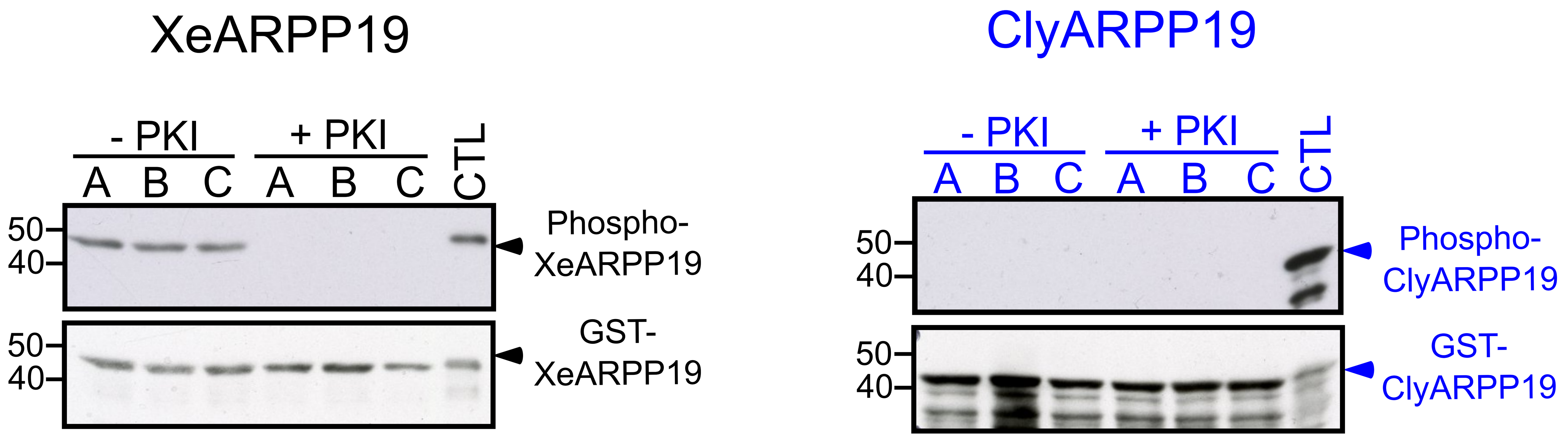

#### Experiment N°3

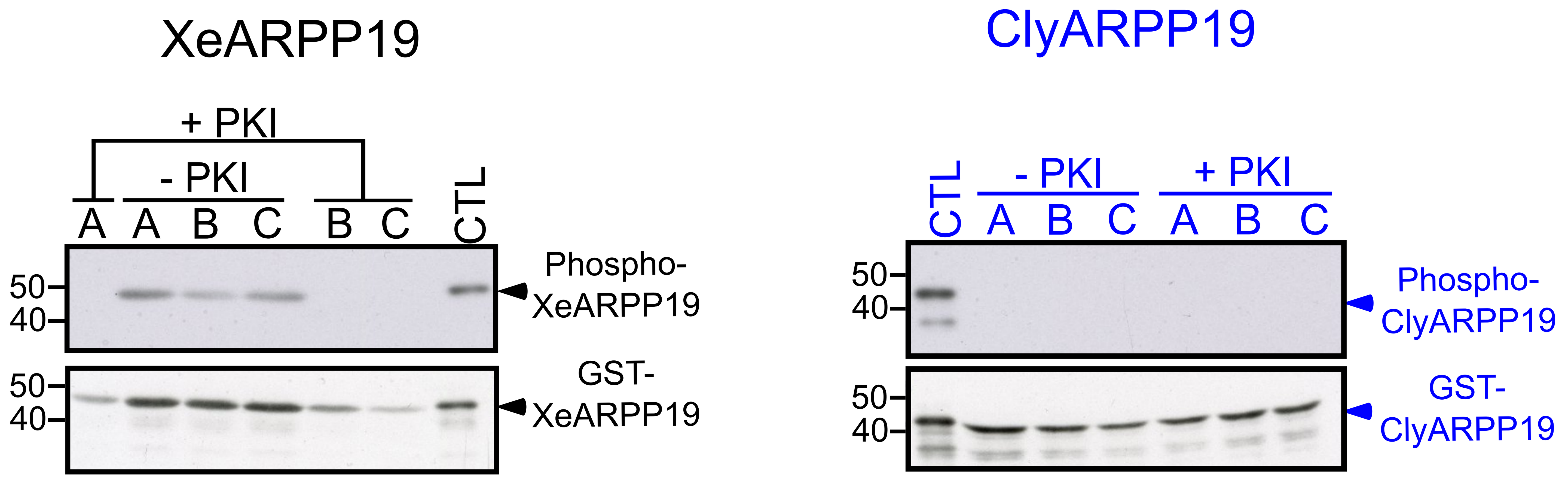

XeARPP19 or ClyARPP19 were injected into Xenopus oocytes previously injected or not with PKI. 30 min after ARPP19 injection, injected ARPP19 proteins were recovered by GST pull-down and their phosphorylation was monitored by western blot using antibodies against phosphoS109-XeARPP19[1] or phosphoS81-ClyARPP19 (Covalab, Supp Fig. S6). The total ARPP19 amount was detected with an anti-GST antibody (1:5,000, Sigma A-7340). Three groups of 5 oocytes (A, B and C) were analyzed in both conditions. An in vitro phosphorylated form of ARPP19 was used as a positive control of phosphorylation (CTL). Two replicates from 3 independent experiments are shown. The third experiment is illustrated in **Fig. 9A**.

[1]Dupre A, Daldello EM, Nairn AC, Jessus C, Haccard O. Phosphorylation of ARPP19 by protein kinase A prevents meiosis resumption in Xenopus oocytes. Nature communications 5, 3318 (2014).

### Supplementary Fig. S12
### Sequence alignment of Xenopus, Clytia and bovine PKA catalytic subunit

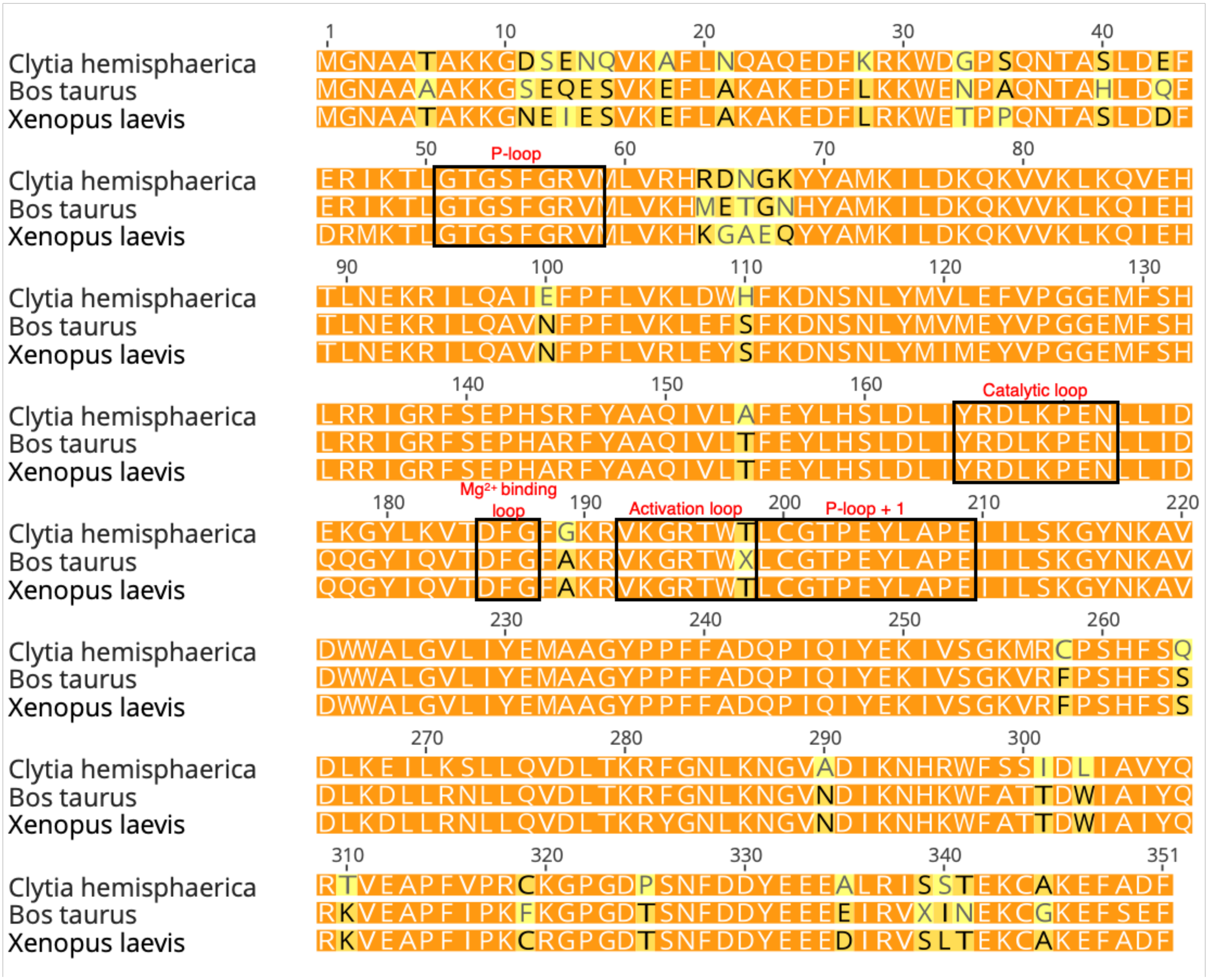

Sequence alignment of the sequences of the α-catalytic subunit of PKA of Clytia hemisphaerica, Xenopus laevis and Bos taurus, obtained with Geneious 2023.0 (Dotmatic). Colors represent the % of identity: dark orange: 100% identity; light orange: 100-80% identity; yellow: 80-60% identity; white: below 60% identity. The catalytic loop, the activation loop, the Mg2+-binding loop and the two P-loops are framed.

**Supplementary Table S1. Detailed data of injection experiments in *Clytia* oocytes.**

The following recombinant GST-fused proteins diluted in PBS were injected into *Clytia* prophase-arrested oocytes: wild-type (WT) ClyARPP19, wild-type (WT) XeARPP19, S49A-mutated ClyARPP19, S67A-mutated XeARPP19, SS81-82DD-double mutated ClyARPP19, S109D-mutated XeARPP19, Gwl-thiophosphorylated ClyARPP19 (GwlThio-ClyARPP19), Gwl-thiophosphorylated XeARPP19 (GwlThio-XeARPP19), PKA-thiophosphorylated ClyARPP19 (PKAThio-ClyARPP19) and PKA-thiophosphorylated XeARPP19 (PKAThio-XeARPP19). Okadaic acid (OA) or control buffer (PBS at various dilutions) were also injected.

Within one to two hours after injection, some oocytes underwent GVBD ($4^{th}$ column). In some experiments, oocytes were observed closely or filmed during this period and the occurrence of strong contractions was noted, in some cases resulting in lysis ($5^{th}$ column). In experiments where oocytes were not continuously observed during this period, only lysis was scored and any earlier contractions were not recorded ($6^{th}$ column). In experiments where maturation (scored as GVBD) was induced using MIH, any oocytes showing previous GVBD or contractions/lysis were excluded. MIH treatment was $10^{-7}$ M WPRP-amide in all experiments, except one in which $10^{-8}$ M WPRP-amide was used and led to 100% maturation.

Not monitored: oocytes not continuously observed, any earlier contractions were not recorded although they may have occurred.

### Some non-injected oocytes also lysed in this experiment.

| | | | Phenotypes before MIH | | | Phenotypes after MIH | |
|---|---|---|---|---|---|---|---|
| **Injected solution** | **Date** | **Concentration** | **GVBD** | **Contractions** (in some cases followed by lysis) | **Lysis** | **GVBD** | **Other phenotypes** |
| WT ClyARPP19 | 07-08-15 | 2mg/ml | 0/21 | Not monitored | 2/21 | 19/19 | - |
| WT ClyARPP19 | 21-05-15 | 2mg/ml | 3/49 | Not monitored | 4/49 | 45/49 | 4 lysis |
| WT ClyARPP19 | 04-10-22 | 4mg/ml | Not monitored | Not monitored | 5/10 | 5/5 | - |
| WT ClyARPP19 | 05-10-22 | 3mg/ml | 2/30 | Not monitored | 8/30 | 20/20 | - |
| WT ClyARPP19 | 24-01-23 | 3mg/ml | 0/30 | Not monitored | 2/30 | 18/18 | - |
| WT ClyARPP19 | 27-01-22 | 4mg/ml | 2/40 | Not monitored | 14/40 | 1/1 | - |
| WT XeARPP19 | 30-06-15 | 2mg/ml | 1/31 | Not monitored | 3/31 | 27/27 | |
| WT XeARPP19 | 22-09-22 | 4mg/ml | 0/20 | Not monitored | 20/20 | | |
| WT XeARPP19 | 27-09-22 | 4mg/ml | | | | 26/26 | |
| WT XeARPP19 | 27-09-22 | 4mg/ml | | | | 5/5 | - |
| WT XeARPP19 | 05-10-22 | 3mg/ml | 0/32 | Not monitored | 17/32 | 15/15 | - |
| WT XeARPP19 | 27-01-22 | 4mg/ml | 2/40 after 1h | Not monitored | 5/40 | 6/6 | - |
| WT XeARPP19 | 27-01-22 | 3mg/ml | 1/40 after 1 h | Not monitored | 6/40 | 1/1 | - |
| WT XeARPP19 | 31-01-22 | 4mg/ml | 0 | | 0 | 10/10 | 1 contraction; 1 lysis; 1 multiple pb |

| | | | | | | | |
|---|---|---|---|---|---|---|---|
| S49A-ClyARPP19 | 21-05-15 | 2mg/ml | 0/25 | | 0/25 | 26/26 | - |
| S49A-ClyARPP19 | 05-10-22 | 3mg/ml | 1/30 | Not monitored | 8/30 | 17/17 | - |
| S49A-ClyARPP19 | 14-10-22 | 2mg/ml | 8/43 | Not monitored | 4/43 | 23/24 | - |
| S67A-XeARPP19 | 30-06-15 | 2mg/ml | 0/14 | Not monitored | 2/14 | 17/21 | 4 lysis |
| S67A-XeARPP19 | 11-10-22 | 3mg/ml | | Not monitored | 21/30 | 18/18 | - |
| S67A-XeARPP19 | 11-10-22b | 3mg/ml | 0/23 | | 8/23 | | - |
| S67A-XeARPP19 | 13-10-22 | 3mg/ml | 5/21 | Not monitored | 9/21 | 7/7 | - |
| S67A-XeARPP19 | 14-10-22 | 2mg/ml | 10/26 | Not monitored | 2/26 | 22/23 | - |
| 81-82DD-ClyARPP19 | 22-05-15 | 2mg/ml | 0/18 | Not monitored | 1/18 | 27/28 | - |
| 81-82DD-ClyARPP19 | 05-10-22 | 3mg/ml | 0/30 | Not monitored | 13/30 | 17/17 | - |
| 81-82DD-ClyARPP19 | 11-10-22 | 3mg/ml | | Not monitored | 30/30 | | |
| 81-82DD-ClyARPP19 | 11-10-22b | 3mg/ml | | Not monitored | 20/23 | 19/20 | - |
| 81-82DD-ClyARPP19 | 14-10-22 | 2mg/ml | 12/36 | Not monitored | 2/36 | 22/22 | - |
| S109D-XeARPP19 | 19-05-15 | 2mg/ml | 0/25 | | 2/25 | 23/25 | |
| S109D-XeARPP19 | 22-09-22 | 4mg/ml | 0/20 | Not monitored | 20/20 | | |
| S109D-XeARPP19 | 27-09-22 | 4mg/ml | | | | 14/15 | 3 lysis; 1 contraction/fission |
| S109D-XeARPP19 | 14-10-22 | 2mg/ml | 9/38 | Not monitored | 4/38 | 23/23 | - |
| GwlThio-ClyARPP19 | 28-03-23 | 4mg/ml | 0/9 | 2/9 | 1/9 | | |
| GwlThio-ClyARPP19 | 28-03-23b | 4mg/ml | 0/13 | 3/13 | 1/13 | | |
| GwlThio-ClyARPP19 | 04-04-23 | 4mg/ml | 0/13 | 1/13 | 0/13 | | |
| GwlThio-ClyARPP19 | 04-04-23b | 4mg/ml | 0/13 | 1/13 | 0/13 | 10/11 | 1 exploded |
| GwlThio-XeARPP19 | 21-03-23 | 4mg/ml | 0/11 | 1/11 | 2/11 | 6/7 | 1 lysis |
| GwlThio-XeARPP19 | 23-03-23 | 4mg/ml | 1/20 | 2/20 | 1/20 | 6/6 | some minor contractions |
| GwlThio-XeARPP19 | 28-03-23 | 4mg/ml | 1/12 | 2/9 | | - | |
| GwlThio-XeARPP19 | 07-04-23 | 4mg/ml | 0/12 | 1 or 2/12 | | | |
| GwlThio-XeARPP19 | 07-04-23b | 4mg/ml | 0/12 | 3/12 | 4/12# | | |
| PKAThio-ClyARPP19 | 29-04-22 | 2mg/ml | 3/34 | Not monitored | 0/34 | 31/31 | - |
| PKAThio-ClyARPP19 | 03-05-22 | 4mg/ml | 6/32 | Not monitored | 3/32 | 23/23 | 0/23 |
| PKAThio-XeARPP19 | 29-04-22 | 2mg/ml | 6/27 | Not monitored | 0/27 | 21/21 | - |
| PKAThio-XeARPP19 | 03-05-22 | 4mg/ml | 0/30 | Not monitored | 0/32 | 30/32 | 0/32 |
| PBS | 21-05-15 | Undiluted | 1/13 | Not monitored | 1/13 | 12/12 | - |

| | | | | | | | |
|---|---|---|---|---|---|---|---|
| PBS | 22-05-15 | Undiluted | 0/14 | | 0/14 | 23/24 | |
| PBS | 30-06-15 | Undiluted | 0 | 0 | 0 | 100% | - |
| PBS | 29-04-22 | Undiluted | 0/28 | Not monitored | 0/28 | 28/28 | - |
| PBS | 03-05-22 | Undiluted | 1/30 | Not monitored | 1/30 | 28/28 | - |
| PBS | 22-09-22 | Undiluted | Not monitored | Not monitored | 10/20 | | |
| PBS | 27-09-22 | Undiluted | | | - | 16/18 | - |
| PBS | 05-10-22 | Undiluted | 0/30 | Not monitored | 15/30 | 15/15 | |
| PBS | 13-10-22 | 1/3 | 3/23 | Not monitored | 0/23 | 20/20 | |
| PBS | 14-10-22 | 1/5 | 10/46 | Not monitored | 6/46 | 30/30 | |
| PBS | 04-04-23 | Undiluted | 0/12 | | 11/12 | | |
| PBS | 04-04-23 | 1/2 | 0/13 | 0/13 | 0/13 | | |
| Okadaic Acid | 16-05-23 | 2μM in 50% PBS | 0/14 | 14/14 minor contractions then lysis | | | |
